# A Bayesian Study to Optimally Unveil LDH Kinetic Mechanisms *In Vivo*

**DOI:** 10.64898/2026.07.30.741466

**Authors:** David Gomez-Cabeza, Lluis Mangas-Florencio, Gergő Matajsz, Adriana Gonzalez, Irene Marco-Rius

## Abstract

Many life-threatening diseases present characteristic changes in metabolic profiles throughout their progression. For example, an increase in lactic fermentation is characteristic of cancer and is described by the Warburg effect. Yet, only a few methodologies allow for detecting these changes non-invasively and in real-time. Furthermore, the frequent oversight of the system’s biochemical mechanisms and the elevated time and resources required for experimentation risk stagnating research. Here, we address both problems thanks to mathematical modelling and Bayesian statistics. Thanks to modelling, we guided understanding of the lactate dehydrogenase system, highlighting the importance of cellular microenvironment effects, membrane transport and enzymatic repression. To deal with increasing model complexity, we introduced and validated a novel Bayesian optimal experimental design approach to maximise the informative content of experiments. Hence, our approach generated fast and efficient lactate dehydrogenase kinetics characterisation, resulting in generalisable and reliable predictions *in vitro* and *in vivo*.

## 1 Introduction

An increase in lactic fermentation, mediated by the cytoplasmic lactate dehydrogenase (LDH) enzyme, is a pivotal dysregulation biomarker in many human diseases. Glycolysis is a well-studied fundamental process of glucose breakdown in cellular metabolism. The primary end product of glycolysis (i.e., pyruvate) can undergo lactic fermentation under anaerobic conditions in some mammalian cell types [1]. Yet, in certain cases, the reaction also occurs under aerobic conditions, including inflammatory immunological responses and embryonic development [2, 3]. Hence, lactic fermentation of pyruvate is a pathological trait linked to autoimmune diseases, neurodegenerative diseases, and cancer [4, 5]. Particularly, in the context of aerobic cancer metabolism, this excessive pyruvate fermentation, known as the Warburg effect, was a prominent research topic of interest over the past century [6]. However, despite extensive efforts to exploit lactic fermentation as a diagnostic biomarker [7] and a cancer treatment target [8], a lack of consensus remains regarding its underlying mechanisms.

The tetrameric structure of LDH introduces heterogeneity across different tissues, complicating its mechanism studies, which are mostly limited to *ex vivo* and *in vitro* assays. The two core LDH subunits are the muscle (i.e., LDHA) and heart (i.e., LDHB) ones, which are tissue dependent, and their intracellular expression determines the proportion of the five possible combinatorial LDH isoforms [9]. LDH-5 accommodates four LDHA subunits and drives the metabolic reaction from pyruvate to lactate. In comparison, LDH-1 is the isoform with four LDHB subunits, pushing for the backwards reaction, with a gradient of activities for the other three alternatives [10]. To understand the mechanics of LDH, *in vitro* assays can assess its activity and isoform tissue specificity, such as mass spectrometry, colourimetry and spectrophotometry [11–13]. Unfortunately, LDH *in vitro* analyses are unrepresentative of *in vivo* kinetics, which are influenced by the crowded cytoplasmic microenvironment [14]. Besides analysis of fresh biopsies [15], one of the few available techniques to study LDH kinetics in real-time, non-invasively and non-destructively is hyperpolarised nuclear magnetic resonance (HP-NMR) [12, 16–20]. Nevertheless, demonstrated by the inconsistency in both methodology and results across literature, we need a systematic way to study and, hence, utterly understand lactic fermentation.

Mathematical modelling is a powerful analytical tool that can aid in understanding, predicting, controlling and even re-designing biological systems [21]. Hence, to better understand a biological system (e.g., LDH kinetics), we can describe it using mass action kinetics containing specific reaction rates (i.e., parameters). Yet, despite the potential of these models to unlock a profound understanding of biology [22], their general uptake has been generally stagnant or nonexistent. Besides the expertise needed to develop such models, parameters are hard or impractical to measure experimentally, leaving researchers to face complex optimisation problems to infer them [23]. Bayesian statistics is an enticing approach to infer these model parameters, reducing normality assumptions, and introducing accurate uncertainty estimates, highly needed for biological systems [24]. However, due to relying on sampling-based methods, Bayesian strategies are computationally expensive and time-consuming, particularly for complex models. Furthermore, different experiments will carry diverse amounts of information for the task, unknown ahead of performing the inference [25]. To aid with these issues, optimal experimental design (OED) presents an enticing approach focused on increasing efficiency [26]. Yet, most OED strategies use frequentist statistics based on the Fisher information matrix [27], and the few Bayesian ones rely on the usually computationally infeasible mutual information [28, 29]. Furthermore, in biological contexts, OED has been limited to computational studies [30], without substantial practical implementations and validations *in vivo*.

Here we present a systematic study to thoroughly investigate the LDH kinetics (both *in vivo* and *ex vivo*) using mathematical modelling and Bayesian statistics to fully unravel and understand enzymatic repression dynamics caused by the order of insertion of substrate and cofactors in the catalytic pocket of the enzyme (Fig. 1a-b). For the *in vivo* experimentation, we used HP-NMR to acquire, in real-time and non-invasively, the intracellular LDH dynamics of HepG2 (hepatocellular carcinoma) cells (Fig. 1c-e). Moreover, to overcome the high economic, resource and time-consuming HP-NMR experiments, we introduced OED to design highly informative experiments, presenting a novel approach based on maximising prior prediction uncertainty (Fig. 1f). Furthermore, we validated our Bayesian OED (bOED) algorithm *in silico* and, for the first time in a biological context, *in vivo*, demonstrating the numerous advantages of the design approach to aid in accurately inferring model parameters. Thanks to our bOED algorithm, we generated a calibrated LDH model, which we used to increase understanding of the system by predicting different intermediate reaction state dynamics and to elucidate the higher dynamics present in traditional *ex vivo* set-ups to assess LDH activity.

**Fig. 1:**
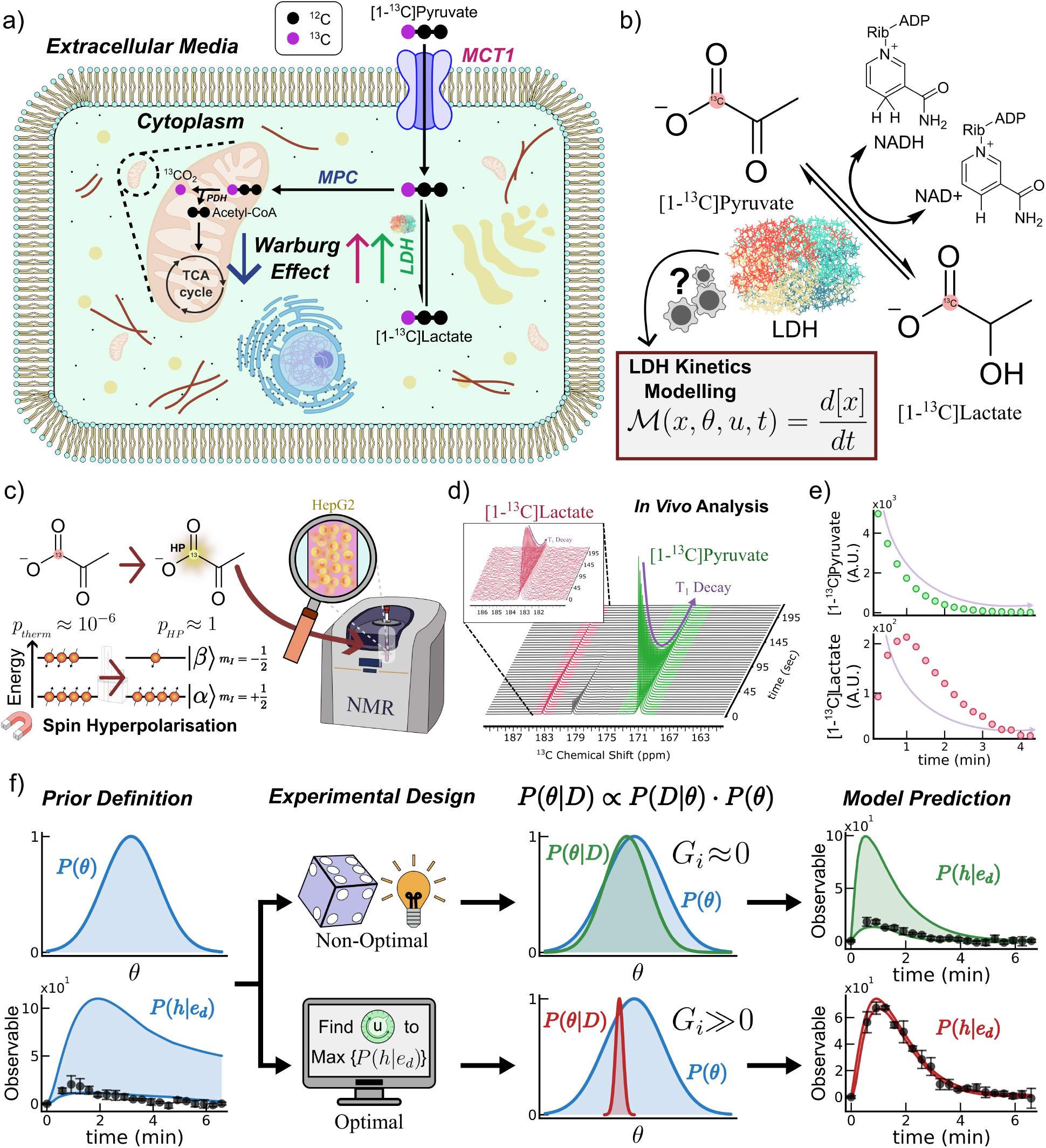
*In vivo* LDH kinetics identification platform. **a**, Schematic representation of an eukaryotic cell highlighting the role of LDH and exogenous pyruvate MCT1 transporter in the Warburg effect identification employing ^13^C labelled pyruvate. **b**, Reaction of interest with the labelled substrate ([1-^13^C]pyruvate) and product ([1-^13^C]lactate) chemical structures, with the reduced cofactor NADH and the catalysing LDH enzyme. We employ mathematical modelling to elucidate and understand the LDH action mechanics. **c**, *In vivo* experimental setup where we polarise [1-^13^C]pyruvate using dDNP, administer it to HepG2 suspensions enclosed in 5 mm diameter NMR tubes and measure the flux of the ^13^C signal in a benchtop NMR spectrometer. Example measured spectra stack (**d**) where HP pyruvate (green) is introduced to HepG2 cells, metabolising it to lactate (red) over time and peak integration (**e**). The purple lines represent the exponential polarisation signal loss attributed to *T*_1_. **f**, Schematic representation of the proposed bOED scheme compared to traditional experimental design methods.

## 2 Results

### 2.1 HP-NMR to probe *in vivo* metabolism

Metabolism dysregulation is a pivotal effect in human diseases. Here, we focus on studying the lactic fermentation of pyruvate. In cancerous cells, pyruvate metabolism flux shifts from respiration to cytosolic conversion to lactate via LDH (Fig. 1a-b). Thanks to HP-NMR, we can study this metabolic route in real-time and in a non-destructive manner (Fig. 1c). We introduced exogenous HP [1-^13^C]pyruvate to samples containing HepG2 cells in suspension, uptaking the substrate and metabolised it to [1-^13^C]lactate. We can observe both substrate and products with a single scan thanks to HP and its increase in signal sensitivity. Hence, using NMR, we scanned our samples over time (every 5 seconds) to observe in the spectra the consumption of pyruvate (Fig. 1d, green) and production of lactate (Fig. 1d, red), both subject to signal decay due to longitudinal relaxation time constant (i.e., exponential decay) and additional sources of signal loss (e.g. RF pulses). At this point, by computing the integrals of each peak, we quantified the amounts of observable HP pyruvate and lactate in our samples at every scan (Fig. 1e).

### 2.2 NMR linearity between cells and product

At a fixed pyruvate concentration, the lactate produced in HP-NMR experiments is directly proportional to the cell number *N* employed. We must ensure three main experimental conditions for this linearity. First, the amount of pyruvate molecules needs to be in excess. Throughout this work, we worked with 9 · 10^16^ (0.6 mM) to 6 · 10^18^ (40 mM) molecules of pyruvate per 2 − 8 · 10^6^ cells. Second, the total cell volume should be less than the total sampled volume (250 *µ*L for us). Third, both cells and pyruvate must be homogeneously distributed. We added exogenous HP pyruvate at 3.2 mM to samples containing two, four or eight million HepG2 cells (Fig. 2a). The results (*R*^2^ = 0.984) confirm the linearity, since the AUC for four million cells was approximately twice than for two million and half than for eight (control baseline experiments in Suppl. Fig. 4). Basic mathematical modelling aids in understanding this linearity. We used an oversimplified model [18] to describe the system (ℳ (*θ*_*α*_), Fig. 1e, blue). Here, we can decompose these kinetic rates as *N* times the kinetic rate *k*_*n*_, with *n* = {*PL, LP*} . We used Bayesian statistics to infer the posterior distributions for a unique set of rates *k*_*n*_ (*N* = 1 · 10^6^) and 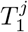 with *j* = {*P, L*} (Fig. 2f, blue). Besides *k*_*LP*_, which was poorly identified (*G*_*i*_ = 0.1), the rest of the parameters had a higher *G*_*i*_ than 2 (maximum of 4.6). Nonetheless, we could use this unique posterior to predict (Fig. 2g, blue shaded areas) experimental data (pyruvate curves in Suppl. Fig. 5a).

**Fig. 2:**
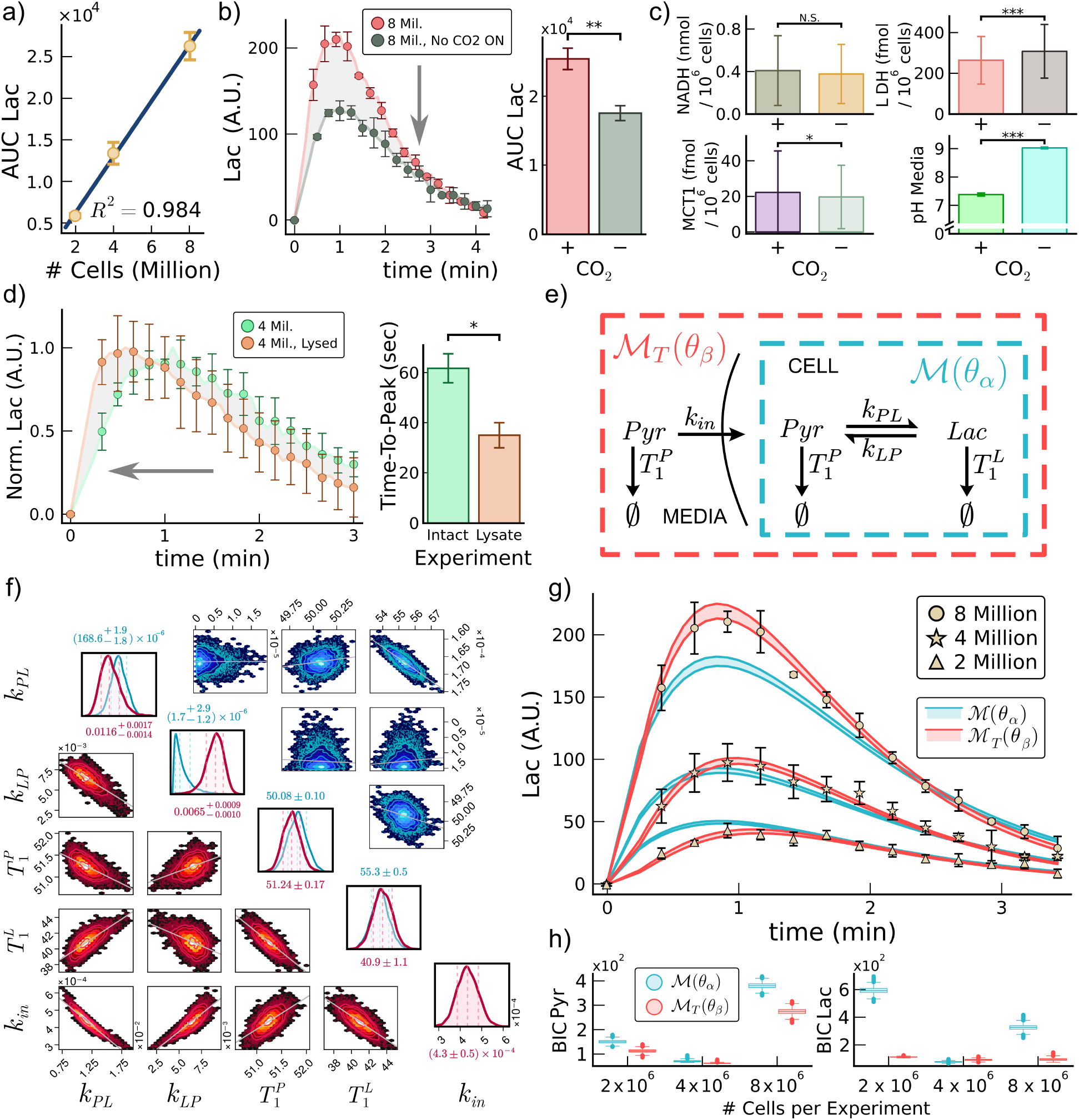
Reaction scalability and transporter effects. **a**, [1-^13^C]lactate produced AUC after administering [1-^13^C]pyruvate 3.2 mM using two, four and eight million HepG2 cells for experiments. **b**, Lactate production comparison (left) between eight million healthy HepG2 cells and a same parallel sample incubated without CO_2_ overnight, with their lactate AUCs (right). **c**, Measured variations in intracellular NADH (top left), intracellular LDH (top right), membrane MCT1 (bottom left) and extracellular media pH (bottom right) for HepG2 cells incubated with (+) or without (-) CO_2_ overnight. **d**, Lactate production (left) comparing intact 4 · 10^6^ HepG2 cells (green) to the *ex vivo* enzymatic activity (orange) of lysed ones, with the time to reach their maximum observable production (right). **e**, Reaction description for the traditional model (*θ*_*α*_) (blue) and expanded model with MCT1 transport ℳ_*T*_ (*θ*_*β*_) (red). **f**, ℳ (*θ*_*α*_) (blue, upper triangular matrix) and ℳ_*T*_ (*θ*_*β*_) (red, lower triangular matrix) posterior distribution inferred using the experiments from panel **a**. We scaled *k*_*in*_, *k*_*P L*_ and *k*_*LP*_ for one million cells. Dashed lines indicate 16, 50 and 84 % quantiles. **g**, HP-NMR [1-^13^C]lactate data used for parameter inference overlaid with ℳ (*θ*_*α*_) (blue) and ℳ_*T*_ (*θ*_*β*_) (red) posterior predictions (0.5-99.5 percentiles, shaded areas). **h**, ℳ (*θ*_*α*_) and ℳ_*T*_ (*θ*_*β*_) BIC for panel **g**. We performed all statistical pairwise comparisons with a T-Test (*: 0.05 ≥ p *>* 0.01, **: 0.01 ≥ p *>* 0.001, ***: 0.001 *>* p).

### 2.3 State perturbations break linearity

We can alter the linearity between lactate production and *N* by inducing cellular stress. Fig. 2b shows lactate production by eight million HepG2 cells administered with 3.2 mM pyruvate in ordinary culture conditions (red) and when excluding CO_2_ overnight (grey). The absence of CO_2_ negatively affected cell growth (i.e., absence of cell division) and viability (i.e., decrease of ≈ 2 % cell survival). We also observed a significant negative effect on lactate metabolism, with ≈ 31% reduction (2b, right). According to ℳ (*θ*_*α*_), the kinetic rate *k*_*P L*_ was approximately half if we maintained *N* = 8 · 10^6^. Hence, to better understand the decrease in metabolism, we quantified the main factors involved in the reaction (Fig. 2c). We observed a non-significant change in intracellular NADH (Fig. 2c, top-left) and a moderate decrease in MCT1 (Fig. 2c, bottom-left). Surprisingly, we observed an increase of LDH in the absence of CO_2_ (Fig. 2c, top-right). Yet, the most significant change was the increase in media pH from 7.4 to 9 (Fig. 2c, bottom-right), reducing by two orders of magnitude ^+^H, which are needed by MCT1. Such results stress the importance of maintaining consistent experimental conditions within and between research groups for comparability.

### 2.4 Factoring in MCT1 introduces delays

Using HP-NMR, we can directly measure the reaction contribution of MCT1 in lactate production. Despite being commonly shunned, cells can only uptake exogenous pyruvate through MCT transporters. Hence, due to this limiting factor, when removing this step, we should reach the maximum HP lactate observed sooner. To confirm this hypothesis, we compared HP-NMR lactate curves of four million HepG2 cells (Fig. 2d, green) with the corresponding extract of lysed cells (Fig. 2d, orange) and supplemented NADH (≈ 0.46 mM), showing indeed faster dynamics for the latter with a ≈ 27 second delay (Fig. 2d, right). We normalised the HP-NMR lactate curves to focus on the dynamics’ change, since lysed samples showed increased LDH activity, despite comparable AUCs with only a 1.13 times increase for the cell extract. Yet, due to the evident pyruvate transport effect, we expanded model ℳ (*θ*_*α*_) to introduce a transport term (i.e., *k*_*in*_) as model ℳ_*T*_ (*θ*_*β*_) (Fig. 2e, blue). We intended this new model to be a better representation of the LDH system.

### 2.5 Transport improves model predictions

Thanks to incorporating a transporter term *k*_*in*_, ℳ_*T*_ (*θ*_*β*_) significantly outperforms ℳ (*θ*_*α*_). To prove the advantages of ℳ_*T*_ (*θ*_*β*_), we inferred model parameters using the data from Fig. 1a. *G*_*i*_ was between 1.17 and 2.53 for all parameters, with a unified *k*_*j*_ (with *j* = {*PL, LP in* }) for one million HepG2 cells (Fig. 2f, red). Yet, we noted that *k*_*in*_ had high correlations with the other parameters. We observed improved model predictions (Fig. 2g, red shaded areas), with better time-to-peak matches (see two and four million cells) and signal intensities (see four and eight million cells). We also observed visual improvements in the HP pyruvate curve predictions (Suppl. Fig. 5b). Furthermore, by observing the BIC (Fig. 2h), ℳ_*T*_ (*θ*_*β*_) significantly outperforms ℳ (*θ*_*α*_) in predicting both observables generally. Since the metric penalises models for their complexity, the results validate ℳ_*T*_ (*θ*_*β*_) as a better model despite its extra parameter.

### 2.6 LDH repression at high pyruvate levels

LDH fails to reach a saturation level *in vivo* at increasing pyruvate concentrations, resulting in a decrease in lactate above a critical level. If we use ℳ (*θ*_*α*_) or ℳ _*T*_ (*θ*_*β*_)), the lactate production of both at increasing pyruvate concentrations will scale linearly (Fig. 3a) due to the lack of any rate-limiting step in the models. In Fig. 3b, we show the mean lactate production at increasing pyruvate concentrations. HP lactate data with error in Suppl. Fig. 6, and pyruvate in Suppl. Fig. 7 and Suppl. Fig. 4b. The data shows a decrease in the signal after 3.2 mM pyruvate, with similar productions when using 0.8 or 40 mM. Moreover, although their lactate AUCs are similar, the curve dynamics differ, with faster production at 40 mM. To ensure validity, we ensured linear correlation (*R*^2^ = 0.978) between the HP pyruvate administered and observed (Suppl. Fig. 12) and the absence of changes in cell viability or death (Suppl. Fig. 8). We identified a quick lactate production growth at low pyruvate concentrations (0-3.2 mM), followed by a steady decrease at higher concentrations (Fig. 3c, red). We further validated this behaviour with a conventional *ex vivo* LDH activity assay (LDHaa), showing faster dynamics (Fig. 3c, right y axis). These results demonstrate that ℳ (*θ*_*α*_) and ℳ _*T*_ (*θ*_*β*_) are only capable of predicting one fixed experiment (Fig. 3f-g, blue and red). Hence, we need better model formulations to predict and ultimately understand the LDH system.

**Fig. 3:**
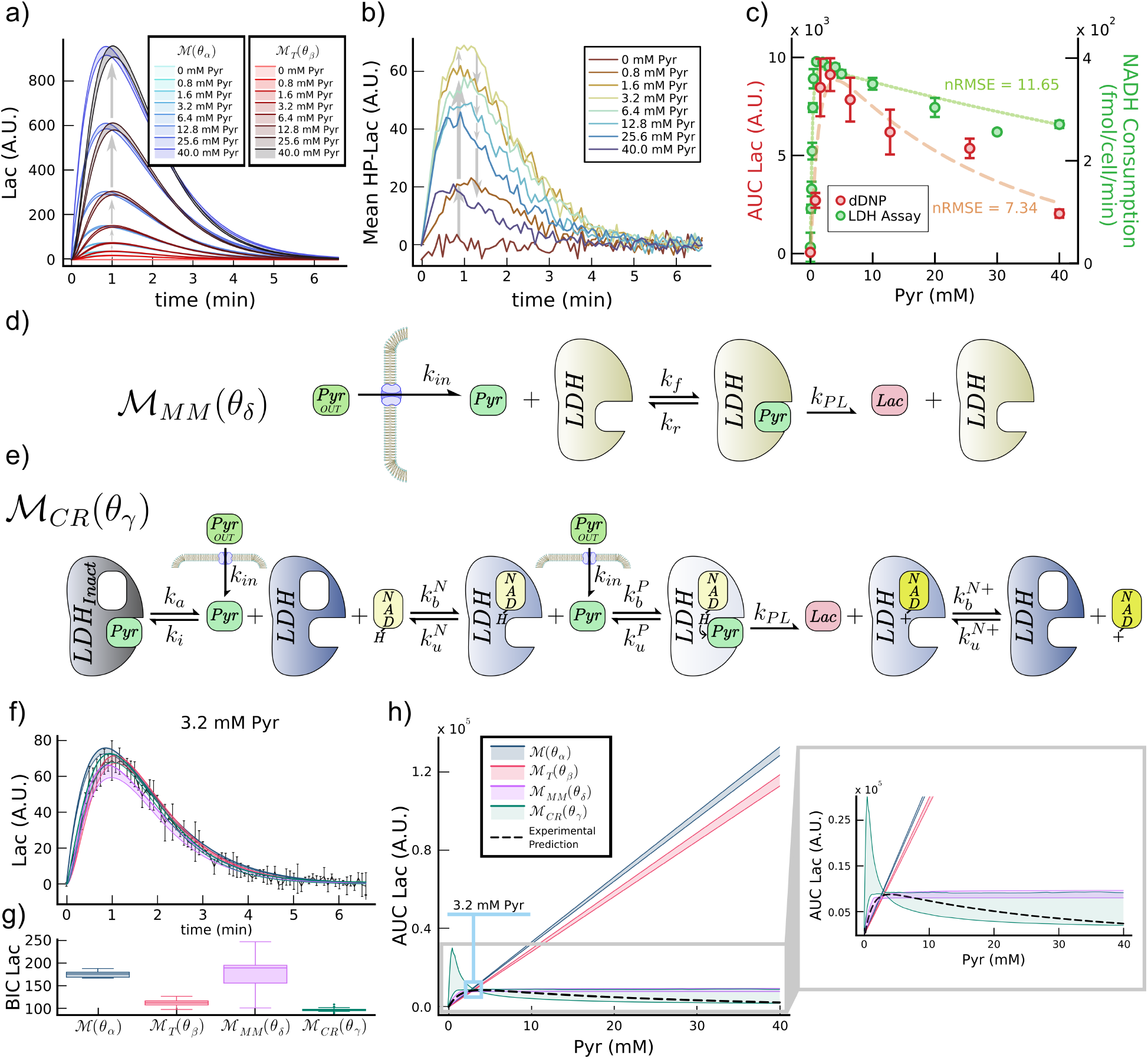
LDH repression at high pyruvate concentrations. **a**, ℳ (*θ*_*α*_) (blues) and ℳ_*T*_ (*θ*_*β*_) (reds) posterior lactate predictions for three million cells at increasing [1-^13^C]pyruvate concentrations. **b**, Experimental data means for the measured [1-^13^C]lactate production using the conditions from panel **a. c**, Experimental *in vivo* lactate AUC (red) and *ex vivo* NADH consumption (green) at increasing pyruvate concentrations. Dashed and dotted lines represent double negative exponential fitted functions. **d**, ℳ _*MM*_ (*θ*_*δ*_) schematic describing exogenous pyruvate transport and its conversion to lactate using basic Michaelis-Menten kinetics for LDH. **e**, ℳ _*CR*_(*θ*_*γ*_) schematic including MCT1 transport and LDH biologically relevant reaction steps for cofactor (NADH/NAD+) binding/unbinding and inactive species. **f**, Posterior predictive distribution for ℳ (*θ*_*α*_) (blue), ℳ _*T*_ (*θ*_*β*_) (red), ℳ _*MM*_ (*θ*_*δ*_) (pink) and ℳ _*CR*_(*θ*_*γ*_) (green) for an experiment using [1-^13^C]pyruvate at 3.2 mM and their BIC (**g**). **h**, All models *in vivo* posterior prediction for the substrate ([1-^13^C]pyruvate) to product ([1-^13^C]lactate) relationship of LDH. Dashed black line represents the fitted double exponential from panel **c** (red). Shaded areas represent model posterior predictions for the 0.5-99.5 percentiles. We used *N* = 3 · 10^6^ cells for all experiments.

### 2.7 Michaelis-Menten is structurally inadequate

Michaelis-Menten kinetics are limited to reaction saturation rather than the observed decrease in lactate production. The first step to avoid input-output linearity in models is to introduce biologically relevant enzymatic steps. Hence, using Michaelis-Menten kinetics, we developed ℳ _*MM*_ (*θ*_*δ*_) (Fig. 3d). With ℳ _*MM*_ (*θ*_*δ*_), it is the amount of cytoplasmic LDH that directly relates to the amount of lactate produced rather than the kinetic rates (except for *k*_*in*_). To account for its variability, we considered LDH concentration as an additional model parameter. After inferring model parameters (Suppl. Fig. 9), we observed that ℳ _*MM*_ (*θ*_*δ*_) predicted the experiment used for inference (Fig. 3f, purple) similarly to ℳ (*θ*_*α*_) and ℳ _*T*_ (*θ*_*β*_), yet with a much broader prediction uncertainty and BIC (Fig. 3g). Nonetheless, when using the model to predict the data from Fig. 3b (Suppl. Fig. 10a), ℳ _*MM*_ (*θ*_*δ*_) poorly predicted unseen experimental data (Fig. 3h, purple). The model accurately predicted experimental data from pyruvate 0 to 3.2 mM, at which point it reaches the characteristic steady state saturation, incapable of reproducing the observed LDH activity decrease by formulation. Hence, the results unveil the need for including more information from the biological system to predict experimental observations.

### 2.8 Competition between pyruvate and NADH

Experimental data, both *ex vivo* (Fig. 3c, green) and *in vivo* (Fig. 3c, red), show a higher level of LDH system complexity. For the reaction to occur, the cofactor needs first to bind to LDH, followed by pyruvate. If pyruvate binds first, it obstructs the entry of the cofactor, generating a pool of “inactivated” enzyme molecules [31, 32]. Consequently, the average number of inactive LDH molecules increases proportionally to the amount of pyruvate in the system. Hence, formulated the model ℳ _*CR*_(*θ*_*γ*_) (Fig. 3e). Yet, despite its biological relevance, the model comes with a higher complexity level, also introducing LDH and NADH/NAD+ as additional variables of the model. By introducing competition between substrate and cofactor, ℳ _*CR*_(*θ*_*γ*_) proves to be the superior model for predicting experimental data. Consequently, we inferred the model’s posterior parameter distributions (Suppl. Fig. 11) using the same 3.2 mM pyruvate experiment as for the other models (Fig. 3f), obtaining the best BIC scores (Fig. 3g). If we observe at ℳ _*CR*_(*θ*_*γ*_) predictions for increasing pyruvate concentrations (Fig. 3h, green), the model performs well at a single concentration (i.e., 3.2 mM) but has high uncertainty for the others (Suppl. Fig. 10b). Therefore, due to the model complexity (i.e., degrees of freedom), the informative content of the one experiment used for inference was insufficient. So, to better infer ℳ _*CR*_(*θ*_*γ*_) parameters, we need more experiments or highly informative data.

### 2.9 First bOED *in vivo* validation

We demonstrated for the first time *in vivo* in a biological scenario that OED successfully predicts highly informative experimental designs, introducing a novel Bayesian approach (bOED) for the task (Fig. 1f). For a mathematical model ℳ, our algorithm aims at finding the system input *u* (e.g., pyruvate concentration) that maximises the model’s prior prediction uncertainty. Here, we used ℳ _*CR*_(*θ*_*γ*_), comparing the bOED strategy to traditional experimental design approaches. For all experimental designs, we implemented three iterative steps with one common experiment for *Itr*_1_, followed by *Itr*_2_ and *Itr*_3_ designed using each approach (Suppl. Fig. 13). In all iterations, bOED outperformed the other design approaches, both in theory 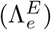 and in minimising posterior uncertainty (*G*_*i*_) after inference (Fig. 4a). Moreover, the non-perfect linear relationship between *G*_*i*_ and − *log*(|Σ|) shown in Fig. 4b showcases the lack of Gaussian behaviour in biological contexts. All the results aligned with the computational validation of the algorithm, further corroborating bOED for designing highly informative experiments for parameter inference (Suppl. Section B). We observed a mix of posterior results between experimental designs, with poorly identified parameters in all (Fig. 4c, top), highly identified ones (Fig. 4c, bottom) and parameters whose posterior drastically changed between designs (Fig. 4c, middle). Other parameters, such as *k*_*bp*_ or *k*_*a*_, were only highly identified when using bOED experiments. Overall, we observed a high correlation between 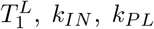 and *y*_0_s in all posteriors, with moderate (*k*_*IN*_ in the random design and 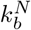 for the other two) to none amongst the rest (Suppl. Fig. 14). Hence, we demonstrated the superiority in posterior uncertainty reduction when using bOED to design experiments.

**Fig. 4:**
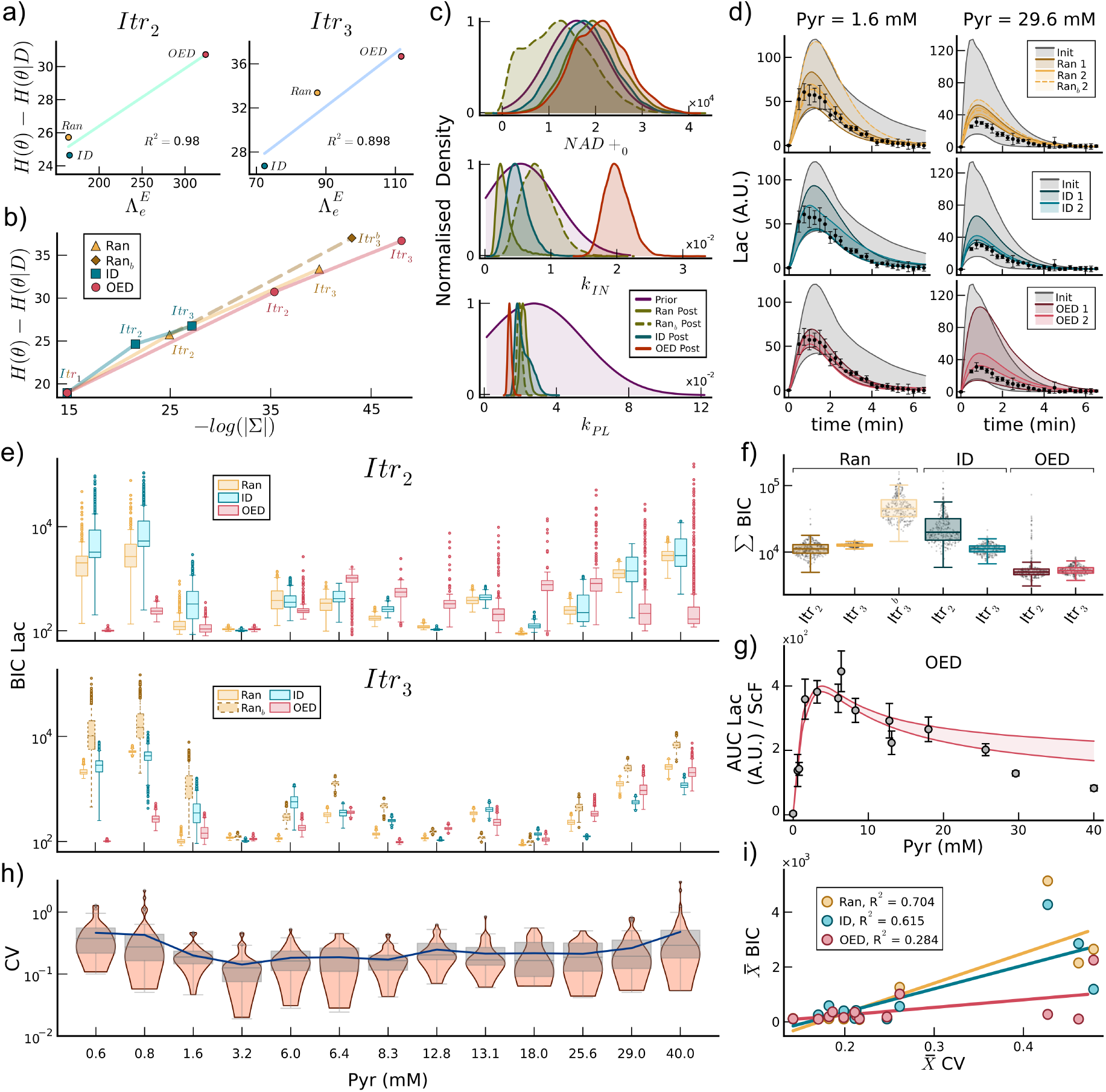
*In vivo* bOED validation using ℳ_*CR*_(*θ*_*γ*_). **a**, Correlation between prior prediction uncertainty and *G*_*i*_ of the main experimental designs at the second (*Itr*_2_, left) and third (*Itr*_3_, right) round of sequential inference iterations. **b**, Relationship between *a posteriori* informative content metrics. **c**, Example kernel density estimates of prior and posteriors at *Itr*_3_. **d**, Posterior predictions at each iteration for each experimental design for two unseen validation experiments. **e**, ℳ_*CR*_(*θ*_*γ*_) BIC for all the experiments generated in this work using all posteriors from *Itr*_2_ (top) and *Itr*_3_ (bottom) and their cumulative BICs (**f**). **g**, ℳ_*CR*_(*θ*_*γ*_) substrate (i.e., [1-^13^C]pyruvate) to product (i.e., [1-^13^C]lactate) relationship prediction after inference using bOED experiments (*Itr*_3_). **h**, HP lactate coefficient of variation for experimental data points outside of the base noise level. **i**, Correlation between the average CV and BIC for all experiments in this work using all *Itr*_3_ posteriors. Simulations shaded areas represent model posterior predictions for the 0.5-99.5 percentiles. We used *N* = 3 · 10^6^ cells for all experiments.

### 2.10 bOED enhances model prediction

Using bOED also results in better and more general predictive models. If we analyse predictions of two validation experiments (Fig. 4d), we observe an adequate progressive improvement with all experimental designs. Nonetheless, if we observe the general predictions, the posteriors obtained with bOED data outperform the other two designs (Fig. 4e and Fig. 4f). When using posteriors obtained using Ran or ID experiments, at *Itr*_2_, the model poorly predicted lactate production at low (i.e., < 3.2 mM) and high (i.e., > 18 mM) pyruvate concentrations. For bOED posteriors, it did better on average when observing the means (Fig. 4e, top). At *Itr*_3_, the bOED posterior obtains the best predictions overall, contrary to the other two design schemes (Fig. 4e, bottom). Fig. 4f indicates that even at the second iteration, bOED experiments yielded the best predictions (Fig. 4g), with an average cumulative BIC 2.35 times better than for Ran and 2.1 times better than for ID. To emphasise the advantages of bOED, we also evaluated an alternative third random design 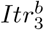. In this case, the posterior uncertainty improvement was as high as for bOED (Fig. 4b). Yet this posterior resulted in the worst individual (Fig. 4e) and overall (Fig. 4f) predictions of the system. We display all the predictions of lactate area under the curves (AUC) and two additional summary statistics in the Suppl. Fig. 15. Furthermore, when analysing the noise levels of the experimental data, we found that experiments using low and high pyruvate concentrations resulted in higher coefficients of variation (Fig. 4h). The results correlate with the experimental low lactate production levels. Computationally, we validated that increases in experimental noise result in worse posterior uncertainties (Suppl. Fig. 16). However, we observed that experiments designed with bOED achieved a decorrelation CV and model prediction quality (Fig. 4i).

### 2.11 ℳ_CR_ improvement with informative priors

Thanks to the bOED and experimental measurements of cellular LDH (Fig. 5a, right, measured with flow cytometry) and NADH/NAD+ (Fig. 5a, left, measured with absorbance quantification assay) levels, we fine-tuned ℳ_*CR*_(*θ*_*γ*_) to be highly biologically relevant. Since we fully validated bOED, we employed it to select the best HP-NMR experiments from all performed ones in this work to infer ℳ_*CR*_(*θ*_*γ*_) parameters and initial states (Suppl. Fig. 17). Fig. 5b displays the 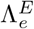 for all experiments at each iteration, showing the change in informative content of experiments at each iteration caused by the use of updated priors at each step. As expected, each inference iteration improved parameter uncertainty, avoiding stagnation and proving that every experiment used provided information for the posterior update (Fig. 5c). We observed a general progressive *P* (*θ D*) improvement for parameters, contrary to the initial states *y*_0_. We observed a high similarity between prior and posteriors for NADH (Fig. 5d, middle) and NAD+ (Fig. 5d, top), and a similar uncertainty for LDH, with an apparent underestimation revealed by the posterior means (Fig. 5d, bottom). The only correlation we observed in some steps was between the *LDH*_0_ and the parameter *k*_*IN*_, and between the *NADH*_0_ and 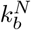 (Suppl. Fig. 18). Similar to previous inferences, we observed poorly (Fig. 5e, top) and highly (Fig. 5e, bottom) identified parameters, while others changed or improved across the different sequential inference steps (Fig. 5e, middle). Following the same trend, we identified moderate to high correlations between 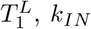 and *k*_*P L*_ and moderate ones between these three and 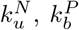 and 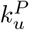 (Suppl. Fig. 18).

**Fig. 5:**
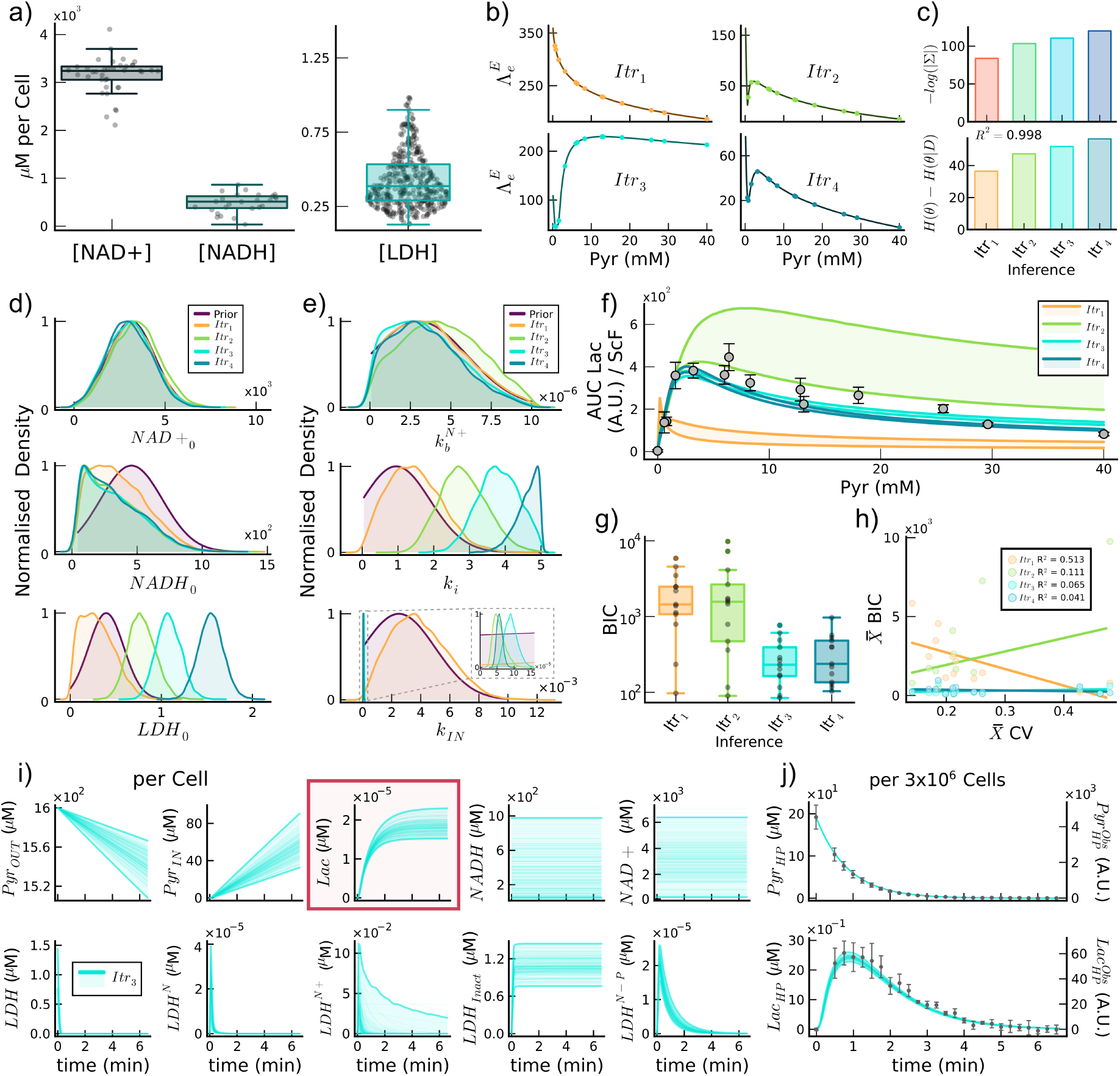
ℳ_*CR*_(*θ*_*γ*_) refinement and predictions. **a**, Experimentally measured cofactor NADH/NAD+ (left) and LDH enzyme (right) concentrations per HepG2 cell. **b**, bOED utility function prediction for all the HP pyruvate concentrations used in this work. Each iteration *Itr*_*n*_ where *n* > 1 uses as prior the posterior obtained in the previous iteration after inference using the experiment with higher 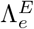. *Itr*_1_ employed an uninformative prior. **c**, Posterior covariance metric (top) and *G*_*i*_ (bottom) obtained from each optimal experimental inference. *y*_0_ (**d**) and example *θ* (**e**) priors and posteriors kernel density estimates. **f**, ℳ_*CR*_(*θ*_*γ*_) substrate to product relationship posterior predictions for each iteration. Dashed lines represent a double negative exponential fitted function. **g**, Cumulative BIC for all HP-NMR experiments in this work (scatter plot) for each posterior obtained after each sequential inference iteration. **h**, Correlation between the average CV and ℳ_*CR*_(*θ*_*γ*_) BIC for all experiments in this work using all the posteriors inferred after each sequential inference. **i**, ℳ_*CR*_(*θ*_*γ*_) system states *y* posterior predictions for an experiment using 1.6 mM [1-^13^C]pyruvate, using the *P* (*θ* | *D*) obtained at *Itr*_3_ for one average cell. All concentrations are referenced to the total sample volume. **j**, System observable predictions for the experiment from panel **i**. Shaded areas in simulations represent model posterior predictions for the 0.5-99.5 percentiles. We used *N* = 3 · 10^6^ cells for all experiments.

### 2.12 Biological data refines ℳ_*CR*_ predictions

We got a model with biologically representative model parameter posteriors, capable of predicting the general *in vivo* LDH behaviour. When predicting the system as a whole, we observed a quick, progressive improvement, with an underestimation in *Itr*_1_, overestimation in *Itr*_2_, and a good result in *Itr*_3_ (Fig. 5f). Despite a reduction in posterior uncertainty, *Itr*_4_ barely improved predictions with respect to *Itr*_3_. Looking at cumulative BIC, *Itr*_3_ obtained the best results (Fig. 5g), suggesting a trade-off between posterior and prediction quality in *Itr*_4_. Individual lactate predictions for all iterations in Suppl. Fig. 19. The BIC for individual experiments (Suppl. Fig. 20) exhibited similar trends to the observed predictions in Fig. 5f, with the third and fourth iterations yielding the best results. Aligned with previous results, we observed a steep progressive decorrelation between model predictions and data quality measured with CV (Fig. 5h), further validating our bOED strategy.

### 2.13 Modelling unravels LDH state transitions

Besides experimental observables, mathematical models support the understanding of all biological states involved. Hence, we can simulate the different states of ℳ_*CR*_(*θ*_*γ*_) to comprehend the state transitions in the LDH reaction (Suppl. Fig. 21). When observing individual cell behaviour (Fig. 5i), we see that the model predicts a low steady intake of exogenous pyruvate (*Pyr*_*OUT*_ and *Pyr*_*IN*_). Only a small amount of this pyruvate seems to be metabolised into lactate (*Lac*, red square), reaching a steady state at around four minutes. We also observe a high prediction uncertainty with *NADH* and *NAD*+. Only a small amount of the cofactor binds to *LDH*, fully bound in less than 30 seconds. We can also discern a peak at ≈ 20 seconds followed by a steep decrease in LDH intermediates (i.e., *LDH*^*N*^, *LDH*^*N*+^ and *LDH*^*N*−*P*^), due to the high level of enzyme inactivation (*LDH*_*Inact*_). Furthermore, the model shows a good prediction of pyruvate (Fig. 5j, top) and lactate (Fig. 5j, bottom) for the overall HP-NMR experiment. Hence, the results demonstrate the outstanding capabilities of the proposed ℳ _*CR*_(*θ*_*γ*_) model to describe the LDH system, at least *in vivo*.

### 2.14 *Ex vivo* LDH activity breaks *M*_*CR*_(*θ*_*γ*_)

*ℳ* _*CR*_(*θ*_*γ*_) enables the exploration of transporter effects, successfully describing LDH kinetics *in vivo* and *ex vivo*. While HP-NMR enables us to observe the intracellular LDH behaviour (Fig. 6a, red), an LDHaa excludes transport and the cellular crowded microenvironment (Fig. 6b, cyan). Hence, to ultimately understand the system’s behaviour, we hypothesised two alternative scenarios: 1) HP-NMR experiments without MCT1 (Fig. 6a, green), and 2) LDHaa requiring MCT1 transport (Fig. 6b, pink). Fig. 6c shows a faster LDH inhibition *ex vivo* prediction at a lower pyruvate concentration (≈ 0.05 mM) compared to when factoring in MCT1 (≈ 2.7 mM). The prediction of LDHaa experiments shows the same trends (Fig. 6d), yet with higher prediction uncertainty. Both results align with the trends observed in Fig. 3c, but with discrepancies in concentrations and inhibition degree. The *ex vivo* enzymatic dynamics were much faster (Suppl. Fig. 23) compared to ℳ_*CR*_(*θ*_*γ*_) predictions (Fig. 3d, cyan). Note that the lactate concentration predicted (Fig. 3c, red) was higher than the one extrapolated experimentally (Suppl. Fig. 22) because of the longitudinal relaxation decay 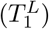. According to the model, the maximum lactate observed (≈ 55 seconds after mixing) was 60.36 *±* 3.2 % of the predicted system at steady state.

**Fig. 6:**
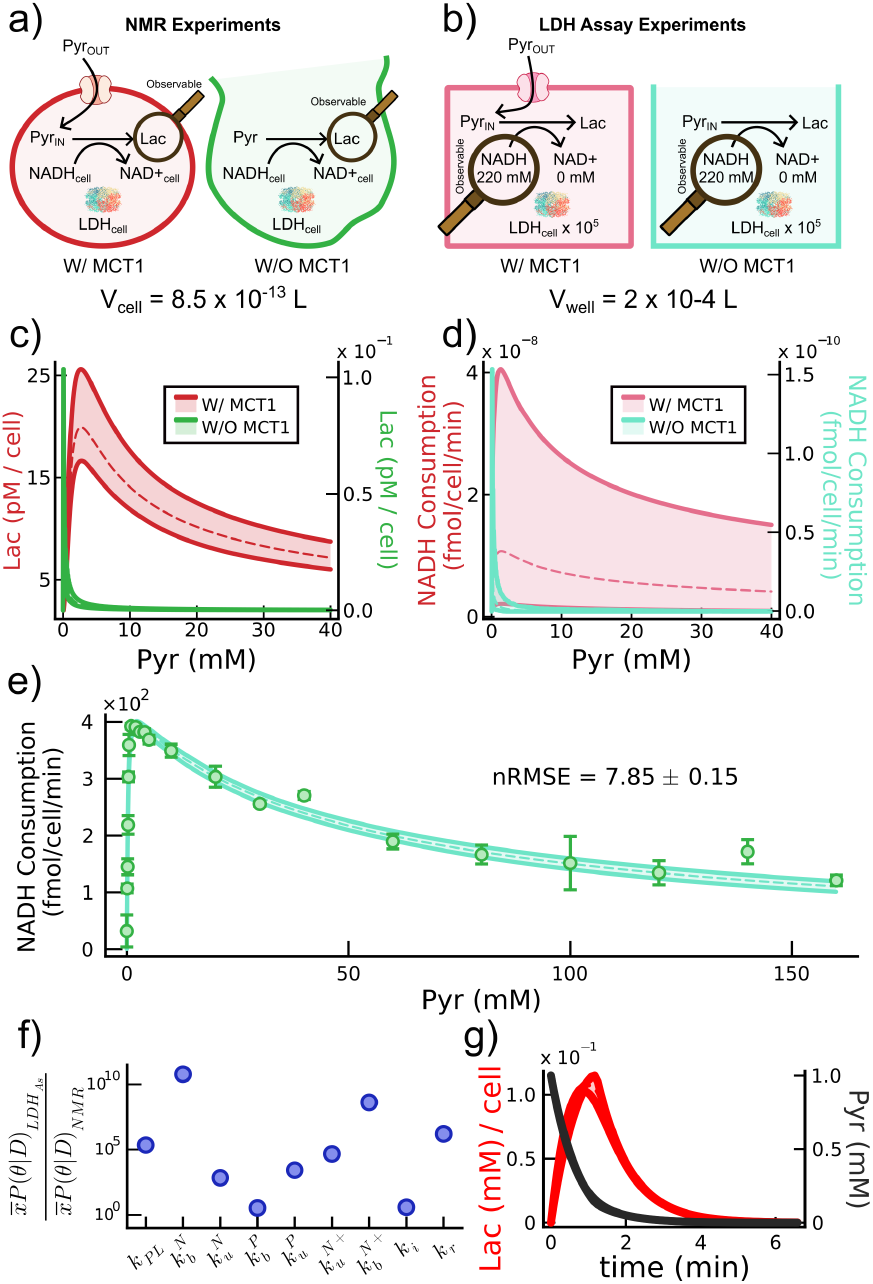
*In vivo* against *ex vivo* LDH activity. Schematics for the HP-NMR experiments (**a**, red) also excluding MCT1 transport (**a**, green) and LDHaa (**b**, cyan) also including MCT1 transport (**b**, pink). ℳ_*CR*_(*θ*_*γ*_) posterior predictions including transport (**c** red and **d** pink) and excluding it (**c** green and **d** cyan). **e**, ℳ _*CR*_(*θ*_*γ*_) NADH consumption posterior prediction (shaded area) after inference using all available data (scatter plot). **f**, Comparison of ℳ _*CR*_(*θ*_*γ*_) posterior parameter means after inference using LDHaa experiments 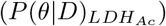 over HP-NMR ones (*P* (*θ* | *D*)_*NMR*_, *Itr*_3_ from Fig. 5). **g**, Posterior prediction for the lactate observed per one HepG2 cell in a hypothetical HP-NMR experiment using 1 mM [1-^13^C]pyruvate (right y axis) using 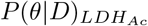. Shaded areas represent posterior predictions for the 0.5-99.5 percentiles. Dashed lines indicate means.

### 2.15 Universal *M*_*CR*_(*θ*_*γ*_) predictability

To demonstrate the correctness of ℳ_*CR*_(*θ*_*γ*_), we inferred the model parameters (excluding *k*_*IN*_ from the equations) using the LDHaa experimental data. As displayed in Fig. 3e, ℳ_*CR*_(*θ*_*γ*_) accurately predicted the experimental data with the updated set of parameter posteriors. While we attempted to infer the model parameters using *in vivo* and *ex vivo* data together, no parameter could represent both. Parameter means obtained in this inference were from three times 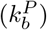 to ten orders of magnitude 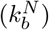 higher compared to the HP-NMR ones (Fig. 3f). Surprisingly, the binding rate for NAD+ 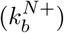 was also eight orders of magnitude higher, introducing a new species pool of inactivated LDH. As for parameter correlations, we observed moderate ones for *LDH*_0_ with most parameters (Suppl. Fig. 24). We elucidated the much faster system dynamics when using 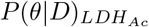 to predict an HP-NMR experiment and observing the lactate trend (Fig. 3g). The pyruvate curve (Fig. 3g, black) decayed quickly, dominated by both longitudinal relaxation decay and high substrate consumption by LDH. While the lactate curve had the correct shape (Fig. 3g, red), only one HepG2 cell would consume one tenth of the pyruvate, making it incompatible with the *in vivo* data. If we observe the model predictions for all model states, the faster dynamics are better elucidated by 1) the fast consumption of pyruvate and NADH and 2) the high concentration reached by the inhibitory species *LDH*_*Inact*_ and *LDH*^*N*+^ (Suppl. Fig. 25). Hence, ℳ_*CR*_(*θ*_*γ*_) provides an accurate general representation of LDH kinetics, but the model is susceptible to different microenvironment conditions.

## 3 Discussion

Despite the biological and health importance of LDH and advances in structural crystallography [32], a lack of consensus persists regarding its actual enzymatic mechanics [12, 18, 19, 33, 34]. In this study, we combine biological experimentation with mathematical modelling to reveal the enzyme’s kinetic mechanism of action while pondering the impact of diverse elements involved in the reaction (e.g., NADH or MCT1 transporter). For simplicity, we used HepG2 cells, which predominantly express the LDH isozyme 5, favouring the production of lactate [35]. As a first characteristic behaviour, when examining the LDH substrate-to-product behaviour, the enzyme first displays a quick increase in lactate production, followed by a steady decrease at higher substrate concentrations. Yet, we are the first to provide experimental data displaying this behaviour *in vivo*, accounting for cellular context (Fig. 3c, red). Moreover, to unravel rate-limiting steps, we also characterised the concentrations of MCT1 (Fig. 2c), LDH (Fig. 5a) and NADH (Fig. 5a). As expected, the concentration of LDH enzyme was three orders of magnitude lower than the coenzyme, since NADH/NAD+ intervene in hundreds of reactions [36]. These results prove that the enzyme is a reaction-limiting factor. Yet, the highest limiting component we identified due to its low concentration was the MCT1 transporter, underlining its pivotal role in the reaction [37].

To obtain *in vivo* LDH experimental data in real-time, we leveraged the metabolic imaging capabilities of HP-NMR. First, we validated that when using a fixed pyruvate concentration, the lactate produced linearly correlated with the number of cells used (Fig. 2a). Hence, the increase in cancerous cells in a tumour partly explains the increase in LDH metabolism [38]. HP-NMR also allowed us to validate the importance of the MCT1 pyruvate transporter (Fig. 2d) with further consolidation using mathematical modelling (Fig. 2f-h). However, a major limitation of HP-NMR is the short observation window of the HP signal (≈ 4-5 minutes at best), which is largely governed by the *T*_1_ of the substrate and its products. Moreover, the variety of hyperpolarisable ^13^C molecules is quite limited, with pyruvate being the best one. Hence, the technique can identify effects but fails at providing information on causes. An example of this limitation was our study on the effect of CO_2_ cell incubations. While HP-NMR identified a decrease in lactate production (Fig. 2b), we needed other techniques to understand the cause of the changes (Fig. 2c). Our results indicate that changes in LDH, MCT1 or NADH were nonexistent or incapable of explaining the depletion in metabolism. The main cause of the decrease in metabolism was the increase in media pH, reducing the concentration of available ^+^H for the transporter. Hence, lactate overproduction can be a side effect of the common acidic media in cancerous regions [39].

Researchers elucidated the complex *ex vivo* LDH kinetics as early as the 1960s [40]. We replicated these experiments using HepG2 cells to compare them with our *in vivo* data and identify any differences. First, the assay will never include the rate-limiting effects of MCT1. Furthermore, the assay disregards the crowded cellular microenvironment and homeostatic controls [14]. We attribute this lack of cellular contextualisation as the main cause of the differences in LDH behaviour. With our computational analysis, we revealed that the *ex vivo* and *in vivo* reactions were intrinsically different, with the former being faster. Our analysis suggested that MCT1 alone was incapable of explaining the increase in LDH speed dynamics. Another element justifying the lack of cellular microenvironment effect is that the NADH used for the assay is similar to the total intracellular amount [12], rather than the cytosolic one. Moreover, the use of a sample volume of 200 *µ*L in the assay resulted in the total extracted LDH enzyme being less concentrated (≈ 8.5 x 10^−4^ *µ*M) than that inside a single cell (≈ 0.4 *µ*M). We also observed this increase in enzyme dynamics in HP-NMR experiments (Fig. 2d), where, despite the much lower NADH concentration in the lysed sample (0.46 *µ*M), the maximum peak intensity was 1.45 times higher than for intact cells (data normalised for visualisation purposes).

To fully understand and describe the LDH system’s behaviour, we used mathematical modelling. In the HP-NMR field, researchers commonly rely on kinetic rates derived from intrinsically wrong (e.g., see ℳ (*θ*_*α*_)) linear models [18]. These models are incapable of reproducing general experimental data due to their oversimplification (Fig. 3h). The first improvement is the addition of a transporter term for pyruvate (*M*_*T*_ (*θ*_*β*_)), rarely found across literature. Furthermore, LDH lacks the characteristic enzyme saturation profiles of Michaelis-Menten kinetics (ℳ_*MM*_ (*θ*_*δ*_)). Hence, we selected a model structure that follows competitive repression of LDH by substrate (ℳ_*CR*_(*θ*_*γ*_)), in accordance with our experimental data, literature one [40], and molecular structural information [32]. Yet, the main limitations of the model are the assumptions of no backwards reaction (valid for HepG2) and the assumption of the absence of intracellular pyruvate. Moreover, while we focused on *LDH*_*Inact*_ as the main inhibitory species, *ex vivo* data highlighted *LDH*^*N*+^ as a second major inactive species (Suppl. Fig. 25). While we observed the *LDH*^*N*+^ effect when predicting *in vivo* data (Fig. 5i), the complex had a lower contribution to the total LDH inactivation. Finally, ℳ_*CR*_(*θ*_*γ*_) high level of biological detail emphasised the importance of experimentally measuring (when possible) initial *y*_0_ states (i.e., *LDH, NADH, NAD*+) to generate informative priors for Bayesian inferences and ultimately get biologically relevant models (Fig. 5).

We leveraged Bayesian statistics to update informed prior beliefs, discarding general frequentist assumptions. The advantages of Bayesian inference become clear in biological contexts, where model parameters and predictions are rarely normally distributed [41]. This lack of normality becomes apparent in the imperfect correlation between *G*_*i*_ and − *log*(|Σ|) (Fig. 4b). Moreover, Bayesian inferences aid in determining parameter identifiability issues (Fig. 4c top and Fig. 5d-e top) and parameter correlations (Fig. 2f), such as we generally see between *k*_*IN*_ and *k*_*P L*_. Moreover, we can encode previous experimental information in *P* (*θ*) to reduce identifiability issues and obtain a more biologically sound model, as we did for *y*_0_s. For example, we saw that the *θ* means changed significantly when using informative *y*_0_ priors (Fig. 5) compared to when using uninformative ones (Fig. 4). However, to this day, Bayesian inferences rely on sampling-based methods [42], which come at high computational costs. For example, to infer one experiment (Fig. 3f), ℳ (*θ*_*α*_) took about five times less duration than ℳ_*T*_ (*θ*_*β*_) and 498 times less than for ℳ_*CR*_(*θ*_*γ*_). For this reason, one should ensure to obtain highly informative data for parameter inference, since different experiments provide different amounts of information [25].

Here, we present a novel bOED algorithm, which is computationally feasible even for complex models. Furthermore, for the first time in a biological context, we validated the robustness and soundness of an OED algorithm *in vivo* (Fig. 4). Its introduction in biology has been limited to computational works, generally following frequentist statistics [30] due to the excessive complexity of Bayesian methods [28, 29]. Our algorithm is practical and time-feasible, even for highly dimensional models, highly sensitive to prior beliefs (Fig. 5b). First, in the *in silico* bOED validation (Suppl. Section B), we demonstrated the strength of our bOED algorithm to outperform other design approaches. These results directly translate to *in vivo* settings (Fig. 4a). However, despite its consistent positive results, our bOED presents some drawbacks. When using highly uninformative priors, the algorithm identifies any experiment as a good one to gain information. Besides, while we can use bOED for simpler and quicker experiments, we regard it as unnecessary. Hence, for the LDHaa (Fig. 6), we used an intuition-derived approach, since measuring additional conditions was faster than performing computations. Yet, we foresee further future applications for our bOED. For example, even in frequentist scenarios, one could use normal distributions drawn from overestimated confidence intervals. Moreover, it would be valuable to test bOED to focus on non-observable states to improve prediction uncertainties, since informative posteriors might not translate to good predictions.

We calibrated ℳ_*CR*_(*θ*_*γ*_) to accurately predict *in vivo* LDH kinetics in HepG2 cells thanks to bOED. Yet, for a correct verification of any model, we need one crucial component: the generation of a validation set [43]. For this reason, we used additional HP-NMR experiments as validation datasets to assess the general predictive capabilities of our models. It is also important to choose adequate quality metrics (e.g., BIC or nRMS) that penalise models by their complexity. Furthermore, we also want to emphasise the need for reproducibility in experimentation. For HP-NMR experiments, we have an inherently low product signal, characterised by a basal homoscedastic noise level added to the biological heteroscedastic one. At both low and high pyruvate concentrations, we obtained the lowest lactate intensity signals with lower quality (Fig. 4h). These experiments have the worst model predictions, except when using bOED. Despite the higher gain in information provided by bOED, we can reach a saturation point for either posterior uncertainty reduction or prediction quality improvement. Hence, one should consider the trade-off between computational and experimental resources (Fig. 5, *Itr*_4_). Moreover, we also recommend bOED to avoid generating experiments that have a deleterious effect on predictions (Fig. 4,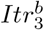). Finally, good predictive models permit the investigation of the non-observable kinetics. For example, in Fig. 5i, we identified low cellular pyruvate uptake, rapid lactate and *LDH*_*Inact*_ saturation, low cofactor consumption, and accelerated intermediate states reduction.

In summary, we fully characterised the intracellular LDH mechanism of action for HepG2 in depth thanks to predictive mathematical modelling and experimentation using a combination of biological techniques. Moreover, we introduced a novel bOED algorithm to efficiently design highly informative experiments for parameter inference, validating it *in vivo* for the first time in a biological context.

## 4 Methods

### 4.1 Cell culture

We cultured HepG2 human hepatocarcinoma cells (CliniSciences) in Eagle’s Minimum Essential Medium (EMEM) containing L-Glutamine (ATCC-30-2003), supplemented with 10 % Fetal Bovine Serum (Gibco, ThermoFisher) and 1 % penicillin-streptomycin (10^4^ *U/mL*) P/S (Gibco, ThermoFisher). We grew cells in T175cm^2^ flasks and incubated them at 37 ^°^C in a humidified atmosphere at 5 % (v/v) CO_2_. Where indicated, we cultured HepG2 in absence of CO_2_ in a portable incubator (N-BIOTEK) at 37 ^°^C. We passaged cells every 72 h or before reaching 80 % confluency as required. For spectroscopic experiments, we seeded cells at least 48 h before each measurement in T75cm^2^ flasks. On the day of the experiment, we retrieved cells by trypsinisation. Then, we determined the cell number with 0.4 % trypan blue exclusion stain using an automatic cell counter (Countess™3.0, ThermoFisher). We resuspended the cell suspensions with fresh medium for a final total volume of 250 *µ*L, including cells, media and [1-^13^C]pyruvate.

### 4.2 Cell viability

We resuspended 1 million HepG2 cells in a mixture of medium and buffered [1-^13^C]pyruvate remainders from experiments for 5 min to replicate exposure in experiments. We sampled nine pyruvate concentrations ranging from 0 to 40 mM (Suppl. Fig. 8). First, we determined cell viability (by triplicate) via trypan blue staining in an automatic cell counter (Countess™3.0, ThermoFisher). Second, we measured (by triplicate) the metabolic effect of the same pyruvate concentrations using alamarBlue HS assay (ThermoFisher) in a microplate reader (Synergy H1, BioTek) with 560 nm excitation and 590 nm emission filters.

### 4.3 Cell lysate experiments

After quantifying cell number, we incubated the cells with 50 *µ*L of lysis buffer free of sodium deoxycholate and SDS to minimize protein denaturalisation (50 mM Tris HCl pH 8.5, 120 mM NaCl, 5 mM EDTA, 1% Triton X-100) and for ≈ 30 min at 4 ^°^C. We also added protease inhibitor capsules following manufacturer’s intructions (cOmplete Mini, Roche). Then, we added 0.46 *µ*M of NADH per 1 million cells as the reaction cofactor and cell media for a total final volume of 250 *µ*L (i.e., lysate + media + buffered [1-^13^C]pyruvate).

### 4.4 LDH activity assay (LDHaa)

To assess the extracellular HepG2 LDH activity, measured as NADH consumption, we followed the methodology described by T.H. Witney *et. al*. in [12]. We performed the assay in a buffer containing 200 mM of potassium chloride (KCl) and 50 mM 4-(2-hydroxyethyl)-1-piperazineethanesulfonic acid (HEPES), adjusting pH to 7.1 with 1 M sodium hydroxide (NaOH). We added to the buffer reduced nicotinamide adenine dinucleotide (NADH) to a final concentration of 0.22 mM. We used the same lysis buffer described in the previous section (i.e., *Cell lysate experiments*) to extract the LDH enzyme from 1 · 10^5^ cells per assay. Right before measurement, we added the desired amount of sodium pyruvate to obtain a final pyruvate concentration ranging from 0 to 160 mM. We measured the NADH absorbance at 340 nm using a microplate reader (Synergy H1, BioTek) at 37 °C. We scanned each assay sample in quadruplicate for five minutes every 30 seconds, subtracting pyruvate’s absorbance (unconsumed) at the same wavelength. Consumption rates per minute are at the first minute of the reaction.

### 4.5 Cellular NADH quantification

We quantified NAD+ and NADH levels using the NAD/NADH Quantification Kit (MAK037, Sigma-Aldrich) according to the manufacturer’s instructions. We washed 2·10^6^ cells with cold PBS, centrifuged and mixed them with the NADH/NAD extraction buffer for cell lysis. Next, we centrifuged the lysates at 13,000 g for 10 min, and used the supernatant for analysis. Then, we transferred the samples for NAD_*total*_ and NADH quantification into a 96-well plate and measured the absorbance change at 450 nM with a microplate reader (Infinite M200 PRO, Tecan). We calculated the NAD+ concentration as NAD_*total*_ - NADH.

### 4.6 Cellular LDH and MCT1 enzyme quantification

We estimated the concentration of intracellular LDH enzyme and membrane monocarboxylate transporter MCT1 enclosed in 1 · 10^6^ HepG2 cells through fluorescent antibody staining and flow cytometry quantification. For LDH enzyme, after fixing and permeabilising cells, we stained them with Alexa Fluor™ 647 Rabbit Anti-Human LDH antibody BDBiosciences) at 9 *µ*g*/*mL during incubation. For MCT-1, we stained the cells with Human MCT1/SLC16A1 Alexa Fluor™488-conjugated antibody (R&D systems) at 50 *µ*g*/*mL during incubation. We analysed the samples using a Gallios multi-color flow cytometer instrument (Beckman Coulter) with a red excitation laser of 638 nm, measuring FL6 660/20 nm fluorescence emission for LDH, and with a blue excitation laser of 488 nm, measuring FL1 525/BP nm fluorescence emission for MCT1. We excluded aggregates by gating single cells according to their area and peak fluorescence signal. As a negative control to quantify non-specific antibody background signal, we used Alexa Fluor™ 647 Rabbit IgG Isotype Control (BD Biosciences) and Mouse IgG2A Alexa Fluor™ 488-conjugated Antibody at same concentrations. For LDH and MCT1 fluorescence to concentration conversions, we used AccuCheck ERF reference particles (ThermoFisher) as a calibration curve. Then, we calculated the Degree of Labelling (DOL) to determine the protein and fluorophore molar concentrations of the conjugated using absorbance measurements.

### 4.7 Hyperpolarisation of [1-^13^C]pyruvate via dissolution Dynamic Nuclear Polarisation

We prepared our metabolic substrate from 1-^13^C labelled pyruvic acid (14.21 M, Merck Life Sciences), trityl OX063 radical (18.085 mM, GE Healthcare) and Gd-DOTA (1.43 mM, Guerbet). We enhanced the sample’s NMR signal via dissolution Dynamic Nuclear Polarisation (dDNP) with a HyperSense polariser (Oxford Instruments Plc.). This technique increases the polarisation level of ^13^C nuclei within our pyruvic acid solution in a static magnetic field (more details in Suppl. Section C). After polarising 24 *µ*L (31.01 ±0.89 mg) of [1-^13^C]pyruvate, the dDNP machine dissolved it in 4.73 ± 0.33 mL of a heated buffer solution (40.04 mM HEPES, 94.00 mM NaOH, 0.34 mM EDTA-H4, 3.00 mM NaCl in MilliQ water) adjusted by pyruvate weight (Suppl. Section C). The ejected buffered sample (pH ≈ 7.4) was at 80 mM, which we further diluted to the desired final concentration for each experiment. Samples retained ≈ 12% polarisation at the first data acquisition point.

### 4.8 Hyperpolarised nuclear magnetic resonance data acquisition

We acquired all spectroscopic data in a 1.4 T benchtop NMR spectrometer (Pulsar, Oxford Instruments Ltd.) with an internal probe temperature of 37 ^°^C. We ensured that all samples were within the ≈ 12.5mm-long effective detection region (250 *µ*L) of the spectrometer by using 5mm diameter NMR tubes with PDMS droplets at the bottom [16]. We acquired our spectroscopic data with a 15^°^ flip angle using a ^13^C free induction decay (FID) pulse sequence with WALTZ-4 ^1^H channel decoupling and receiver attenuation to 0.78 dB. Each acquisition consisted of 80 scans with a 5 second repetition time (TR). We obtained spectroscopic data for at least *n* = 3 replicates for each experimental condition.

### 4.9 Hyperpolarised nuclear magnetic resonance spectroscopic data processing

We processed all our spectroscopic data (i.e., 1 Hz exponential apodisation, phase and baseline correction) using MestreNova (Mestrelab Research, v. 14.2.0). Then, we integrated the [1-^13^C]pyruvate (≈ 171 ppm) and [1-^13^C]lactate (≈ 183 ppm) peaks for metabolite quantification over time. Next, we computed the mean and standard deviations for each sample of each triplicate experiment. We developed custom Julia code for further analysis, such as *T*_1_ fittings or pyruvate signal extrapolation at the time of mixing with cells (*Pyr*_0_), which occurred ≈ 25 − 30 s before the first scan due to sample handling. For simplicity, all *T*_1_s estimated, reported and used are apparent, and hence the data is uncorrected for pulsing effects.

### 4.10 Mathematical models development

We developed all our deterministic models of the form *d*[*x*]*/dt* = *f* (*x, θ, t*) following mass action kinetics, where [*x*] is the concentration of the different state variables at a given time *t*, and *θ* is the set of parameters. First, we retrieved from literature a commonly used [18] mathematical model ℳ (*θ*_*α*_) (Fig. 2e blue, Suppl. Eq. 1) describing the conversion of pyruvate to lactate (and vice-versa) produced in the cells plus the signal decay due to longitudinal *T*_1_ relaxation of the hyperpolarised molecules. From this model, we derived ℳ_*T*_ (*θ*_*β*_) (Fig. 2e red, Suppl. Eq. 2) including a term for pyruvate transport from the extracellular medium to the cytoplasm through monocarboxylate transporters (MCT). Next, we expanded the previous model to include Michaelis-Menten kinetics (Fig. 3d, Suppl. Eq. 3). Finally, we developed a larger model ℳ_*CR*_(*θ*_*γ*_) (Fig. 3e, Suppl. Eq. 6), including transport and more complex enzymatic mechanisms from the lactate dehydrogenase (LDH), such as competitive repression between substrate (pyruvate) and co-enzyme (NADH). For this model, we considered the initial concentration of *NADH*_0_, *NAD*+_0_ and *LDH*_0_ as additional model parameters. We checked all models for potential structural unidentifiabilities (absent) using the Matlab toolbox STRIKE-GOLDD [44].

### 4.11 Mathematical models simulations

We developed all simulation code in Julia programming language (V 1.9.4) in a computer with an Intel(R) Core(TM) i7-10700 CPU with 16 logical processors (2.90GHz) and 32 GB of RAM. For all numerical predictions, we used non-stiff solvers, these being CVODE with backward differentiation formula (BDF) for both Julia and Stan. We used the same methods to simulate the models to generate pseudo-data applying heteroscedastic noise as observed in experimental data, with their proportions estimated experimentally. For pyruvate, the simulation remained as the mean vector with a noise of 12 % of this for *σ*. Lactate followed the form 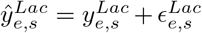 where *ŷ* is the pseudo-data, *y* the model simulation, *s* the sampling time, *e* the experimental design and *ϵ* the additive heteroscedastic Gaussian noise equal to 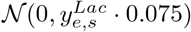.

### 4.12 Bayesian parameter inference on time-series data

For each model considered, we used Bayesian statistics to infer the posterior probabilities for all the rate constants *θ*_*i*_ ∈ ℝ^*k*^, with *k* = 4, 5, 11 and *i* = *α, β, γ* and three initial state conditions *Y*_0_ ∈ ℝ^*j*^, with *j* = 3 (*NADH*_0_, *NAD*+_0_ and *LDH*_0_), for model ℳ _*CR*_(*θ*_*γ*_). For this larger model, we assumed all the rest of the initial state conditions to be zero except for the initial exogenous pyruvate added *Pyr*_*OUT*_, which is set experimentally. Note that we also fixed additional model parameters 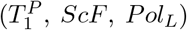 to experimentally estimated values to reduce the complexity of the problem (see Suppl. Table 5).

Bayes’ theorem states that the posterior probability *P* (*θ* | *D*) equals the likelihood *P* (*D* | *θ*) times the prior distribution *P* (*θ*) divided by the marginal likelihood of the data *P* (*D*), giving the formula:

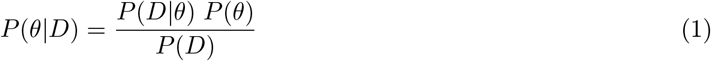

To sample the target posterior distributions, we used Stan [45] embedded in Julia. Stan uses the No-U-Turn Sampler (NUTS), an algorithm with a low need for user hyper-parameter tuning that leverages the proportionality:

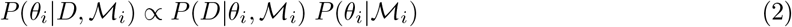

to approximate the model posteriors through Markov-Chain Monte Carlo (MCMC) algorithm. For all inferences, we assumed a normally distributed likelihood *P* (*D* | *θ*), drawing 8000 samples from the equilibrium target posterior after 400-1000 warmup iterations and dividing the task between four to ten parallel Markov chains. We started with highly uninformative priors derived from initial maximum likelihood estimation (MLE) explorations within biologically relevant bounds. All parameter priors, bounds, descriptions and sets used to generate pseudo-data are in Suppl. Table 1 for ℳ (*θ*_*α*_), Suppl. Table 3 for ℳ_*T*_ (*θ*_*β*_) and Suppl. Tables 5, 6 for ℳ_*CR*_(*θ*_*γ*_). For initial quick parameter estimates and to compute good initial guesses for the MCMCs, we solved the non-linear optimisation problem of minimising the MLE formula:

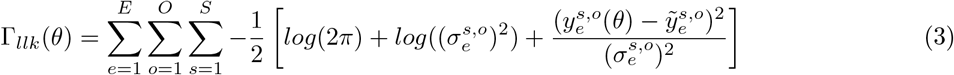

where 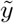 is the mean experimental data with standard deviation *σ, y*(*θ*) the model simulations, *E* the total number of experiments, *O* the total number of observables, *S* the total number of samples. To tackle the non-convexity and high complexity of the problem, we used the meta-heuristic genetic algorithm optimiser differential evolution.

### 4.13 Model comparison metrics

To compare the prediction quality of different models ℳ_*i*_ for the experimental data 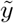 of a given experiment, we used the Bayesian information criteria (BIC):

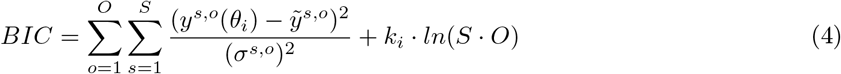

where *k*_*i*_ is the dimensionality of the parameter vector *θ*_*i*_, randomly sampled from the posterior *P* (*θ*_*i*_ | *D*) for a given model. Hence, for all results shown, we computed the BIC metric with the simulation *y*(*θ*_*i*_) using all the 8000 posterior samples obtained from Stan’s Bayesian inference. To corroborate the robustness of the assessment, in the same fashion, we included as a second metric a modified version of the normalised root-mean-square error (nRMSE), including and Occam’s factor to penalise model complexity:

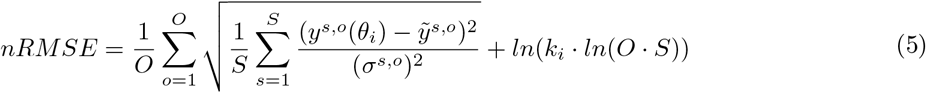

### 4.14 Model prediction uncertainty evaluation

In this work, we propose a novel approach to assess the informative content of experiments before performing these in a Bayesian setting. Instead of being based on mutual information, we propose to evaluate the informativeness of experiments based on the level of prior prediction uncertainty. The higher the prior prediction uncertainty, the more informative an experiment poses since the model has more to learn from the data thanks to the direct relationship between the model prediction *h*(*θ*) and the prior knowledge *P* (*θ*) for parameters. Hence, the prior prediction uncertainty for a particular experimental design *e*_*d*_ can be defined as:

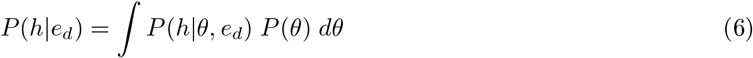

To approximate *P* (*h* | *e*_*d*_), we introduce a sample-based method Λ^*E*^ that for an experiment *e* and each model observable *O* computes the Shannon entropy [46] of the model prediction for each sampling time *S* as:

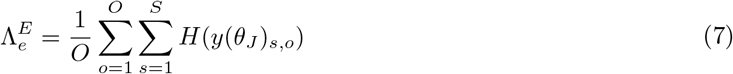

Note that *y*(*θ*_*J*_) are the model simulations for all the *j* randomly selected samples drawn from the prior distribution *P* (*θ*). An alternative approximation of this prediction uncertainty is the Euclidean distance between percentiles calculated on all the model simulations [47]. Yet, this approach makes the metric unsusceptible to the shape of the prediction uncertainty, assuming general normality.

### 4.15 Bayesian optimal experimental design (bOED)

With our novel approach, the problem of optimally designing experiments using a Bayesian framework is treated as a non-linear optimisation problem to find an input *u* that maximises Λ^*E*^. In our context, the input variable *u* is the initial concentration of exogenous [1-^13^C]pyruvate provided to cells. We bounded the pyruvate concentration in the form *u*^*L*^ *< u < u*^*U*^ where *u*^*L*^ = 0.1 mM and *u*^*U*^ = 40 mM following all the standard experimental conditions defined previously (i.e., experimental times and sampling points). Due to experimental limitations, we discretised the range of pyruvate concentrations to one decimal point. Hence, the algorithm calculated Λ^*E*^ using the prior prediction of a complete experiment considering only [1-^13^C]lactate (i.e., the product of the reaction), since for the total [1-^13^C]pyruvate the only prediction uncertainty affecting parameter was 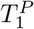, which was fixed experimentally.

To efficiently solve the non-linear optimisation problem, we leveraged Bayesian optimisation (BayesOpt) algorithms. We used the default parameters with an ARD squared exponential covariance kernel and the expected improvement sampling algorithm with a maximum of 500 iterations. While the total number of allowed pyruvate concentrations was 400, we implemented the BayesOpt approach to be extrapolated to more complex scenarios, both discrete and continuous. Furthermore, since we had to discretise the inputs selected, we rounded the input specified by BayesOpt, resulting in a scenario where one value could be chosen more than once, yet without affecting the optimisation scheme since it is not a gradient-based algorithm. To speed up the process, we down-sampled the priors to 1000 total random samples.

### 4.16 Random and intuition-derived experimental design

As a more traditional approach to designing our type of experiments, we selected two approaches, a random one (Ran) and a design based on user decisions based on acquired information, which we called intuition-derived (ID). For the random experiments, we selected a random value for the administered [1-^13^C]pyruvate concentration between the allowed concentration bounds following the uniform distribution *U*(0.1, 40), again discretised at the first decimal point. For the intuition-derived design, first, we computed the average vector value for the prior distribution used. Then, we simulated the system for each pyruvate concentration allowed and computed the total [1-^13^C]lactate signal. By observing the relation between input (pyruvate) and output (lactate) we decided in each case the point in the curve that would be, intuitively, more informative to observe (e.g., maximum point before repression or a point halfway between maximum and minimum production).

### 4.17 Assessment of *a posteriori* information content of experiments

To assess the quality of the diverse posteriors inferred, particularly when comparing experimental design strategies, we selected two information theory-based metrics for all studies and a distance metric for computational ones. For the *in silico* validation of the OED methodology proposed, we introduced the posterior sample distance metric *d*_*ϕ*_:

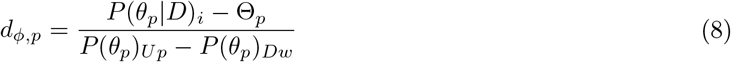

In this proposed metric, *P* (*θ* | *D*)_*i*_ is the *i*th random sample for a parameter *p* of the inferred posterior, Θ the parameter value selected to generate the pseudo-data and *P* (*θ*)_*Up*_ and *P* (*θ*)_*Dw*_ the upper and lower allowed bounds for the priors as a normalisation factor. As a summarising metric of this distance, we also computed the absolute average 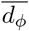 as:

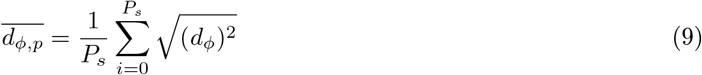

where *P*_*s*_ is the total number of samples drawn from the posterior.

As an information theory-based metric, we selected the gain in information *G*_*i*_ defined as the difference between the prior and posterior Shannon entropy:

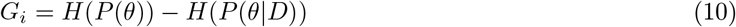

where the entropy of a probability *P* (*X*) is defined as:

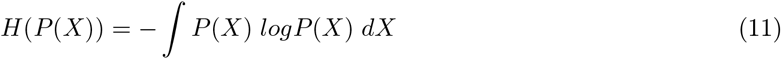

For the starting prior distributions, since we assumed parameter independence 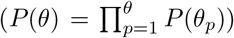, we computed their entropies analytically. For the sampled posteriors, we approximated their Shannon entropy by fitting them to a multidimensional Gaussian mixture model and estimating their entropy [25]. As a quick-to-compute metric to approximate the gain in information, we also used the negative of the natural logarithm of the determinant of the posterior covariance matrix (− *log* |Σ|). This second metric assumes normality of the posterior, and it is derived from *G*_*i*_ by removing all the constant terms of the multivariate Gaussian distribution entropy formulation and the entropy of the prior.

### 4.18 Reporting summary

Further information on research design is available in the Nature Portfolio Reporting Summary linked to this article.

## 5 Data availability

All experimental(raw, processed and analysed) and computational data is available at the Dataverse CORA repository doi.org/10.34810/data3138. This data includes (but is not limited) the source data used to generate all figures and supplementary figures provided in this manuscript.

## 6 Code availability

All the code generated for this study, together with any computational data generated (e.g. pseudo-data for *in silico* studies, simulations or optimisation results) is available at the GitHub repository MM_bOED_HP-NMR.

## Acknowledgements

The authors would like to thank William Mander for his assistance and technical support with the HyperSense polariser. We thank Peter Christian Sperling for his help in cell culturing and biological work. We thank the Scientific and Technological Centers (CCiTUB), Universitat de Barcelona, and staff Jaume Comas for their support and advice on flow cytometry technique. We thank the MicroFabSpace and Microscopy Characterisation Facility, Unit 7 of ICTS “NANBIOSIS” from CIBER-BBN at IBEC.

This work has received funding from: the European Union (GA-101195272, Q-AID) and (GA-101291716, CAMP); a European Union ERC Starting Grant (GA-101165045, LIFETIME); the Spanish grants with reference PID2023-151470OB-I00 funded by MICIU/AEI/ 10.13039/501100011033 and by “ERDF/EU” (METACHIP), RYC2020-029099-I funded by MCIN/AEI10.13039/501100011033 and by “ESF Investing in your future”, PLEC2022-009256 funded by MCIN/AEI/10.13039/501100011033 and by the European Union NextGenerationEU/PRTR (FLASHonCHIP), a grant within the framework Biotechnology Plan Applied to Health funded by MCIN/AEI10.13039/501100011033 and co-financed by the Spanish Ministry of Science and Innovation with funds from the European Union NextGenerationEU from the Recovery, Transformation and Resilience Plan (PRTR-C17.I1); from the Autonomous Community of Catalonia within the framework of the Biotechnology Plan Applied to Health; the BIST (Barcelona Institute of Science and Technology)-”la Caixa” Banking Foundation Chemical Biology programme. Views and opinions expressed are however those of the author only and do not necessarily reflect those of the European Union or the European Research Council. Neither the European Union nor the granting authority can be held responsible for them.

## Author contributions

D.G.C. devised the overall study, developed the computational framework, performed the data analysis, mathematical modelling, script development, and conceptualised and implemented the Bayesian optimal experimental design approach. L.M.F, A.G. and D.G.C. performed cell culturing and preparation for all biological experiments. D.G.C. and G.M. handled the dDNP machine for all HP-NMR experiments. D.G.C. prepared the initial manuscript draft. D.G.C., G.M., L.M.F. and I.M.R. contributed to writing, reviewing and editing the manuscript. I.M.R. conceived the original study, supervised the project, provided scientific guidance and resources, and raised funding.

## Declarations

The authors declare no competing interests.

