## Supplementary Data for "A Bayesian Study to Optimally Unveil LDH Kinetic Mechanisms *In Vivo*"

### A Mathematical Models Derivation

The section contains the mathematical formulations for the three models used in this work, plus information on the model parameters, bounds and priors selected and values selected to generate pseudo-data. The first two models (i.e., $\mathcal{M}(\theta_\alpha)$ and $\mathcal{M}_T(\theta_\beta)$) are mostly designed to describe hyperpolarised nuclear magnetic resonance (HP-NMR) experiments. As commonly used, we introduce a third model following traditional Michaelis-Menten kinetics ($\mathcal{M}_{MM}(\theta_\delta)$). At the same time, the fourth model (i.e., $\mathcal{M}_{CR}(\theta_\gamma)$) is a more general and biologically relevant one that incorporates HP-NMR signal decays as an addition.

#### A.1 Traditional [1-$^{13}C$]pyruvate hyperpolarisation model ($\mathcal{M}(\theta_\alpha)$)

Model $\mathcal{M}(\theta_\alpha)$ is the simplest model used, considering only a direct bidirectional conversion between pyruvate and lactate plus hyperpolarised signal decay due to their longitudinal relaxation times (Fig. ??b, blue, main manuscript):

$$\begin{aligned}\frac{d[Pyr]}{dt} &= k_{LP}[Lac] - (\frac{1}{T_1^P} + k_{PL})[Pyr] \\ \frac{d[Lac]}{dt} &= k_{PL}[Pyr] - (\frac{1}{T_1^L} + k_{LP})[Lac]\end{aligned} \tag{1}$$

Hence, the only model state variables are the concentration of hyperpolarised [1-$^{13}C$]pyruvate ($[Pyr]$) and [1-$^{13}C$]lactate ($[Lac]$), simplifying at the maximum the biological and biochemical components of the system. We used this model since it is a consensus one used in the HP-NMR community [1–5]. Note that, for simplicity (particularly in the following models), we did not correct signals by the hyperpolarisation decay caused by measurement (i.e., pulsing). Hence, all $T_1$s reported are apparent, and the parameter encapsulates the corrective term $\frac{1}{T_1^r} - (1 - cos(\phi))^{1/TR}$ (where $T_1^r$ is the real longitudinal relaxation decay), making the comparison possible since we used the same pulse angle $\phi$ (15°) and repetition time $TR$ (5 $sec$) for all experiments.

##### A.1.1 Model $\mathcal{M}(\theta_\alpha)$ priors

We considered the biological kinetic rates of this model ($k_{PL}$ and $k_{LP}$) to be for a total of one million HepG2 cells following the linear relationship between cell number and signal measured for this model.

**Suppl. Table 1**: Model $\mathcal{M}(\theta_\alpha)$ parameters with their definition, and bounds and priors used in this work.

| Parameter | Description | Bounds | Prior |
|---|---|---|---|
| $k_{PL}$ | Constant rate of pyruvate conversion to lactate (s$^{-1}$) | $[0, 10^{-2}]$ | $\mathcal{U}(1.68 \cdot 1-^{-5}, 1.68 \cdot 1-^{-3})$ |
| $k_{LP}$ | Constant rate of lactate back-conversion to pyruvate (s$^{-1}$) | $[0, 10^{-2}]$ | $\mathcal{U}(2.23 \cdot 1-^{-8}, 2.23 \cdot 1-^{-4})$ |
| $T_1^P$ | Apparent (no pulse corrected) pyruvate longitudinal relaxation time at 1.4 T (s) | $[30, 63]$ | $\mathcal{N}(50, 5)$ |
| $T_1^L$ | Apparent (no pulse corrected) lactate longitudinal relaxation time at 1.4 T (s) | $[20, 100]$ | $\mathcal{N}(55, 10.5)$ |

##### A.2 [1-$^{13}C$]pyruvate hyperpolarisation model with MCT pyruvate transport ($\mathcal{M}_T(\theta_\beta)$)

To make $\mathcal{M}(\theta_\alpha)$ more biologically relevant, we developed model $\mathcal{M}_T(\theta_\beta)$, which includes a simple term for pyruvate transport from the extracellular cell media to the cytosol (Fig. ??b, red, main manuscript) [6]. As previously stated, all $T_1$s reported are apparent. Publications in the field of HP-NMR already stressed the importance of the pyruvate transport by the MCT transporters and their rate limitations in the whole reaction [3, 7, 8].

$$
\begin{aligned}
\frac{d[Pyr_{OUT}]}{dt} &= -k_{in}[Pyr_{OUT}] - \frac{[Pyr_{OUT}]}{T_1^P} \\
\frac{d[Pyr_{IN}]}{dt} &= k_{in}[Pyr_{OUT}] + k_{LP}[Lac] - (\frac{1}{T_1^P} + k_{PL})[Pyr_{IN}] \\
\frac{d[Lac]}{dt} &= k_{PL}[Pyr_{IN}] - (\frac{1}{T_1^L} + k_{LP})[Lac]
\end{aligned}
\tag{2}
$$

Despite the complexity of the biochemical reactions involved in the transport of a molecule of pyruvate across the MCT transporter [9], we opted for a simplified approach coupled to model $\mathcal{M}(\theta_\alpha)$. This approach incorporates two types of hyperpolarised [1-$^{13}C$]pyruvate molecules, the ones outside the cell ($[Pyr_{OUT}]$) and inside ($[Pyr_{IN}]$). Here, we assume that all pyruvate molecules that enter the cells at a constant rate of $k_{in}$ do not get transported back to the extracellular media (if they do, it is once converted to lactate, but we cannot measure that). We can derive this transport term of the model from the original transport reaction:

$$\mathrm{MCT} + \mathrm{H}^+_{\mathrm{out}} \underset{k_1^u}{\overset{k_1^b}{\rightleftharpoons}} \mathrm{MCT{-}H^+} + \mathrm{Pyr_{OUT}} \underset{k_2^u}{\overset{k_2^b}{\rightleftharpoons}} \mathrm{MCT{-}H^+{-}Pyr} \underset{k_1^R}{\overset{k_1^T}{\rightleftharpoons}} \mathrm{MCT{-}H^+} + \mathrm{Pyr_{IN}} \underset{k_2^R}{\overset{k_2^T}{\rightleftharpoons}} \mathrm{MCT} + \mathrm{H}^+_{\mathrm{in}}$$

setting the reaction for only inside transport ($k_1^u, k_2^u, k_1^R, k_2^R = 0$ since $k_1^b >> k_1^u, k_2^b >> k_2^u, k_1^T >> k_1^R, k_2^T >> k_2^R$) and assuming a quasi-steady-state approximation with constant pH balance ($d[H^+_{out}]/dt,\ d[H^+_{in}]/dt = 0$) and MCT species ($d[MCT]/dt,\ d[MCT{-}H^+]/dt,\ d[MCT{-}H^H{-}Pyr]/dt = 0$). Applying these assumptions result in the term $k_2^b \cdot [MCT - H^+] \cdot [Pyr_{OUT}]$, where we define $k_{in} = k_2^b \cdot [MCT - H^+]$.

###### A.2.1 Model $\mathcal{M}_T(\theta_\beta)$ priors

We considered the biological kinetic rates of this model ($k_{in}$, $k_{PL}$ and $k_{LP}$) to be for a total of one million HepG2 cells following the linear relationship between cell number and signal measured for this model.

**Suppl. Table 2**: Model $\mathcal{M}_T(\theta_\beta)$ parameters with their definition, and bounds and priors used in this work.

| Parameter | Description | Bounds | Prior |
|---|---|---|---|
| $k_{in}$ | Pyruvate transport rate from extracellular media to cytosol via MCT transporters (s$^{-1}$) | $[0, 10^{-1}]$ | $\mathcal{U}(4.896 \cdot 1-^{-5}, 4.896 \cdot 1-^{-3})$ |
| $k_{PL}$ | Constant rate of pyruvate conversion to lactate (s$^{-1}$) | $[0, 10^{-2}]$ | $\mathcal{U}(9.999 \cdot 1-^{-4}, 9.999 \cdot 1-^{-2})$ |
| $k_{LP}$ | Constant rate of lactate back-conversion to pyruvate (s$^{-1}$) | $[0, 10^{-4}]$ | $\mathcal{U}(7.89 \cdot 1-^{-8}, 7.89 \cdot 1-^{-2})$ |
| $T_1^P$ | Apparent (no pulse corrected) pyruvate longitudinal relaxation time at 1.4 T (s) | $[30, 63]$ | $\mathcal{N}(51.21, 5.12)$ |
| $T_1^L$ | Apparent (no pulse corrected) lactate longitudinal relaxation time at 1.4 T (s) | $[20, 100]$ | $\mathcal{N}(41.73, 8.34)$ |

#### A.3 [1-$^{13}C$]pyruvate hyperpolarisation model considering Michaelis-Menten kinetics with pyruvate transport ($\mathcal{M}_{MM}(\theta_\delta)$)

As an alternative to introduce common enzymatic kinetics to the model, we developed $\mathcal{M}_{MM}(\theta_\delta)$ following simple Michaelis-Menten kinetics. Besides the pyruvate transport across the cell membrane, the model considers a step for the binding of pyruvate to the LDH enzyme before it gets metabolised to lactate. This model introduces substrate production saturation, a common and observable phenomenon in biological systems not taken into account for models $\mathcal{M}(\theta_\alpha)$ and $\mathcal{M}_T(\theta_\beta)$. For simplicity reasons, and as for $\mathcal{M}_{CR}(\theta_\gamma)$, we removed the backwards reaction where lactate gets converted to pyruvate. As for the previous models, all $T_1$s reported are apparent.

Reaction:

$$\mathsf{Pyr_{OUT}} \xrightarrow{\mathsf{k_{in}}} \mathsf{Pyr_{IN}} + \mathsf{LDH} \underset{\mathsf{k}_r}{\overset{\mathsf{k}_f}{\rightleftharpoons}} \mathsf{LDH{-}Pyr} \xrightarrow{\mathsf{k_{PL}}} \mathsf{LDH} + \mathsf{Lac}$$

$$\downarrow T_1^P \qquad \downarrow T_1^P \qquad \downarrow T_1^P \qquad \downarrow T_1^L$$
$$\emptyset \qquad \emptyset \qquad \emptyset \qquad \emptyset$$

Ordinary Differential Equations:

$$\begin{aligned}
\frac{d[Pyr_{OUT}]}{dt} &= -k_{in}[Pyr_{OUT}] - \frac{[Pyr_{OUT}]}{T_1^P} \\
\frac{d[Pyr_{IN}]}{dt} &= k_{in}[Pyr_{OUT}] + k_r[LDH - Pyr] - k_f[Pyr_{IN}][LDH] - \frac{[Pyr_{IN}]}{T_1^P} \\
\frac{d[LDH]}{dt} &= k_r[LDH - Pyr] + k_{PL}[LDH - Pyr] - k_f[Pyr_{IN}][LDH] \\
\frac{d[LDH - Pyr]}{dt} &= k_f[Pyr_{IN}][LDH] - k_r[LDH - Pyr] - k_{PL}[LDH - Pyr] - \frac{[LDH - Pyr]}{T_1^P} \\
\frac{d[Lac]}{dt} &= k_{PL}[LDH - Pyr] - \frac{Lac}{T_1^L}
\end{aligned} \tag{3}$$

##### A.3.1 Model $\mathcal{M}_{MM}(\theta_\delta)$ priors

We considered the biological kinetic rates of this model ($k_{in}$) and total enzyme concentration ($[LDH]_T$) to be for a total of four million HepG2 cells following the linear relationship between cell number and signal measured for this model.

**Suppl. Table 3**: Model $\mathcal{M}_{MM}(\theta_\beta)$ parameters and total LDH enzyme with their definition, and bounds and priors used in this work.

| Parameter | Description | Bounds | Prior |
|---|---|---|---|
| $k_{in}$ | Pyruvate transport rate from extracellular media to cytosol via MCT transporters (s$^{-1}$) | $[0, 1]$ | $\mathcal{U}(0, 0.2)$ |
| $k_f$ | Constant rate for pyruvate binding to LDH (s$^{-1}$) | $[0, 1]$ | $\mathcal{U}(0, 0.5)$ |
| $k_r$ | Constant rate for pyruvate unbinding to LDH (s$^{-1}$) | $[0, 1.1]$ | $\mathcal{U}(0, 1.1)$ |
| $k_{PL}$ | Constant rate of pyruvate conversion to lactate (s$^{-1}$) | $[0, 1]$ | $\mathcal{U}(0, 0.4)$ |
| $T_1^P$ | Apparent (no pulse corrected) pyruvate longitudinal relaxation time at 1.4 T (s) | - | 51 |
| $T_1^L$ | Apparent (no pulse corrected) lactate longitudinal relaxation time at 1.4 (s) | - | 41 |
| $[LDH]_T$ | Total LDH enzyme cellular sample concentration (assumed to be equal to $[LDH]$ at the beginning of experiment) ($\mu$M) | $[0, 1600.5]$ | $\mathcal{N}(550, 275)$ |

##### A.4 [1-$^{13}C$]pyruvate hyperpolarisation model with MCT pyruvate transport and LDH competitive repression between pyruvate and NADH ($\mathcal{M}_{CR}(\theta_\gamma)$)

Although models $\mathcal{M}(\theta_\alpha)$ and $\mathcal{M}_T(\theta_\beta)$ can describe the system's behaviour for a specific [1-$^{13}C$]pyruvate concentration, both break when attempting to generalise it. The reason for the failure of their simplicity resides in the fact that such models fail to describe the decreasing lactate production at high pyruvate concentrations, observed since the 60s and was observed to be NADH dependent [10–13]. While some hypotheses have arisen on the biochemical mechanism of the system, such as allosteric and sequential binding mechanisms [14], we adopted a long-hypothesised model postulating competitive repression between NADH (co-factor) and pyruvate [15]. With this model ($\mathcal{M}_{CR}(\theta_\gamma)$), first NADH is required to bind to LDH (lactate dehydrogenase), followed by pyruvate for the reaction to happen in a Michaelis-Menten fashion. If a pyruvate molecule of binds before the co-factor, the enzyme becomes un-catalytic until this unbinds to allow NADH to enter the catalytic centre (Fig. **??**e). The process is in accordance per the structural analysis of the catalytic site of the enzyme [16, 17]. Furthermore, preliminary tests with diverse models pointed out this one as the best predictive. For simplicity, we only considered the conversion from pyruvate to lactate and removed the backward conversion. Such assumption is in concordance with low $k_{LP}$ values obtained in the previous models and the fact that in hepatic cells used (i.e. HepG2), the izoenzyme 5 is the predominant one [18, 19], which favours the production of lactate against the backwards reaction [20].

Reaction:

$$
\begin{array}{ccccccccc}
 & & \mathsf{Pyr_{OUT}} & & & & \mathsf{Pyr_{OUT}} & & \\
 & & \downarrow k_{in} & & & & \downarrow k_{in} & & \\
\mathsf{LDH_{Inact}} & \underset{\mathsf{k}_i}{\overset{\mathsf{k}_a}{\rightleftharpoons}} & \mathsf{Pyr_{IN}} + \mathsf{LDH} + \mathsf{NADH} & \underset{\mathsf{k}_u{}^{\mathsf{N}}}{\overset{\mathsf{k}_b{}^{\mathsf{N}}}{\rightleftharpoons}} & \mathsf{LDH^{N}} + & & \mathsf{Pyr_{IN}} & \underset{\mathsf{k}_u{}^{\mathsf{P}}}{\overset{\mathsf{k}_b{}^{\mathsf{P}}}{\rightleftharpoons}} & \mathsf{LDH^{N-P}} \xrightarrow{\mathsf{k_{PL}}} \mathsf{LDH^{N+}} + \mathsf{Lac} \underset{\mathsf{k}_u{}^{\mathsf{N+}}}{\overset{\mathsf{k}_b{}^{\mathsf{N+}}}{\rightleftharpoons}} \mathsf{LDH} + \mathsf{NAD^{+}} \\
\downarrow T_1^P & & \downarrow T_1^P & & & & \downarrow T_1^P & & \downarrow T_1^P \qquad \downarrow T_1^L \\
\emptyset & & \emptyset & & & & \emptyset & & \emptyset \qquad \emptyset
\end{array}
$$

Ordinary Differential Equations:

$$
\begin{aligned}
\frac{d[Pyr_{OUT}]}{dt} &= -k_{in}[Pyr_{OUT}]\\
\frac{d[Pyr_{IN}]}{dt} &= k_{in}[Pyr_{OUT}] + k_u^P[LDH^{N-P}] + k_a[LDH_{Inact}] - k_b^P[LDH^N][Pyr_{IN}] - k_i[LDH][Pyr_{IN}]\\
\frac{d[NADH]}{dt} &= k_u^N[LDH^N] - k_b^N[LDH][NADH]\\
\frac{d[NAD+]}{dt} &= k_u^{N+}[LDH^{N+}] - k_b^{N+}[LDH][NAD+]\\
\frac{d[LDH]}{dt} &= k_u^N[LDH^N] + k_a[LDH_{Inact}] + k_u^{N+}[LDH^{N+}] - k_b^N[LDH][NADH] - k_i[LDH][Pyr_{IN}]\\
&\quad - k_b^{N+}[LDH][NAD+]\\
\frac{d[LDH^N]}{dt} &= k_b^N[LDH][NADH] + k_u^P[LDH^{N-P}] - k_u^N[LDH^N] - k_b^P[LDH^N][Pyr_{IN}]\\
\frac{d[LDH^{N+}]}{dt} &= k_{PL}[LDH^{N-P}] + k_b^{N+}[LDH][NAD+] - k_u^{N+}[LDH^{N+}]\\
\frac{d[LDH^{N-P}]}{dt} &= k_b^P[LDH^N][Pyr_{IN}] - k_u^P[LDH^{N-P}] - k_{PL}[LDH^{N-P}] \qquad (4)\\
\frac{d[LDH_{Inact}]}{dt} &= k_i[LDH][Pyr_{IN}] - k_a[LDH_{Inact}]\\
\frac{d[Lac]}{dt} &= k_{PL}[LDH^{N-P}]
\end{aligned}
$$

$$
\begin{aligned}
\frac{d[Pyr_{HP}]}{dt} &= \left(\frac{d[Pyr_{OUT}]}{dt} + \frac{d[Pyr_{IN}]}{dt} + \frac{d[LDH_{Inact}]}{dt} + \frac{d[LDH^{N-P}]}{dt}\right) - \frac{[Pyr_{HP}]}{T_1^P}\\
\frac{d[Lac_{HP}]}{dt} &= \frac{d[Lac]}{dt} - \frac{[Lac_{HP}]}{T_1^L}
\end{aligned}
$$

$$
\begin{aligned}
\frac{d[Pyr_{HP}^{Obs}]}{dt} &= \frac{d[Pyr_{HP}]}{dt}(ScF \cdot Pol_L)\\
\frac{d[Lac_{HP}^{Obs}]}{dt} &= \frac{d[Lac_{HP}]}{dt}(ScF \cdot Pol_L)
\end{aligned}
$$

The model also includes the pyruvate transport term of $\mathcal{M}_T(\theta_\beta)$. The model also has additional equations to convert from the biological signal to the hyperpolarised signal ($d[Pyr_{HP}]/dt$ and $d[Lac_{HP}]/dt$) and conversion factors to account for polarisation levels ($Pol_L$) and conversion between $mM$ to $A.U.$ ($ScF$) to generate the observable pyruvate ($d[Pyr_{HP}^{Obs}]/dt$) and lactate ($d[Lac_{HP}^{Obs}]/dt$) signals. As previously stated, all $T_1$s reported are apparent.

##### A.4.1 Model $\mathcal{M}_{CR}(\theta_\gamma)$ priors

We considered the biological kinetic rates of this model ($k_{in}$) and involved states to be for a total of three million HepG2 cells following the linear relationship between cell number and signal measured for the transport, and amount of enzyme ($LDH_{Tot} = LDH + LDH^N + LDH^{N+} + LDH^{N-P} + LDH_{Inact}$) and coenzyme ($NAD_{Tot} = NADH + NAD+ + LDH^N + LDH^{N+} + LDH^{N-P}$). We used Bounds$_1$, Priors$_1$ and $\Theta_1$ for the computational validation of the novel Bayesian optimal experimental design algorithm (bOED), Bounds$_2$ and Prior$_2$ for the experimental validation of the novel bOED algorithm and Prior$_3$ and $\Theta_2$ for the computational study on the effect of data noise in posterior uncertainty.

**Suppl. Table 4**: Model $\mathcal{M}_{CR}(\theta_\gamma)$ parameters. Kinetic rates are represented as $s^{-1}$, $T_1$s as s, $ScF$ unitless and $Pol_L$ as percentage.

| Parameter | Description | Bounds$_1$ | Bounds$_2$ | Prior$_1$ | Prior$_2$ | $\Theta_1$ |
|---|---|---|---|---|---|---|
| $k_{in}$ | Pyruvate transport rate from extracellular media to cytosol via MCT transporters | $[0.001, 0.01]$ | $[0, 2.75 \cdot 10^{-2}]$ | $\mathcal{N}(5.5 \cdot 10^{-3}, 2.75 \cdot 10^{-3})$ | $\mathcal{N}(5.5 \cdot 10^{-3}, 5.5 \cdot 10^{-3})$ | $5.91 \cdot 10^{-3}$ |
| $k_{PL}$ | Constant rate of pyruvate conversion to lactate | $[0.005, 0.05]$ | $[0, 1.38 \cdot 10^{-1}]$ | $\mathcal{N}(2.75 \cdot 10^{-2}, 1.375 \cdot 10^{-2})$ | $\mathcal{N}(2.75 \cdot 10^{-2}, 2.75 \cdot 10^{-2})$ | $1.83 \cdot 10^{-2}$ |
| $k_b^N$ | Constant rate for the binding of the co-factor NADH to an empty LDH enzyme | $[0.01, 0.08]$ | $[0, 2.25 \cdot 10^{-1}]$ | $\mathcal{N}(4.5 \cdot 10^{-2}, 2.25 \cdot 10^{-2})$ | $\mathcal{N}(4.5 \cdot 10^{-2}, 4.5 \cdot 10^{-2})$ | $4.22 \cdot 10^{-2}$ |
| $k_u^N$ | Constant rate for the un-binding of the co-factor NADH from a molecule of LDH without pyruvate | $[0.1, 0.8]$ | $[0, 2]$ | $\mathcal{N}(4.5 \cdot 10^{-1}, 2.25 \cdot 10^{-1})$ | $\mathcal{N}(4.5 \cdot 10^{-1}, 4.5 \cdot 10^{-1})$ | $5.37 \cdot 10^{-1}$ |
| $k_b^P$ | Constant rate for the binding of a molecule of pyruvate to the LDH enzyme with a previously bound NADH molecule | $[0.7, 1.2]$ | $[0, 2]$ | $\mathcal{N}(9.5 \cdot 10^{-1}, 4.75 \cdot 10^{-1})$ | $\mathcal{N}(9.5 \cdot 10^{-1}, 9.5 \cdot 10^{-1})$ | $9.81 \cdot 10^{-1}$ |
| $k_u^P$ | Constant rate for the un-binding (i.e., no catalytic reaction) of a molecule of pyruvate from the LDH enzyme with a previously bound NADH | $[0.7, 1.2]$ | $[0, 2]$ | $\mathcal{N}(9.5 \cdot 10^{-1}, 4.75 \cdot 10^{-1})$ | $\mathcal{N}(9.5 \cdot 10^{-1}, 9.5 \cdot 10^{-1})$ | $9.91 \cdot 10^{-1}$ |
| $k_u^{N+}$ | Constant rate for the un-binding of a molecule of NAD+ from the enzyme LDH without pyruvate or after the catalytic reaction | $[0.7, 1.2]$ | $[0, 2]$ | $\mathcal{N}(9.5 \cdot 10^{-1}, 4.75 \cdot 10^{-1})$ | $\mathcal{N}(9.5 \cdot 10^{-1}, 9.5 \cdot 10^{-1})$ | $9.33 \cdot 10^{-1}$ |
| $k_b^{N+}$ | Constant rate for the binding of a molecule of NADH+ to an empty LDH enzyme | $[0.005, 0.012]$ | $[0, 4.25 \cdot 10^{-2}]$ | $\mathcal{N}(8.5 \cdot 10^{-3}, 4.25 \cdot 10^{-3})$ | $\mathcal{N}(8.5 \cdot 10^{-3}, 8.5 \cdot 10^{-3})$ | $8.24 \cdot 10^{-3}$ |
| $k_i$ | Constant rate for the binding of pyruvate to an empty LDH enzyme (enzymatic inhibition) | $[0.7, 1.2]$ | $[0, 2]$ | $\mathcal{N}(9.5 \cdot 10^{-1}, 4.75 \cdot 10^{-1})$ | $\mathcal{N}(9.5 \cdot 10^{-1}, 9.5 \cdot 10^{-1})$ | $9.94 \cdot 10^{-1}$ |
| $k_a$ | Constant rate for the un-binding of a pyruvate molecule from the LDH enzyme without co-factor (enzymatic activation) | $[0.0008, 0.005]$ | $[0, 1.45 \cdot 10^{-2}]$ | $\mathcal{N}(2.9 \cdot 10^{-3}, 1.45 \cdot 10^{-3})$ | $\mathcal{N}(2.9 \cdot 10^{-3}, 2.9 \cdot 10^{-3})$ | $1.97 \cdot 10^{-3}$ |
| $T_1^P$ | Apparent (no pulse corrected) pyruvate longitudinal relaxation time at 1.4T | - | - | - | Computed experimentally (51.23) | 55 |
| $T_1^L$ | Apparent (no pulse corrected) lactate longitudinal relaxation time at 1.4T | - | $[0.92, 80.92]$ | - | $\mathcal{N}(40.92, 10)$ | 48 |
| $ScF$ | Scaling factor to convert between concentration in $\mu$M to the A.U. measured in the NMR benchtop | - | - | - | Computed experimentally (-) | 15 |
| $Pol_L$ | Average polarisation level (%) of our HyperSense dDNP | - | - | - | Computed experimentally (12%) | 12 %<br>12% |

**Suppl. Table 5**: Model $\mathcal{M}_{CR}(\theta_\gamma)$ parameters and $y_0$ priors and values used to compute the effect of data noise in posterior uncertainty (Suppl. Fig. 16).

| Parameter | $\text{Prior}_3$ | $\Theta_2$ |
|---|---|---|
| $k_{in}$ | $\mathcal{N}(4.75\cdot10^{-3}, 1\cdot10^{-3})$ | $5.91\cdot10^{-3}$ |
| $k_{PL}$ | $\mathcal{N}(1.375\cdot10^{-2}, 5\cdot10^{-3})$ | $1.83\cdot10^{-2}$ |
| $k_b^N$ | $\mathcal{N}(4.5\cdot10^{-2}, 2.5\cdot10^{-3})$ | $4.22\cdot10^{-2}$ |
| $k_u^N$ | $\mathcal{N}(5.5\cdot10^{-1}, 2.5\cdot10^{-2})$ | $5.37\cdot10^{-1}$ |
| $k_b^P$ | $\mathcal{N}(9.5\cdot10^{-1}, 2\cdot10^{-2})$ | $9.81\cdot10^{-1}$ |
| $k_u^P$ | $\mathcal{N}(9.5\cdot10^{-1}, 2\cdot10^{-2})$ | $9.91\cdot10^{-1}$ |
| $k_u^{N+}$ | $\mathcal{N}(9.5\cdot10^{-1}, 2\cdot10^{-2})$ | $9.33\cdot10^{-1}$ |
| $k_b^{N+}$ | $\mathcal{N}(8.5\cdot10^{-3}, 2.5\cdot10^{-3})$ | $8.24\cdot10^{-4}$ |
| $k_i$ | $\mathcal{N}(9.5\cdot10^{-1}, 3\cdot10^{-2})$ | $9.94\cdot10^{-1}$ |
| $k_a$ | $\mathcal{N}(2\cdot10^{-3}, 1.5\cdot10^{-4})$ | $1.97\cdot10^{-3}$ |
| $T_1^P$ | 55 | 55 |
| $T_1^L$ | 48 | 48 |
| $ScF$ | 15 | 15 |
| $Pol_L$ | 12 % | 12 % |
| $[NADH_0]$ | $\mathcal{N}(650, 260)$ | 450 |
| $[NAD{+}_0]$ | $\mathcal{N}(16000, 6400)$ | 12599 |
| $[LDH_0]$ | $\mathcal{N}(550, 275)$ | 226 |

Additionally, we followed assumptions to define the initial state for the simulations ($y_0$). For pyruvate, $[Pyr_{OUT}]$ contained the total pyruvate signal at $t = 0$, so we set this to the selected concentration used for each experiment. Hence, we set $[Pyr_{IN}]$ and $[Lac]$ to 0. For all intermediary species ($LDH^N$, $LDH^{N+}$, $LDH^{N-P}$, $LDH_{Inact}$), we assume their concentration was 0 at the beginning of the experiment. We computed the corresponding initial concentrations and values for $[Pyr_{HP}]$ and $[Pyr_{HP}^{Obs}]$. Finally, we set up prior distributions for the initial concentrations of $LDH$, $NADH$ and $NAD+$ for 3 million HepG2 cells. We used $\text{Bounds}_\alpha$, $\text{Priors}_\alpha$ and $Y_{0\alpha}$ for the computational validation of the novel Bayesian optimal experimental design algorithm (bOED), and $\text{Bounds}_\beta$, $\text{Priors}_\beta$ for experimental validation of the novel bOED.

**Suppl. Table 6**: Model $\mathcal{M}_{CR}(\theta_\gamma)$ inferred initial states ($Y_0$) with their definition, and bounds and priors, and values used to generate pseudo-data used in this work for three million cells.

| Initial State | Description | $\text{Bounds}_\alpha$ | $\text{Bounds}_\beta$ | $\text{Prior}_\alpha$ | $\text{Prior}_\beta$ | $Y_{0\alpha}$ |
|---|---|---|---|---|---|---|
| $[NADH_0]$ | Initial NADH concentration ($\mu$M) at injection of pyruvate to cells (time 0) | $[300, 1000]$ | $[0, 1600]$ | $\mathcal{N}(650, 260)$ | $\mathcal{N}(650, 260)$ | 450 |
| $[NAD{+}_0]$ | Initial NAD+ concentration ($\mu$M) at injection of pyruvate to cells (time 0) | $[7000, 25000]$ | $[0, 4\cdot10^4]$ | $\mathcal{N}(16000, 6400)$ | $\mathcal{N}(16000, 6400)$ | 12599 |
| $[LDH_0]$ | Initial LDH concentration ($\mu$M) at injection of pyruvate to cells (time 0) | $[100, 1000]$ | $[0, 1600]$ | $\mathcal{N}(550, 275)$ | $\mathcal{N}(550, 275)$ | 226 |

Due to the low identifiability of the initial conditions for NADH, NAD+ and LDH and the aim to make the model $\mathcal{M}_{CR}(\theta_\gamma)$ more accurate to the biological process, we estimated these $Y_0$s experimentally. We estimated the concentrations of NADH and NAD+ using a colourimetric assay and LDH via flow cytometry. For NADH and NAD+, we set up the results as priors since not all the molecules might end up being available to the pathway, assuming a minimum of 10 % of the mean for NADH and a minimum of 1 % of the mean for NAD+. Due to these modifications, we adjusted some parameter priors to avoid chain divergences in the MCMC inference, with the bounds and priors for all parameters and $Y_0$s shown in Suppl. Table 7. Note that all priors are scaled for one cell, and in the simulations, the results are multiplied by the total number of cells in the sample (i.e., $3 \cdot 10^6$). This decision is based on the fact that each cell faces the concentration of exogenous pyruvate as isolated systems, so the initial conditions should not be scaled to the number of cells, but their reaction product should. We used these priors for the final fine-tuning of the $\mathcal{M}_{CR}(\theta_\gamma)$ model (Fig. 5).

**Suppl. Table 7**: Model $\mathcal{M}_{CR}(\theta_\gamma)$ parameters and initial states for NADH, NAD+ and LDH with their bounds and priors used in this work after estimating the initial conditions experimentally for the final model fine-tuning. Kinetic rates and $y_0$ concentrations are represented for one cell.

| Parameter | Bounds$_\Omega$ | Prior$_\Omega$ |
|---|---|---|
| $k_{in}$ | $[0, 2]$ | $\mathcal{N}(2.5 \cdot 10^{-3}, 2.5 \cdot 10^{-3})$ |
| $k_{PL}$ | $[0, 2]$ | $\mathcal{N}(2.75 \cdot 10^{-2}, 2.75 \cdot 10^{-2})$ |
| $k_b^N$ | $[0, 2]$ | $\mathcal{N}(5 \cdot 10^{-7}, 5 \cdot 10^{-7})$ |
| $k_u^N$ | $[0, 0.3]$ | $\mathcal{N}(1.5 \cdot 10^{-1}, 1.5 \cdot 10^{-1})$ |
| $k_b^P$ | $[5 \cdot 10^{-2}, 5]$ | $\mathcal{N}(9.5 \cdot 10^{-1}, 9.5 \cdot 10^{-1})$ |
| $k_u^P$ | $[0, 2]$ | $\mathcal{N}(9.5 \cdot 10^{-1}, 9.5 \cdot 10^{-1})$ |
| $k_u^{N+}$ | $[0, 2]$ | $\mathcal{N}(1 \cdot 10^{-1}, 1 \cdot 10^{-1})$ |
| $k_b^{N+}$ | $[0, 1 \cdot 10^{-5}]$ | $\mathcal{N}(3 \cdot 10^{-6}, 3 \cdot 10^{-6})$ |
| $k_i$ | $[0, 5]$ | $\mathcal{N}(9.5 \cdot 10^{-1}, 9.5 \cdot 10^{-1})$ |
| $k_a$ | $[0, 2]$ | $\mathcal{N}(9 \cdot 10^{-4}, 9 \cdot 10^{-4})$ |
| $T_1^P$ | - | Computed experimentally (51.23) |
| $T_1^L$ | $[0.92, 80.92]$ | $\mathcal{N}(40.92, 10)$ |
| $ScF$ | - | Computed experimentally per dataset |
| $Pol_L$ | - | Computed experimentally (12%) |
| $[NADH_0]$ | $[45.7, 1409]$ | $\mathcal{N}(457, 238)$ |
| $[NAD{+}_0]$ | $[31.46, 8226.67]$ | $\mathcal{N}(3146, 1270)$ |
| $[LDH_0]$ | $[0, 1.168]$ | $\mathcal{N}(0.3947, 0.193)$ |

To define the parameter priors for the model using experimental data from the LDH activity assay, we first performed a maximum likelihood estimation process to identify viable parameter values for these set of experiments, defining these as $\mu$ and setting $\sigma$ to the same value. The final priors used are shown in Suppl. Table 8.

**Suppl. Table 8**: Model $\mathcal{M}_{CR}(\theta_\gamma)$ parameters and initial states for LDH with their bounds and priors used in this work when inferring parameters using experimental data from the LDH activity assay. Kinetic rates and $y_0$ concentrations are represented for one cell. $y_0$ values are represented in $\mu$M

| Parameter | Bounds$_\psi$ | Prior$_\psi$ |
|---|---|---|
| $k_{PL}$ | $[0, \infty]$ | $\mathcal{N}(2470, 2470)$ |
| $k_b^N$ | $[0, \infty]$ | $\mathcal{N}(5229, 5229)$ |
| $k_u^N$ | $[0, \infty]$ | $\mathcal{N}(152, 152)$ |
| $k_b^P$ | $[0, \infty]$ | $\mathcal{N}(9.5, 9.5)$ |
| $k_u^P$ | $[0, \infty]$ | $\mathcal{N}(915, 915)$ |
| $k_u^{N+}$ | $[0, \infty]$ | $\mathcal{N}(4589, 4589)$ |
| $k_b^{N+}$ | $[0, \infty]$ | $\mathcal{N}(879, 879)$ |
| $k_i$ | $[0, \infty]$ | $\mathcal{N}(11, 11)$ |
| $k_a$ | $[0, \infty]$ | $\mathcal{N}(816, 816)$ |
| $T_1^P$ | - | Computed experimentally (51.23) |
| $T_1^L$ | - | 51.98 |
| $[NADH_0]$ | - | Defined experimentally (220) |
| $[NAD{+}_0]$ | - | Defined experimentally (0) |
| $[LDH_0]$ | $[0, \infty]$ | $\mathcal{N}(0.00268, 0.00268)$ |

#### B Computational analysis exposes our Bayesian OED strategy as the best experimental design approach

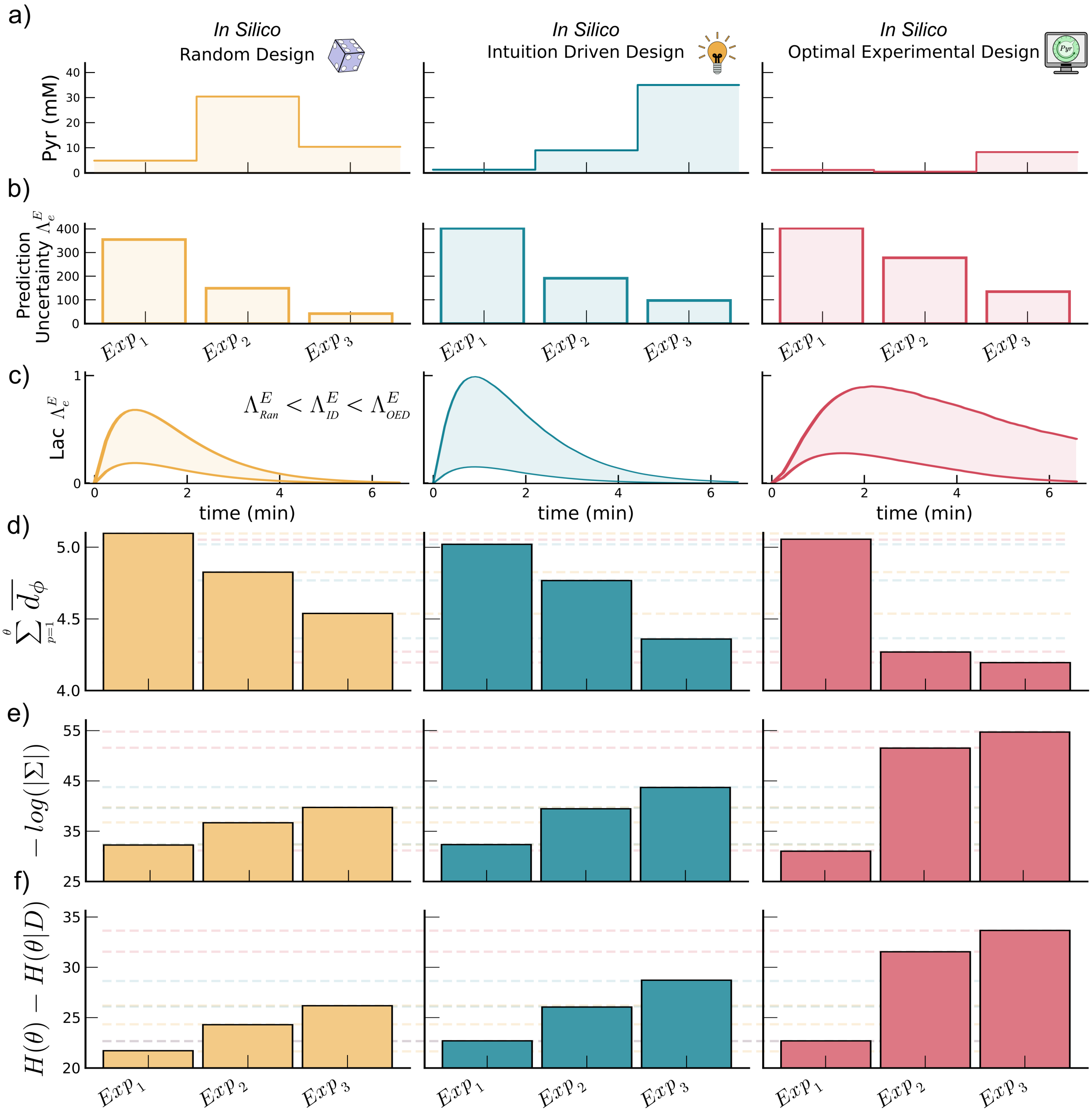


**Suppl. Fig. 1**: Computational validation for a novel Bayesian OED strategy for parameter inference. **a,** Exogenous [1-$^{13}$C]pyruvate concentrations ($mM$) used for three rounds of *in silico* experiments, where we selected the values randomly (yellow), at increasing concentrations by following the average substrate to product curve (blue) or maximising the sample-based prior prediction posterior approximation $\Lambda_e^E$ (red). **b,** $\Lambda_e^E$ value for each pseudo-experiment designed, following the order from panel **a**. **c,** Posterior predictive distributions for the second round of experiments ($Exp_2$) for each design strategy to visualise the uncertainty of the model at that step. Absolute average posterior sample distance $\sum_{p=1}^{\theta} \overline{d_\phi}$ for all model parameters (**d**), negative of the natural logarithm of the determinant of the posterior covariance matrix $-log(|\Sigma|)$ (**e**), and gain in information $G_i = H(\theta) - H(\theta|D)$ (**f**) metrics for each pseudo-experiment designed following the order from panel **a**.

Since parameter inference for model $\mathcal{M}_{CR}(\theta_\gamma)$ poses a complex problem, first, we computationally validated a novel algorithm to design highly informative experiments for the task. For this Bayesian optimal experimental design (bOED) task, we leveraged model prediction uncertainty by seeking to maximise it, to find the experiment our model is the most uncertain about, considering a current prior distribution (Fig.

1e). To do this, we approximate posterior predictive distributions by sampling priors, simulating the model and estimating the uncertainty area of these (e.g., percentile Euclidean distance or entropy of discretised time points). Unlike mutual information approaches, this sample-based algorithm is computationally feasible to execute even for highly complex mathematical models. For example, for models with an elevated number of parameters, the priors could be downsampled to reduce computational times, making it a generalisable and accessible bOED strategy. Yet, as a new bOED approach, we first validated the algorithm using simulated experimental data (i.e., pseudo-data) to estimate its effectiveness in inferring parameters for $\mathcal{M}_{CR}(\theta_\gamma)$. For this quantification, we compared the bOED strategy (Suppl. Fig. 1, red) to traditional experimental design approaches such as random selection of pyruvate concentrations (Suppl. Fig. 1, yellow) or intuition driven ones (Suppl. Fig. 1, blue) where the design followed a pyruvate concentration increase guided by the average pyruvate to lactate curve at each step.

Bayesian optimal experimental design resulted in the leading approach to obtain highly informative data for parameter inference in our computational analysis with pseudo-data. First, we designed three experiments for each design approach selected (Suppl. Fig. 1a). For the bOED strategy, we followed a sequential approach, where we designed each experiment with the last updated posteriors. Except for the first iteration (due to using a highly uninformative prior), bOED achieved the highest prediction uncertainty ($\Lambda_e^E$) compared to the other two designs (Suppl. Fig. 1b) at each step. Suppl. Fig. 1c shows, as an example, the posterior predictive distribution for the second experimental design round to depict the increase in prediction uncertainty. Furthermore, $\Lambda_e^E$ correlated with the gain in information metrics used in all cases (Suppl. Fig. 2). Since we pre-defined the "true" parameter set used to generate the pseudo-data, we can compare the distance of each posterior sample to this vector, resulting in the $\overline{d_{\phi,p}}$ metric. For the first experiment of the round ($Exp_1$), all three design approaches yielded similar results (uninformative priors made any experiment highly informative). In contrast, the other two experiments showed that bOED outperformed the other two strategies (Suppl. Fig. 1d, where lower metrics indicate better performance) with an average $\approx$ 10% increase in each case. The analysis also revealed the most identifiable parameters ($k_{in}$, $k_{PL}$ or $LDH_0$) and the least identifiable ones ($k_b^P$, $k_u^P$ or $k_i$) with this type of experimental data (Suppl. Fig. 3). Information-theoretic metrics corroborated the $\overline{d_{\phi,p}}$ results. For a quick and customary metric to compute (and close to Fisher Information Metric strategies), and assuming posterior normality, we calculated the logarithm of the determinant of the posterior (Suppl. Fig. 1e, where higher values point to better results). The results also highlighted bOED as the best design strategy with a $\approx$ 40% metric increase, with respect to random design, and a $\approx$ 30% for the intuition-driven one. For a more accurate depiction of the gain of information ($G_i$) without assuming normality, we approximated prior and posterior entropies (Suppl. Fig. 1f, where the higher the metric, the better), achieving the same positive results for bOED, with a $\approx$ 30% and $\approx$ 20% metric improvement compared to random or intuition driven designs, respectively. In all cases, the results show that with bOED, two experiments provide more information about the parameters than three experiments in any of the other two approaches. Hence, the results demonstrate the potential effectiveness of bOED to maximise informative content of experiments while minimising time and resources, making it an attractive procedure to introduce in real experimental settings.

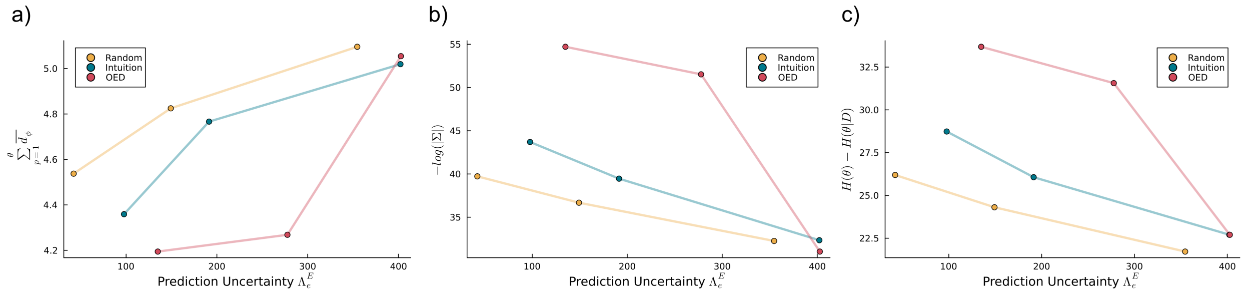


**Suppl. Fig. 2**: Prior prediction uncertainty $\Lambda_e^E$ against posterior uncertainty metrics for computational studies. Scatter plots showing $\Lambda_e^E$ against the average posterior sample distance metric (**a**), the negative logarithm of the posterior covariance matrix (**b**) and the gain in information (**b**) for the computational studies validating the prediction uncertainty Bayesian OED strategy.

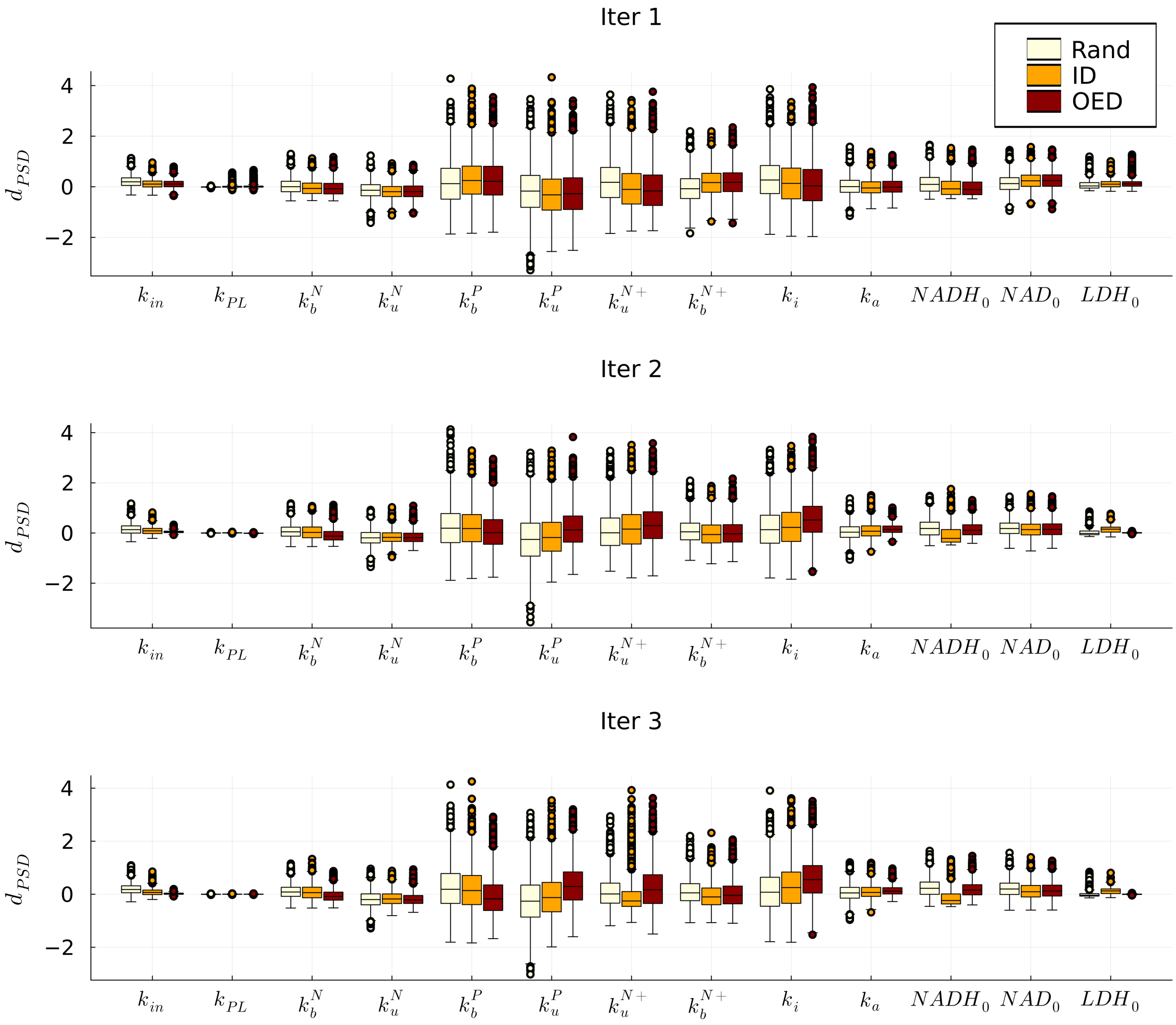


**Suppl. Fig. 3**: Posterior sample distance to true theta for the computational study validating the prediction uncertainty Bayesian OED approach. Box plots showing the posterior sample distance for each round of pseudo-experiments using random (Ran), intuition driven (ID) and Bayesian optimal experimental designs (OED).

### C Dissolution Dynamic Nuclear Polarisation (dDNP)

We used dDNP to increase the polarisation level (p) of $^{13}$C labelled molecules in our samples, as [21]:

$$p = sign(\gamma) \cdot \frac{n_\alpha - n_\beta}{n_\alpha + n_\beta} \tag{5}$$

By applying radio frequency pulses ($\approx$ 94 MHz) at 1.4K in a static 3.35T magnetic field, we increased the population of low energy state nuclei ($n_\alpha$) while decreasing the high energy ones ($n_\beta$). For $^{13}$C nuclei with a positive gyromagnetic ratio $\gamma$=10.71 MHz/T, this produces up to $\approx$ 44.000-fold increase of polarisation compared to thermal equilibrium [22]. In this work, we polarised pyruvate labelled with $^{13}$C at the first position (i.e., [1-$^{13}$C]pyruvate).

##### C.0.1 dDNP pyruvate sample preparation

To prepare our pyruvate samples, first we measured 10.8 mg of trityl OX063 (GE Healthcare) radical in a 1.5 mL eppendorf and 12 $\mu$L of 50 mM Gd-DOTA (Guerbet) in a cut eppendorf lid on a precision balance. Second, we pipetted 408 $\mu$L of 14.63 mM 1-$^{13}$C labelled pyruvic acid (Merck Life Sciences) in the eppendorf and closed the lid to incorporate our Gd-DOTA. Finally, we fully dissolved our solutes by mixing the sample on a vortex for $\approx$5 minutes until no visible crystals remained. We stored all pyruvate stock solutions at -20 °C.

Before each use, we thawed the pyruvate stock sample and mixed it in a vortex for ≈30 seconds to ensure homogeneity. We weighted the sample volume according to our desired final target [1-$^{13}$C]pyruvate concentration ($C_T$), which varied between experiments. For $C_T$ > 20 mM, we weighted 24 $\mu$L of our substrate. When our target concentration reached ≤20 mM and ≤1 mM, we halved (12 $\mu$L) and quartered (6 $\mu$L), respectively, the volume of the used substrate. Note that in this three situations, the pyruvate ejected by the dDNP machine (postdissolution concentration or $C_{PD}$) were 80, 40, 20 mM, respectively. We implemented this three separated pyruvate weights to 1) increase concentration accuracy and 2) ensure that a maximum of 50 % of the final sample volume (i.e., including cells and media) is composed by the dissolution buffer. For dDNP experiments, we use this buffer to: 1) dissolve the frozen polarised substrate [22], 2) reach a final physiological pH of 7.4 in the sample, and 3) adjust the liquid volume for our target [1-$^{13}$C]pyruvate concentration.

##### C.0.2 dDNP pyruvate buffer solution preparation

To make our buffer solution for $C_{PD}$ = 80 mM [1-$^{13}$C]pyruvate samples, we dissolved 4.77 g HEPES, 1.88 g NaOH, 0.05g EDTA-H4 and 0,876 g NaCl in 250 mL MilliQ water. With a magnetic stirrer, we mixed the solution in a volumetric flask until all solutes fully dissolved. We further diluted the solution with MilliQ water for a final volume of 500 mL, thoroughly mixed again, and measured the pH level (12.35 ±0.35). Note that for $C_{PD}$ = 40 and $C_{PD}$ = 20, we halved and quartered the buffer salts while maintaining constant the water volume.

For consistent final [1-$^{13}$C]pyruvate $C_{PD}$ in the dissolved samples, we calculated and used the buffer weight $m_B$ for each experiment according to the [1-$^{13}$C]pyruvate substrate measured weight $m_S$ as:

$$m_B = \left(\frac{m_S}{C_{PD} \cdot M_S} \cdot \rho_B\right) \quad + m_D \tag{6}$$

where $M_S$ is the substrate molecular weight, $\rho_B$ is the buffer solution density and $m_D$ is the buffer mass inside the dead volume of our syringe. We used different syringes for each buffer concentration to avoid cross-contamination and we flushed out the dead liquid volume after each experiment to maintain accurate measurements.

### D Cellular LDH enzyme and MCT1 transporter quantification protocol

We quantified cellular LDH enzyme and MCT1 membrane transporter concentrations of HepG2 cells using flow cytometry. In order to quantify the concentration of the two molecules, we harvested the cells by trypsinisation. For LDH measurement, we washed twice 1 million cells per sample in BD Pharmingen™ Stain Buffer (FBS) containing protein and sodium azide (BD Biosciences, 554656), fixed with BD Cytofix™ (BD Biosciences, 554655) for 30 min at 4 °C and washed using FBS. To permeabilise the fixed cells, we slowly added BD Phosflow™ Perm Buffer III (BD Biosciences, 558050) for 30 min at 4 °C and washed twice using FBS buffer. We resuspended the cells in FBS at $10^7$/ml, stained with Alexa Fluor™ 647 Rabbit Anti-Human LDH antibody (BD Biosciences, 570529) at (9 $\mu$g/mL), and incubated for 30 min at 4 °C. To determine non-specific binding and obtain background levels, we stained the cells with Alexa Fluor™ 647 Rabbit IgG Isotype Control (BD Biosciences, 569345) as an isotype control at same concentration and incubation time. Then, we washed the cells twice with FBS to remove unbound antibodies. Finally we resuspended the cell pellet in 0.5 mL of FBS.

For MCT1 quantification, we washed trice 1 million cells per sample in Flow Cytometry Staining Buffer (FC-SB) containing BSA and sodium azide (R&D systems) and removed the supernatant. We resuspended the cells in FC-SB at $10^7$/ml, stained with Human MCT1/SLC16A1 Alexa Fluor™488-conjugated antibody (R&D systems) at 5 $\mu$g/mL and incubated for 30 min at room temperature. We washed the cells twice with FC-SB to remove unbound antibodies and we resuspended the cells in 0.5 mL of FC-SB. We separately used the isotype Mouse IgG2A Alexa Fluor™ 488-conjugated Antibody at same concentration and incubation time to determine the cell non-specific binding.

We performed the flow cytometry analysis using a Gallios multi-colour flow cytometer instrument (Beckman Coulter, Inc, Fullerton, CA) set up with the 3-lasers 10 colours standard configuration. The machine excited the fluorophore Alexa 647 using a red laser (633 nm) and acquired forward scatter (FS), side scatter (SS) and FL6 (660/20 nm) fluorescence emission and the fluorophore Alexa 488 using a blue laser (488 nm) and acquired FS, SS and FL1 525/BP nm. We excluded aggregates by gating single cells according to their area and peak fluorescence signal.

#### D.1 Calibration curve

We converted the mean fluorescence intensities (MFI) of our cell samples obtained by flow cytometry into number of fluorescent molecules using AccuCheck ERF reference particles (ThermoFisher). We added one drop of particles to a flow cytometry tube and then added 1 mL of FBS. We ran the particles on a flow cytometer experiment, appearing 3 peaks corresponding to 3 different MFI. We created an standard curve by plotting the MFI vs. the Equivalent Reference Fluorophores (ERF) values obtained from the test. Then, we interpolated the results obtained for our samples with the calibration curve to obtain the number of fluorescent molecules $N_{fluor}$.

#### D.2 Conversion from fluorescence units to number of molecules

Once obtaining the number of fluorescent molecules, we quantified the fluorochrome-to-protein (F/P) ratio to determine the number of LDH and MCT1 molecules in our cell sample. First, we calculate the molarity of the antibody (IgG protein) $M$ as:

$$M = \frac{A_{280} - (A_{max} \cdot CF)}{\epsilon} \cdot DF \tag{7}$$

Where $\epsilon$ denotes the protein molar extinction coefficient for IgG (210,000 M $^{-1}$ cm$^{-1}$ according to AAT Bioquest and ThermoFisher).

$A_{280}$ refers to the absorbance used to determine the protein concentration in a sample.

$A_{max}$ corresponds the absorbance of the dye solution at its wavelength maximum for the dye molecule (650 nm for Alexa Fluor 647 and 495 nm for Alexa 488, as reported by AAT Bioquest and ThermoFisher).

Also, correction factor (CF) to adjust the amount of absorbance at 280 nm of the dye (0.03 for Alexa 647 and 0.11 for Alexa 488 as stated by AAT Bioquest) and dilution factor (DF) as dilution of the fluorochrome for absorbance measurement.

Then, the degree of labelling (DOL) was calculated to convert the moles of dye per mole of protein $DOL$:

$$DOL = \frac{A_{max}}{\epsilon' \cdot M} \cdot DF \tag{8}$$

Where $\epsilon'$ is the molar extinction coefficient of the fluorescent dye (270,000 M $^{-1}$ cm$^{-1}$ for Alexa 647 and

73,000 $M^{-1}$ $cm^{-1}$ for Alexa 488, according to AAT Bioquest and ThermoFisher).

Once obtained $N_{fluor}$ and $DOL$, we can calculate the number of IgG molecules $N_{IgG}$ in our samples:

$$N_{IgG} = \frac{N_{fluor}}{DOL} \tag{9}$$

Here, we assume a 1:2 binding ratio between IgG and LDH enzyme. IgG antibodies have two arms, being able to bind two target molecules at same time. Then:

$$N_{enz} = N_{IgG} \cdot 2 \tag{10}$$

#### E Supplementary Figures

This section includes all figures including relevant data for the manuscript that where not included in the main text.

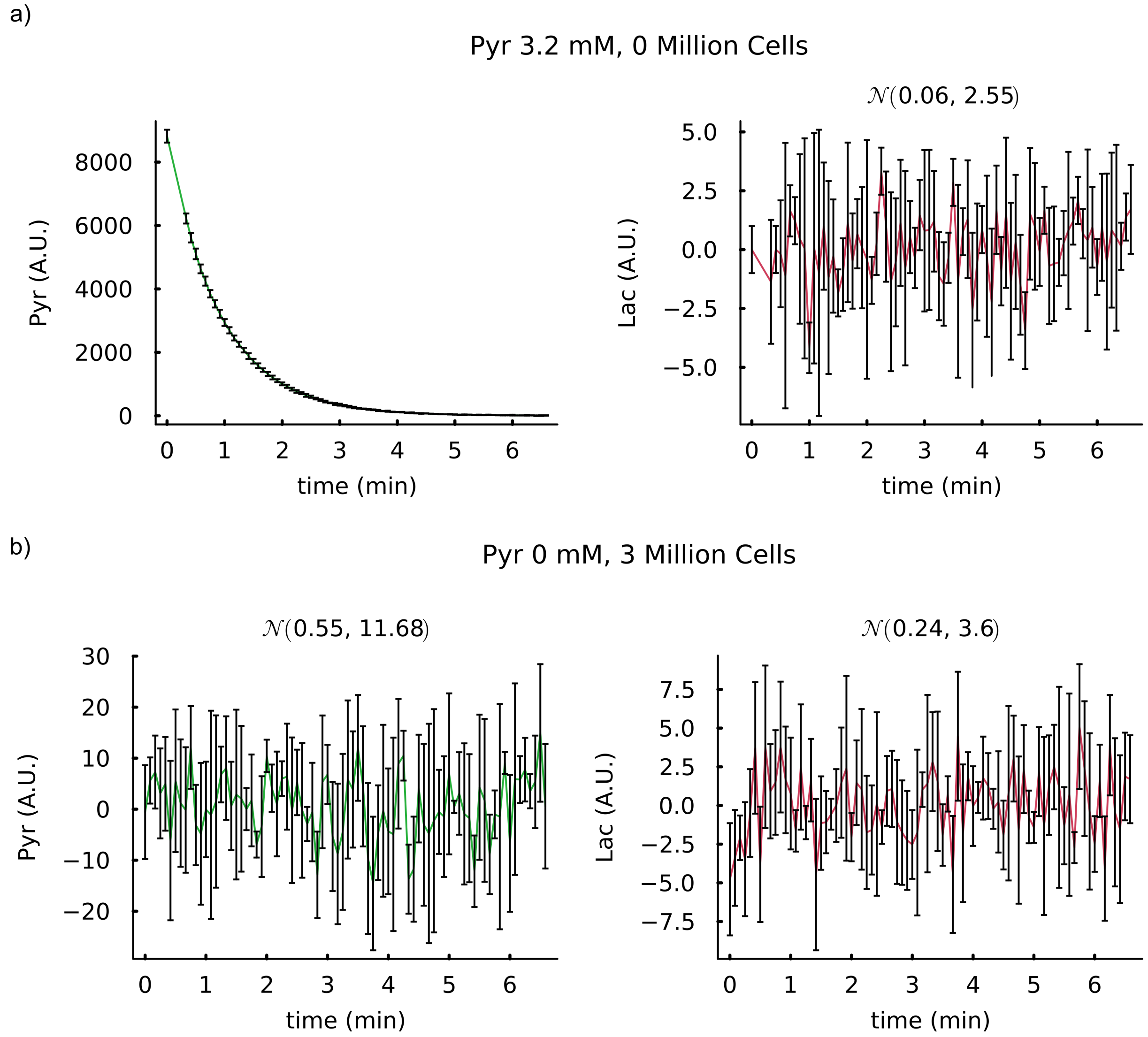


**Suppl. Fig. 4**: NMR control experiments. **a,** NMR experiment including 3.2 mM hyperpolarised [1-$^{13}$C]pyruvate without cells. **b,** NMR experiment using only media without cells or hyperpolarised [1-$^{13}$C]pyruvate.

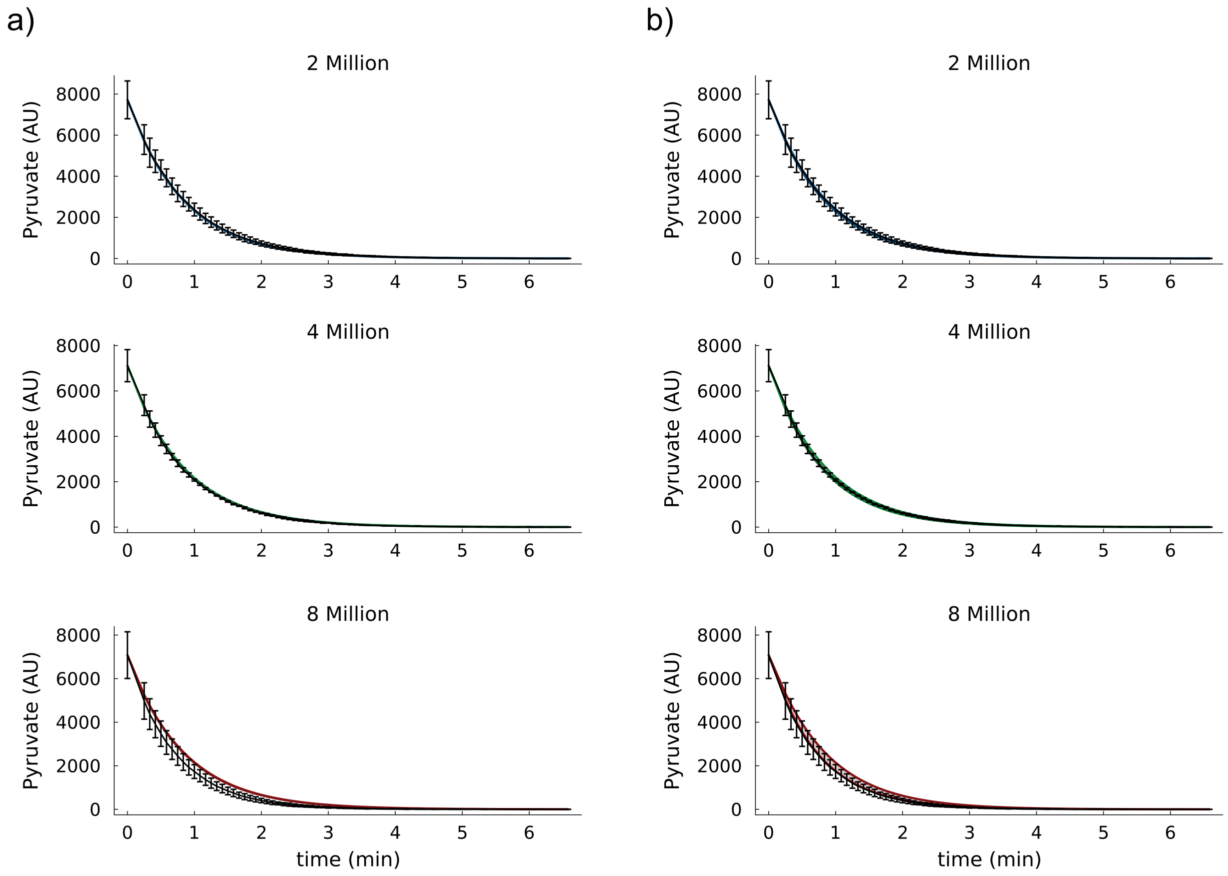


**Suppl. Fig. 5**: Posterior predictions for [1-$^{13}$C]pyruvate signals. Posterior predictive distributions for models $\mathcal{M}(\theta_\alpha)$ (**a**) and $\mathcal{M}_T(\theta_\beta)$ (**b**) for the pyruvate observables of the experiments shown in main Fig. **??** regarding a fix concentration of pyruvate and different number of cells.

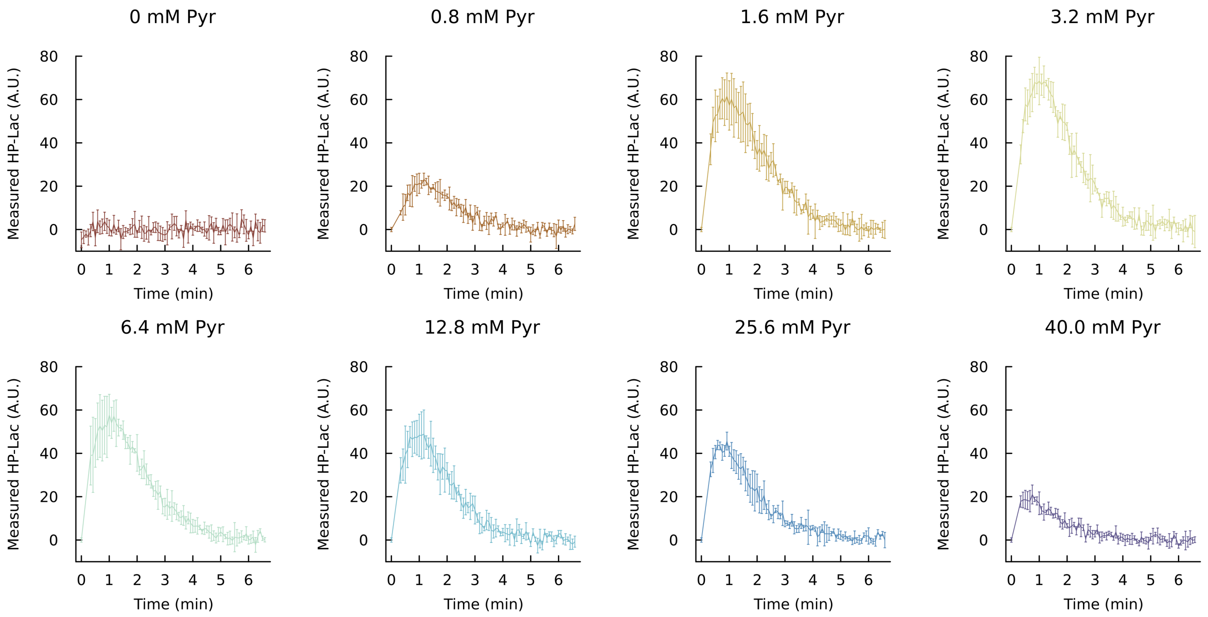


**Suppl. Fig. 6**: Lactate production at increasing pyruvate concentrations. Mean and standard deviation for the observed [1-$^{13}$C]lactate at increasing [1-$^{13}$C]pyruvate concentrations (i.e., 0, 0.8, 1.6, 3.2, 6.4, 12.8 25.6 and 40 mM) for three million HepG2 cells.

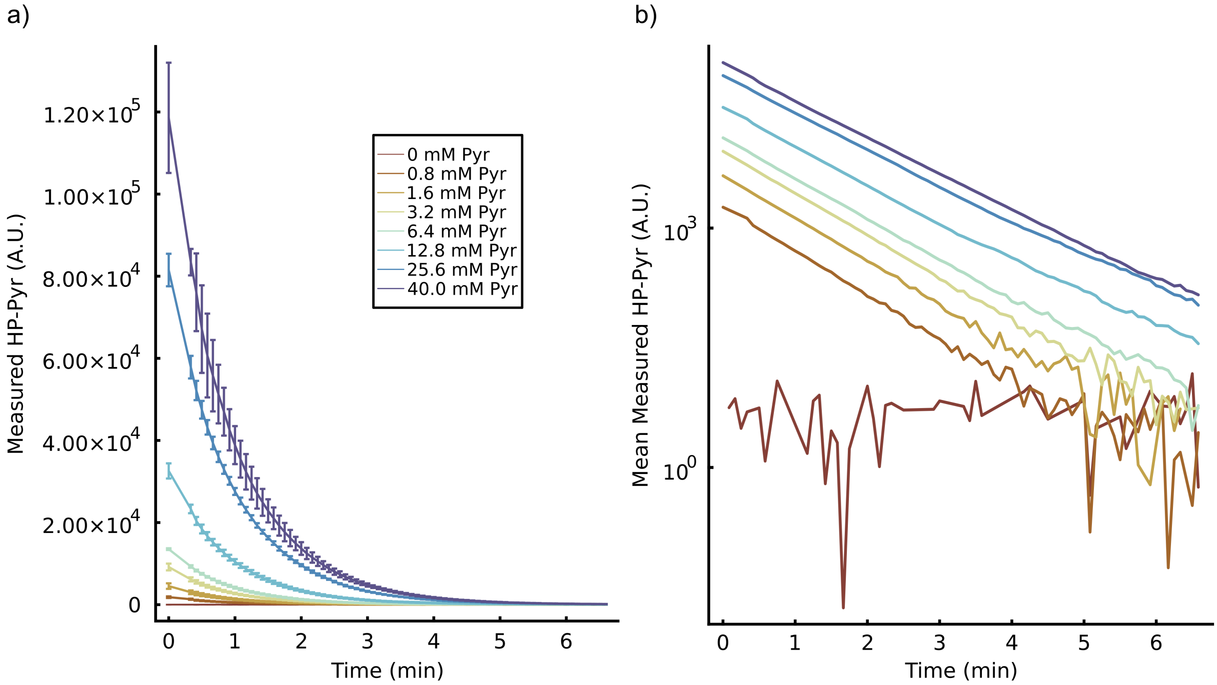


**Suppl. Fig. 7**: Pyruvate experimental data at increasing concentrations. Mean and standard deviation for increasing concentrations of [1-$^{13}$C]pyruvate (i.e., 0, 0.8, 1.6, 3.2, 6.4, 12.8 25.6 and 40 mM) in linear (**a**) and logarithmic (**b**) scales.

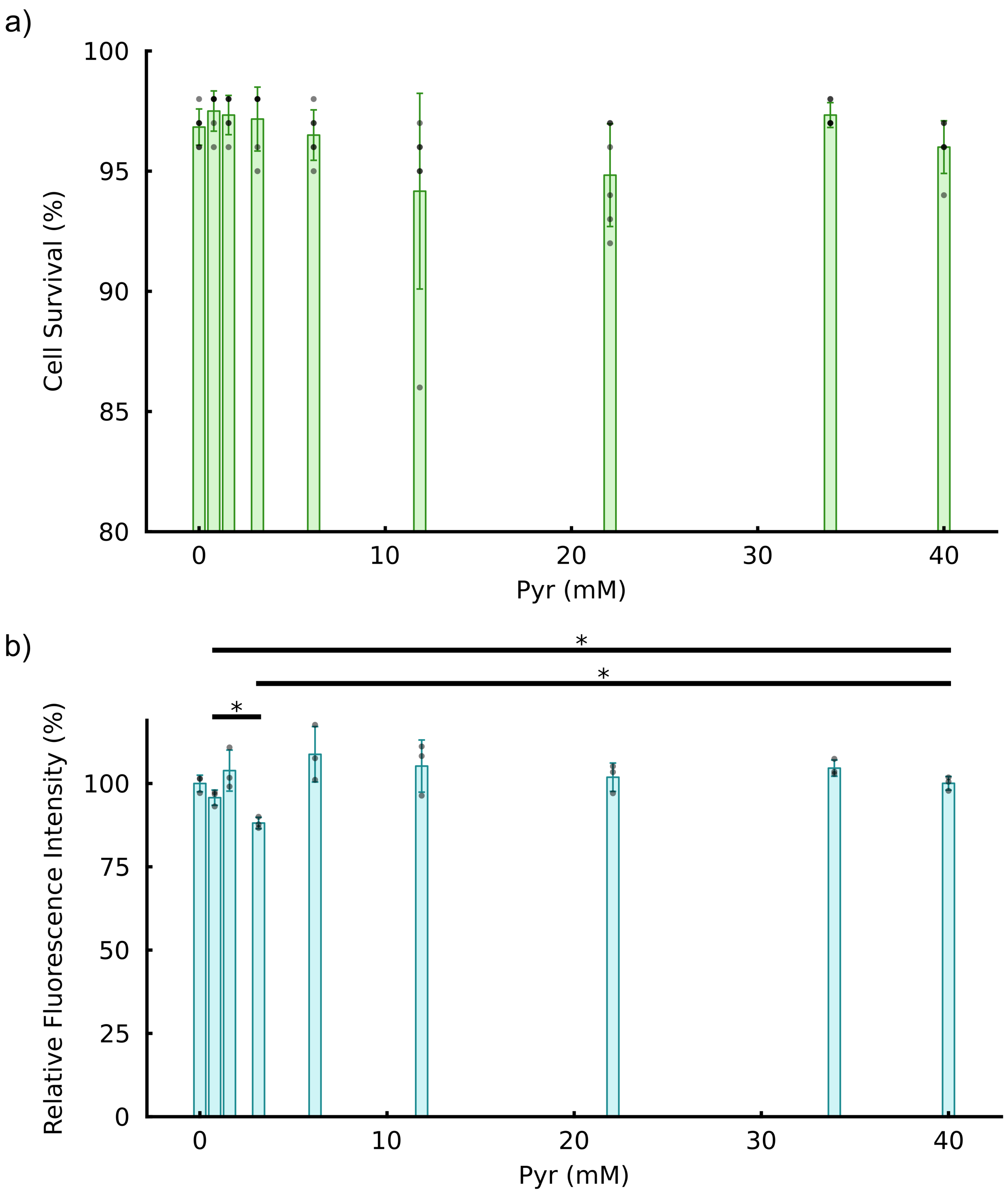


**Suppl. Fig. 8**: Cell viability assays at increasing pyruvate concentrations. **a,** Cell survival (%) at increasing pyruvate concentrations using trypan blue staining. **b,** Cell viability by metabolic alterations measured using alamarBlue HS assay.

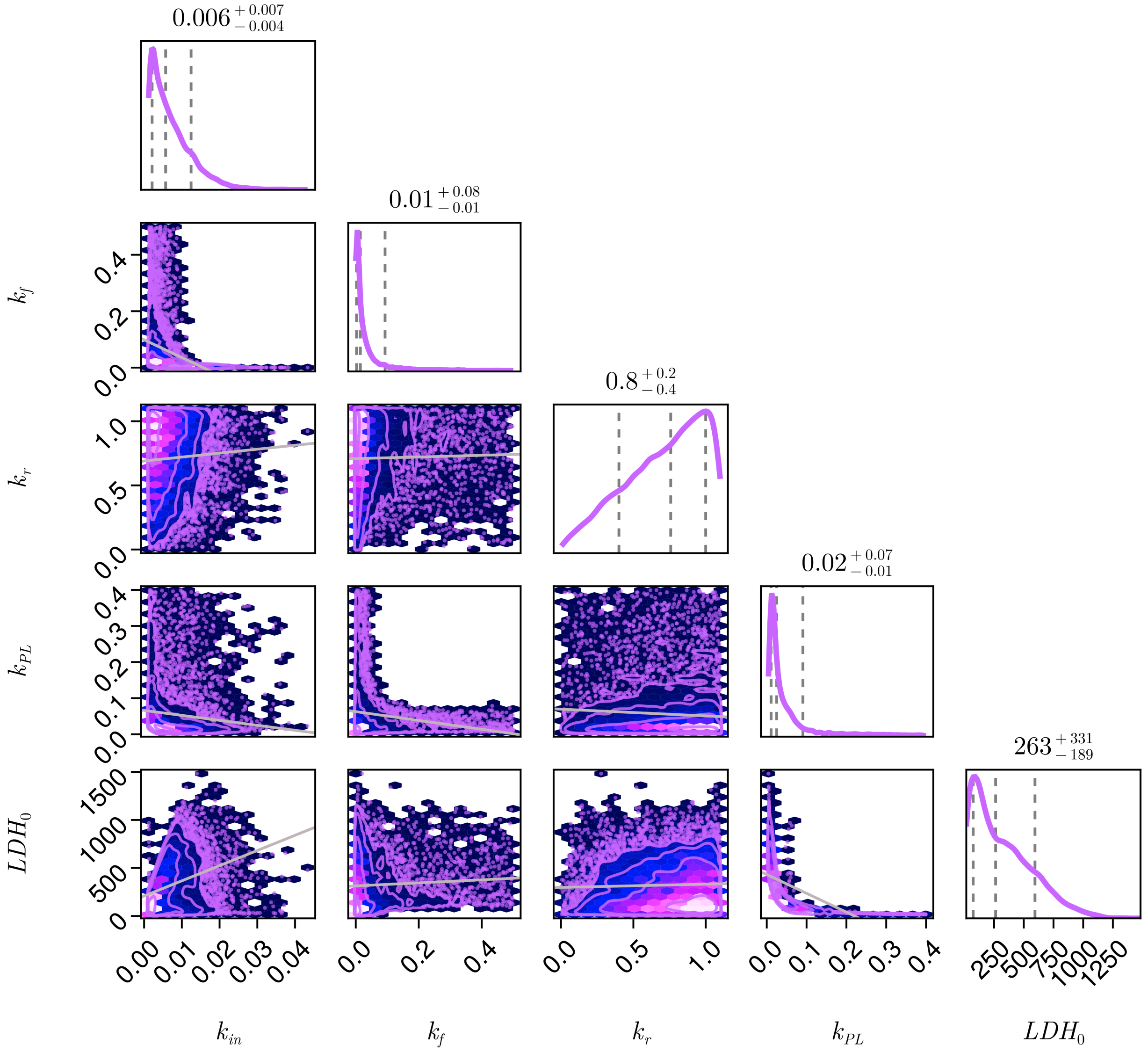


**Suppl. Fig. 9**: Michaelis-Menten model $\mathcal{M}_{MM}(\theta_\delta)$ posterior. Pair plot for all the inferred model parameters ($k_{in}$, $k_f$, $k_r$ and $k_{PL}$) and initial conditions ($[LDH_0]$) for $\mathcal{M}_{MM}(\theta_\delta)$ using an experiment with 3.2 mM [1-$^{13}$C]pyruvate.

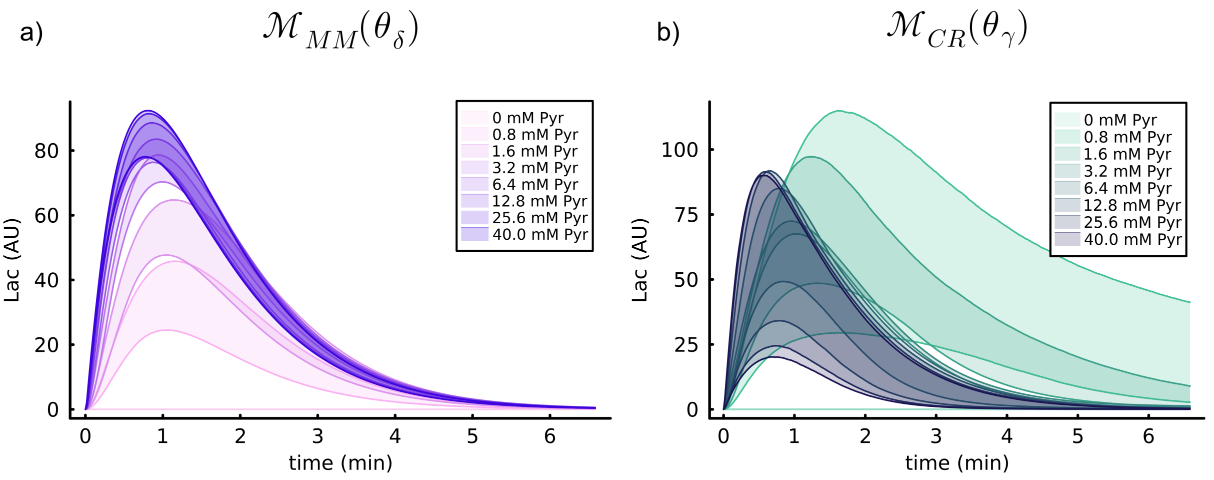


**Suppl. Fig. 10**: Mathematical models $\mathcal{M}_{MM}(\theta_\delta)$ and $\mathcal{M}_{CR}(\theta_\gamma)$ posterior predictive distributions. Simulated [1-$^{13}$C]lactate posterior predictive distributions for $\mathcal{M}_{MM}(\theta_\delta)$ (**a**) and $\mathcal{M}_{CR}(\theta_\gamma)$ (**b**) at increasing [1-$^{13}$C]pyruvate concentrations.

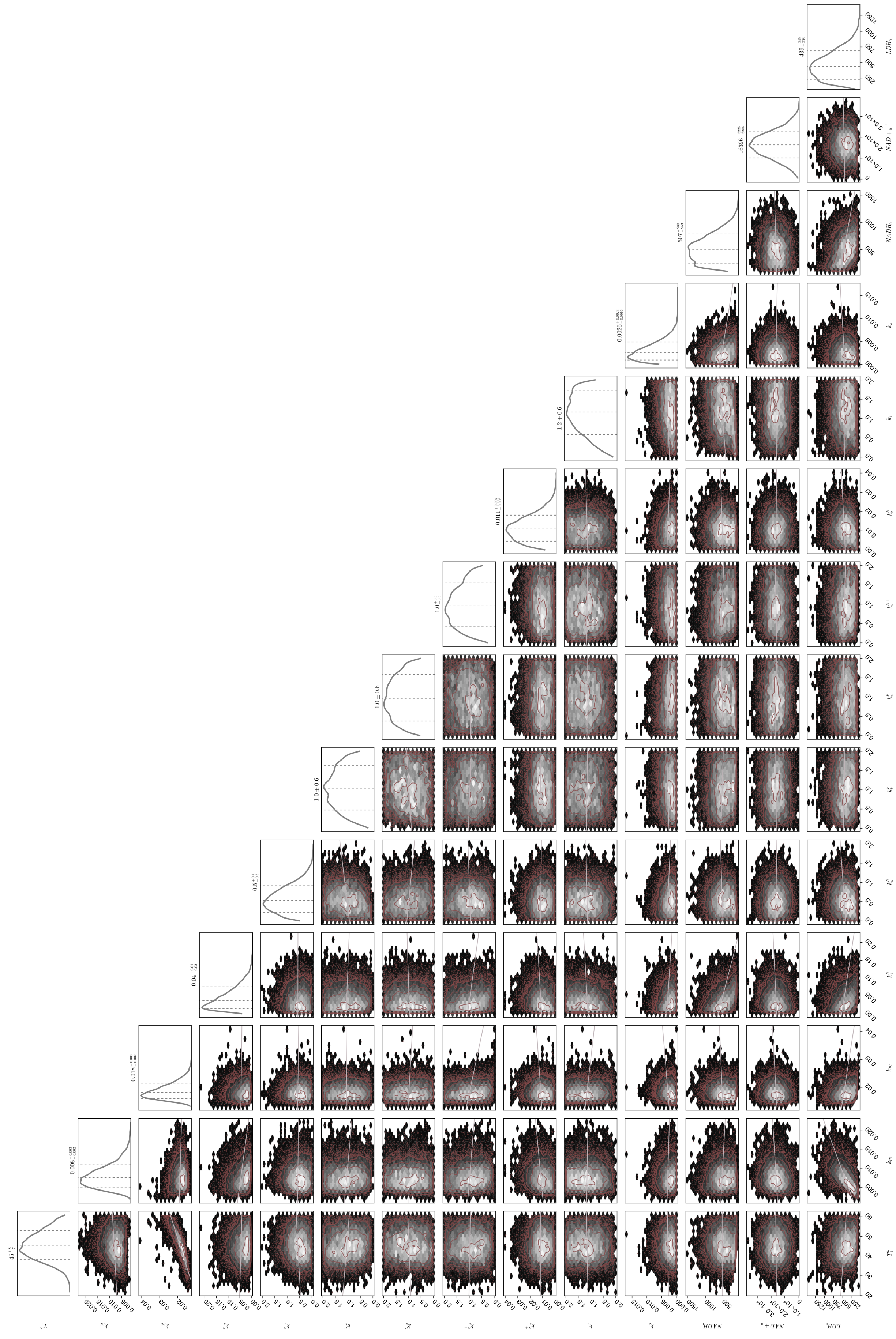

**Suppl. Fig. 11**: Mathematical mode $\mathcal{M}_{CR}(\theta_\gamma)$ posterior distributions after inference with an experiment with 3.2 $mM$ [1-$^{13}$C]pyruvate.

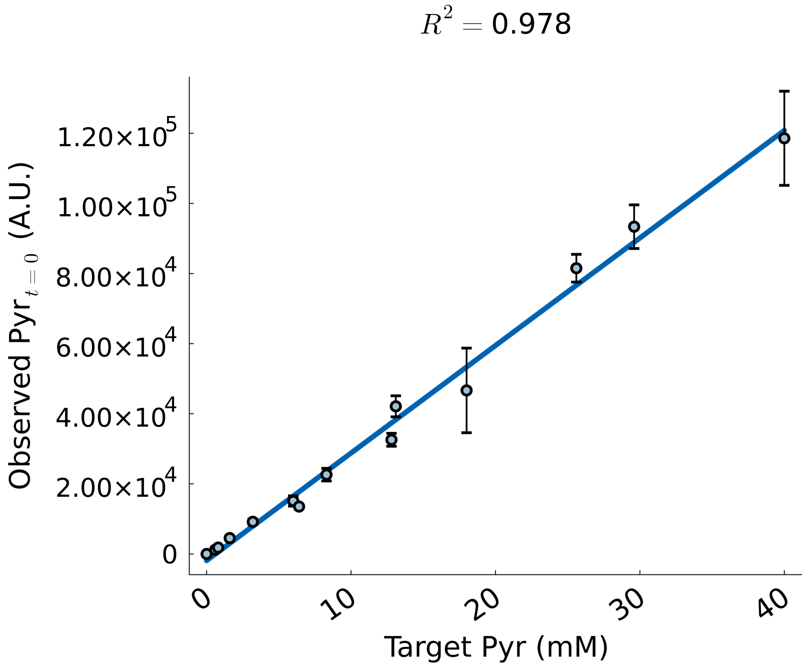


**Suppl. Fig. 12**: Relationship between the target [1-$^{13}$C]pyruvate concentration used in experiments against the observed [1-$^{13}$C]pyruvate NMR signal for all the experiments from this work extrapolated at $t = 0$.

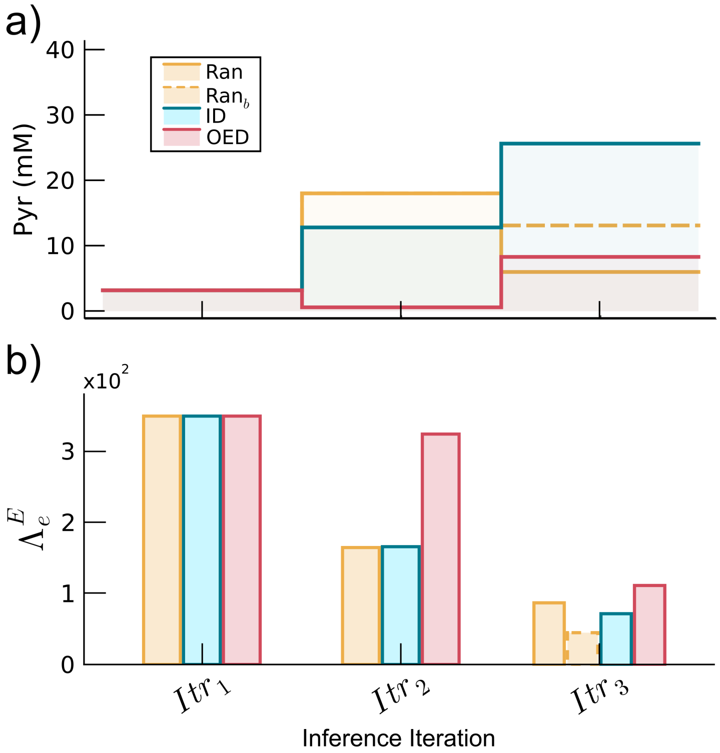


**Suppl. Fig. 13**: Experimental designs for the validation of bOED. [1-$^{13}$C]pyruvate concentrations used for three experimental iterations **a** employing a random (Ran, yellow), intuition driven (ID, blue), or optimal (OED, red) input design and their corresponding prior prediction uncertainty $\Lambda_e^E$ (**b**). The concentrations corresponded to 3.2, 18 and 6 mM for Ran, 3.2, 12.8 and 25.6 mM for ID and 3.2, 0.6 and 8.3 mM for OED.Dashed lines in the third iteration ($Itr_3$) correspond to alternative input designs where, for Ran, it is a substitute random value (13.1 mM) and, for OED, it is a value where, during the optimisation scheme, all predicted signals are normalised by the average maximum of simulations (40 mM).

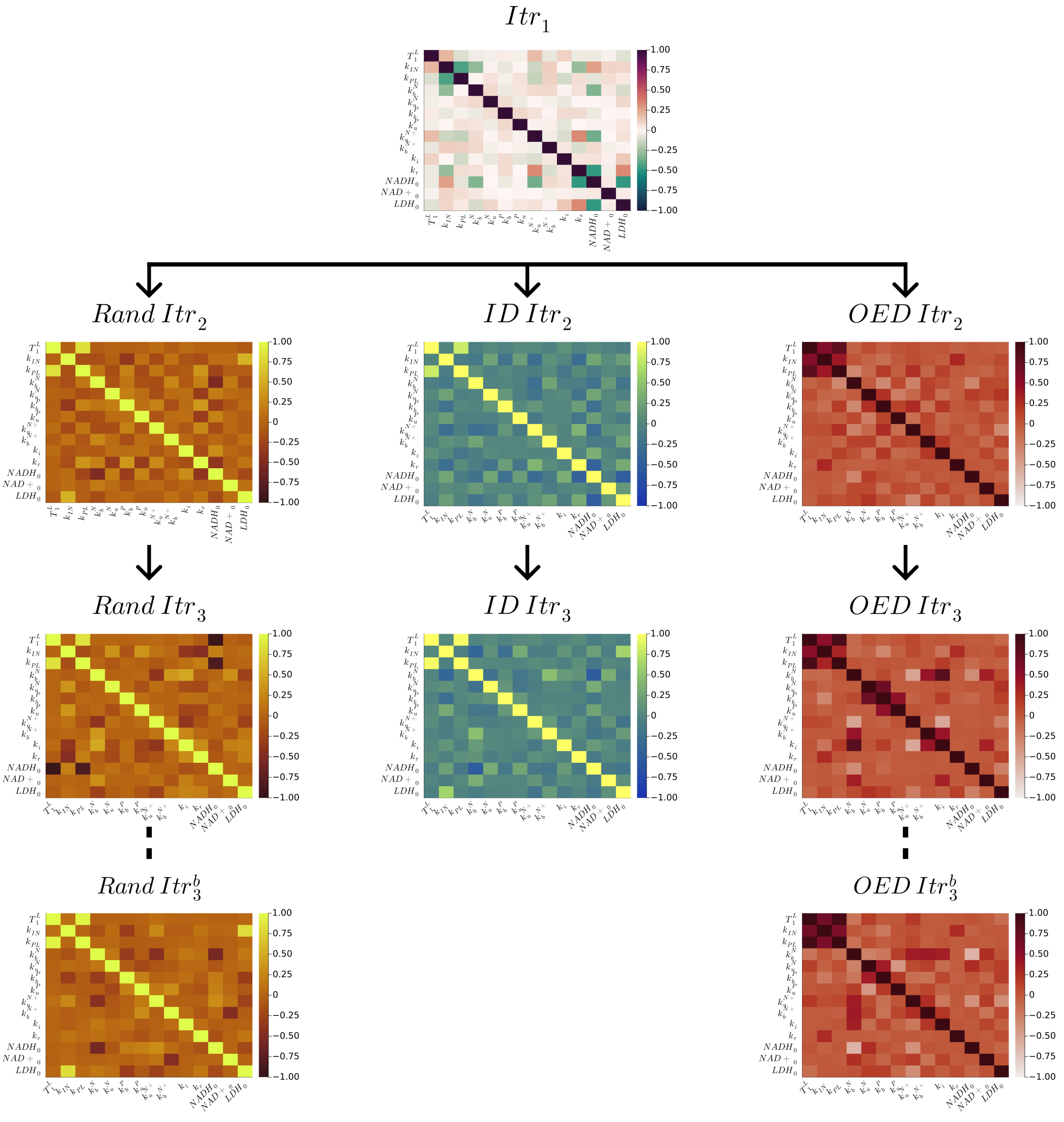


**Suppl. Fig. 14**: Correlation matrices for all the $\mathcal{M}_{CR}(\theta_\gamma)$ posteriors obtained in the *in vivo* validation of the bOED algorithm.

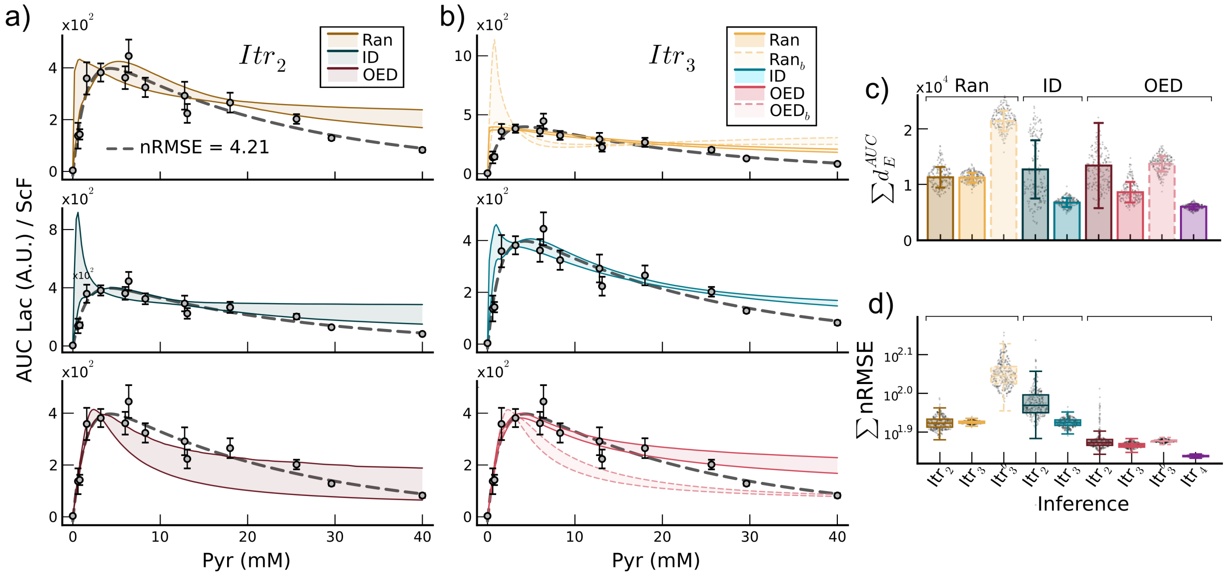


**Suppl. Fig. 15**: Model $\mathcal{M}_{CR}(\theta_\gamma)$ general predictions based on experimental design used to infer parameters. **a,** [1-$^{13}$C]pyruvate to [1-$^{13}$C]lactate relationship prediction for the model $\mathcal{M}_{CR}(\theta_\gamma)$ using all inferred posteriors from random (Ran, yellow, top), intuition driven (ID, blue, middle), and optimal (OED, red, bottom) experiments at the second ($Itr_2$) experimental design iteration. **b,** Same as panel **a**, but for the third ($Itr_3$) experimental design iteration. **c,** Cumulative Euclidean distance $d_E$ between the fitted double exponential substrate-to-product prediction function from panel **a** and the posterior prediction samples for $\mathcal{M}_{CR}(\theta_\gamma)$ (same colour code as panel **a** and **b**) for all the experiments generated in this work. **d,** Cumulative normalised root-mean-square error nRMSE (**d**) for the $\mathcal{M}_{CR}(\theta_\gamma)$ [1-$^{13}$C]lactate posterior prediction samples for all the experiments generated in this work. The parameter posterior distribution used for simulations follows the same colour code as panel **c**.

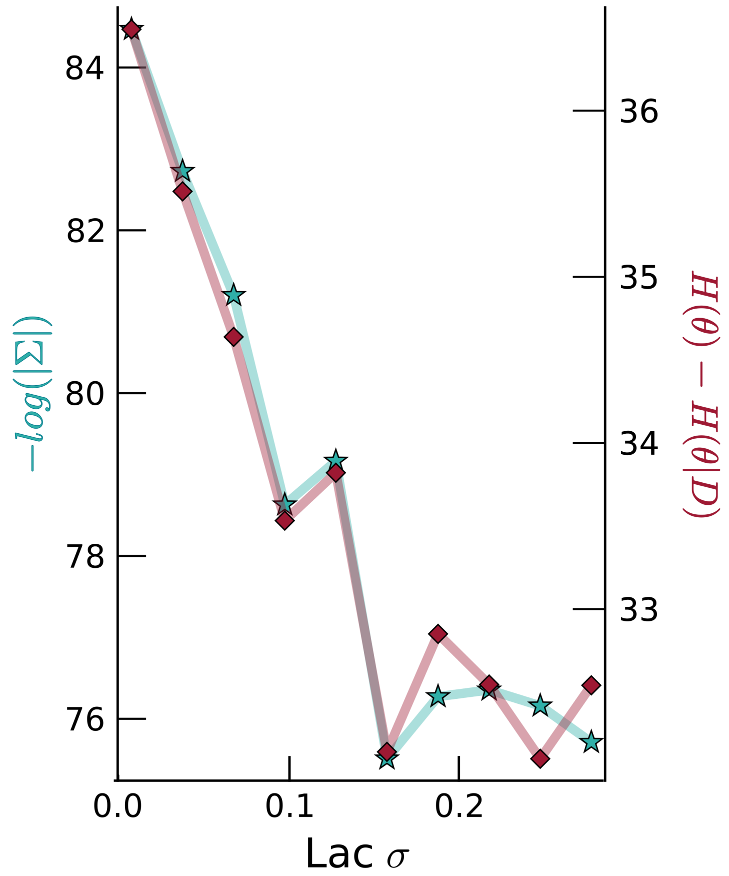


**Suppl. Fig. 16**: Effect of experimental noise in posterior uncertainty. Assessment of posterior uncertainty (using covariance determinant or gain in information) at increasing lactate noise levels in generated pseudo-data.

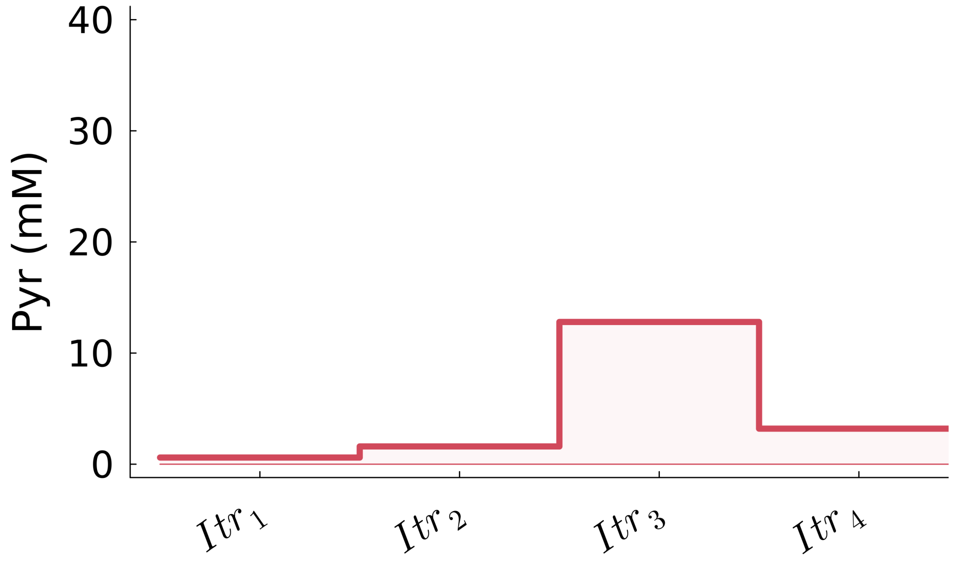


**Suppl. Fig. 17**: Experimental designs for the LDH model $\mathcal{M}_{\mathcal{CR}}(\theta_\gamma)$ fine tuning. [1-$^{13}$C]pyruvate concentrations used for the four experimental iterations (plus alternative forth modulating the objective function, $Itr_4^b$) using OED. The pyruvate concentrations per iteration are 0.6, 1.6, 12.8, 3.2 and 40 mM.

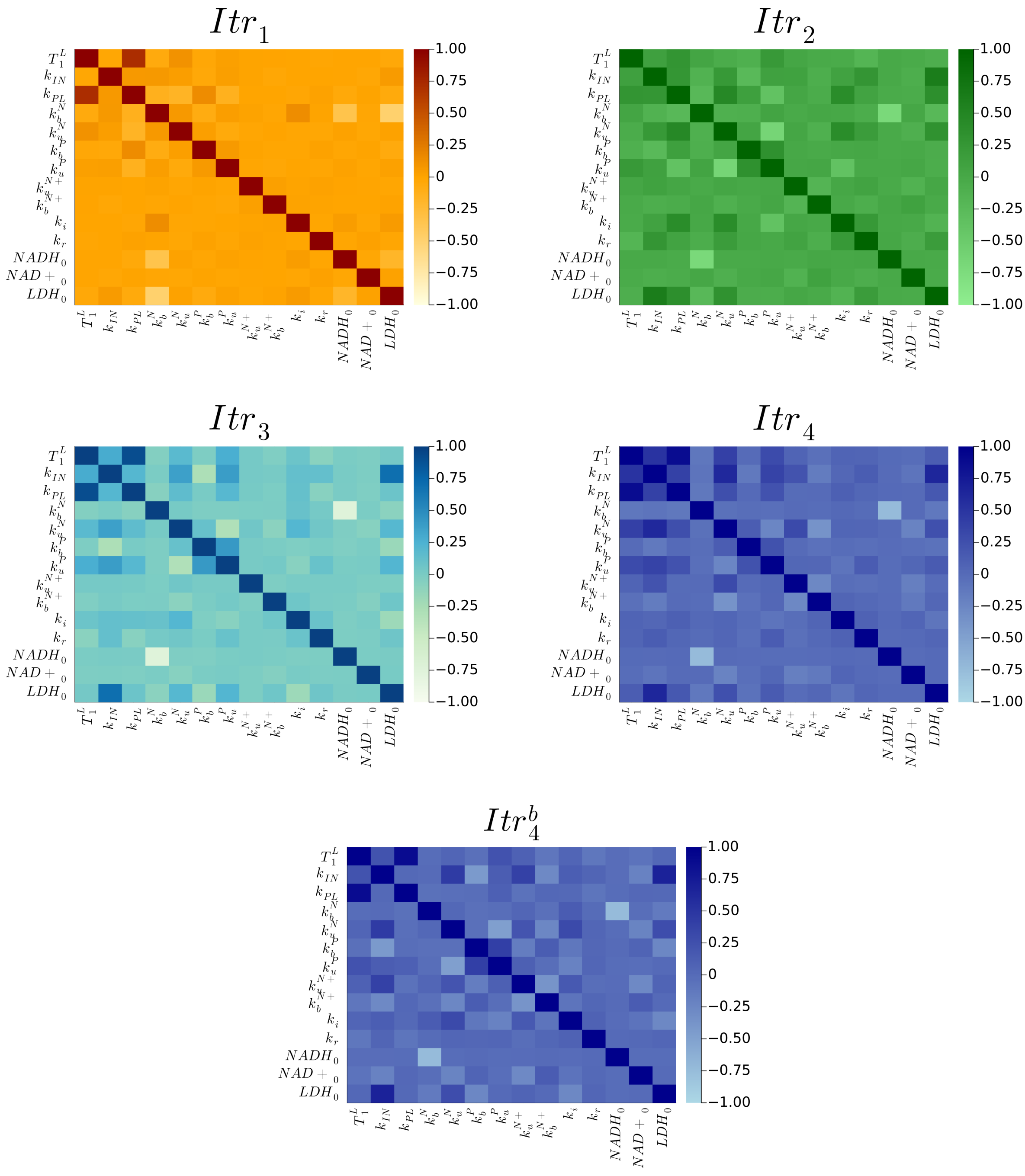


**Suppl. Fig. 18**: Correlation matrices for all the $\mathcal{M}_{CR}(\theta_\gamma)$ posteriors obtained in the model fine tuning for all the inference iterations done.

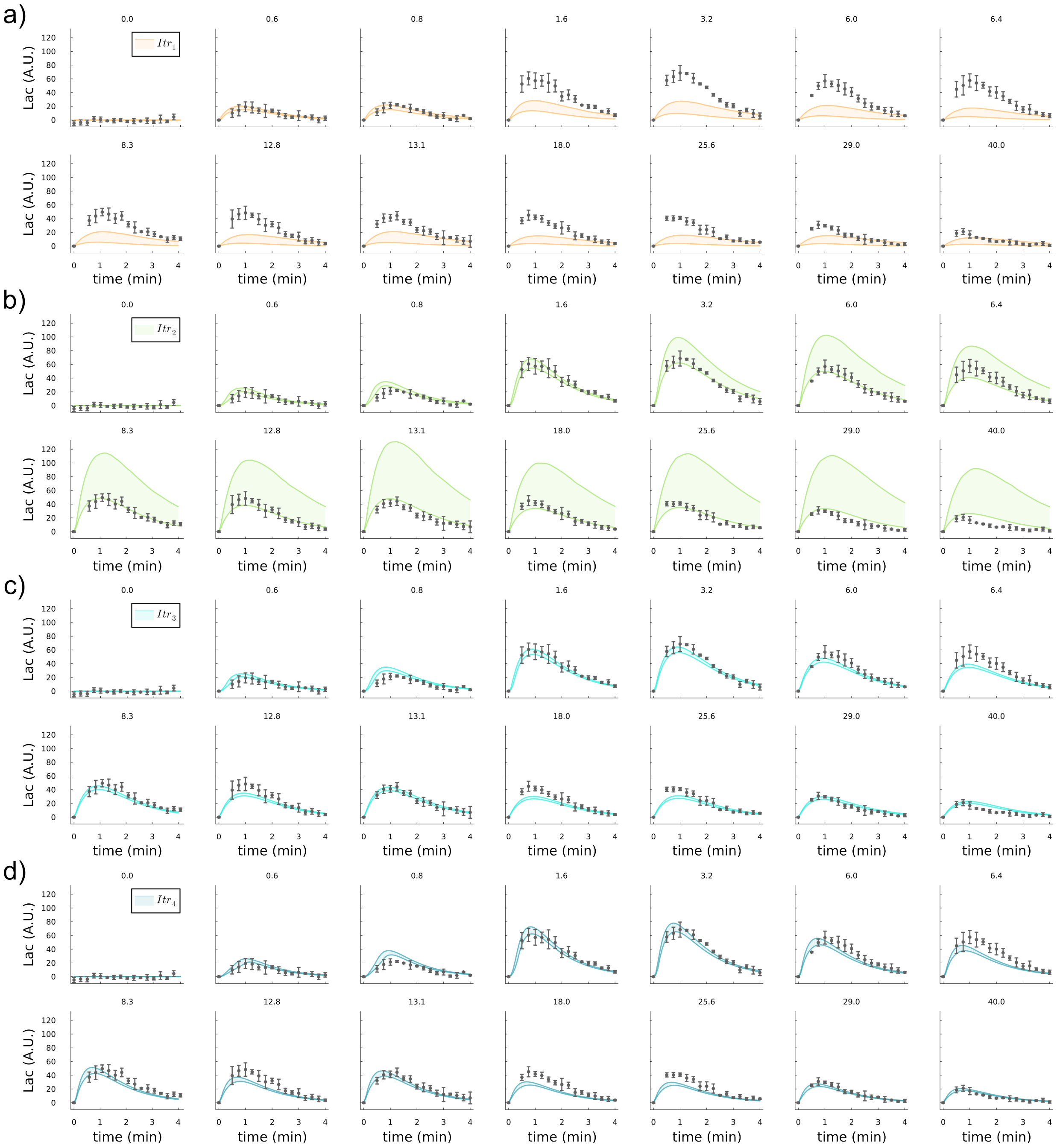


**Suppl. Fig. 19**: Model $\mathcal{M}_{CR}(\theta_\gamma)$ lactate prediction for all experiments in this work, using as parameter posteriors the ones inferred during the fine tuning of the model; $Itr_1$ (**a**), $Itr_2$ (**b**), $Itr_3$ (**c**), $Itr_4$ (**d**).

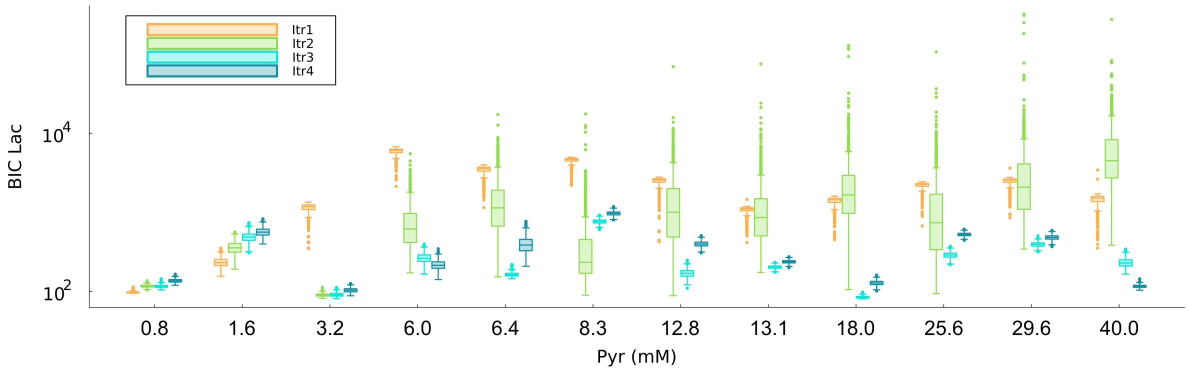


**Suppl. Fig. 20**: Bayesian information criteria (BIC) box plots for all the experiments used in this study predicting them with all the posteriors inferred in the model $\mathcal{M}_{CR}(\theta_\gamma)$ fine tuning. The box plots are generated by using all the inferred parameter samples at each process iteration and then simulating each experiment.

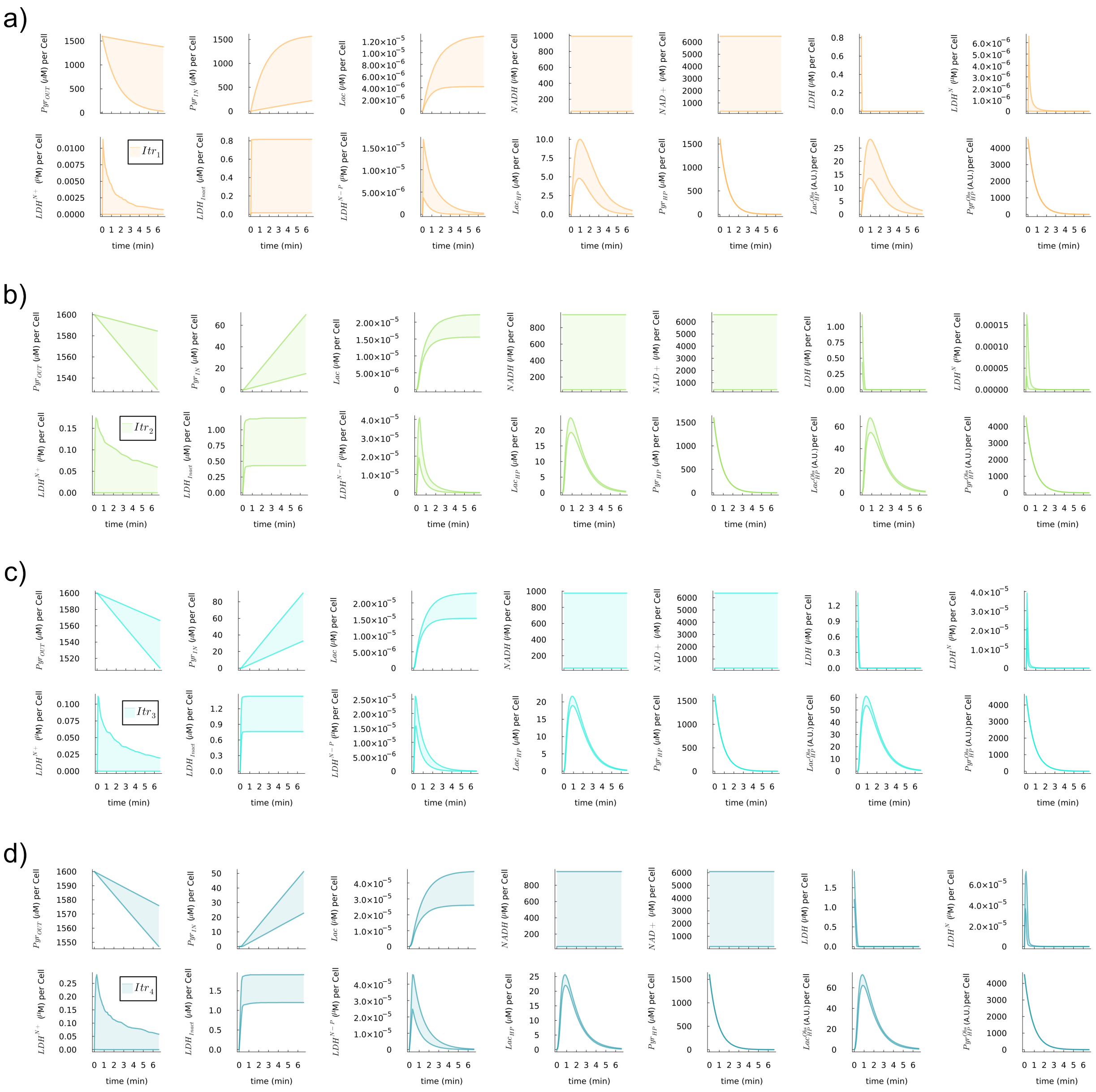


**Suppl. Fig. 21**: Model $\mathcal{M}_{CR}(\theta_\gamma)$ states predictions for an experiment simulating the use of 1.6 mM of exogenous labeled pyruvate, using as parameter posteriors the ones inferred during the fine tuning of the model; $Itr_1$ (**a**), $Itr_2$ (**b**), $Itr_3$ (**c**), $Itr_4$ (**d**). All the biological enzyme states are represented for one cell, while hyperpolarised pyruvate and lactate for $3 \cdot 10^6$ cells. The hyperpolarised pyruvate ($Pyr_{HP}$) and lactate ($Lac_{HP}$), both in $\mu M$, account for a 100 % polarisation, instead of the 12 % of the observables.

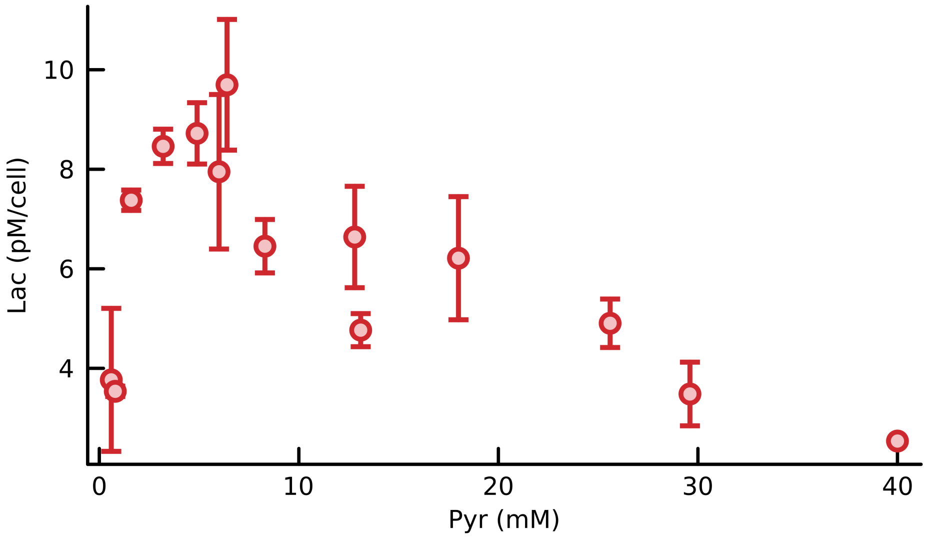


**Suppl. Fig. 22**: Lactate observability in hyperpolarised NMR experiments. Maximum amount of lactate observed by a HepG2 cell at increasing pyruvate concentrations by extrapolating the concentration of the highest lactate $^{13}C$ peak with respect to the highest pyruvate one.

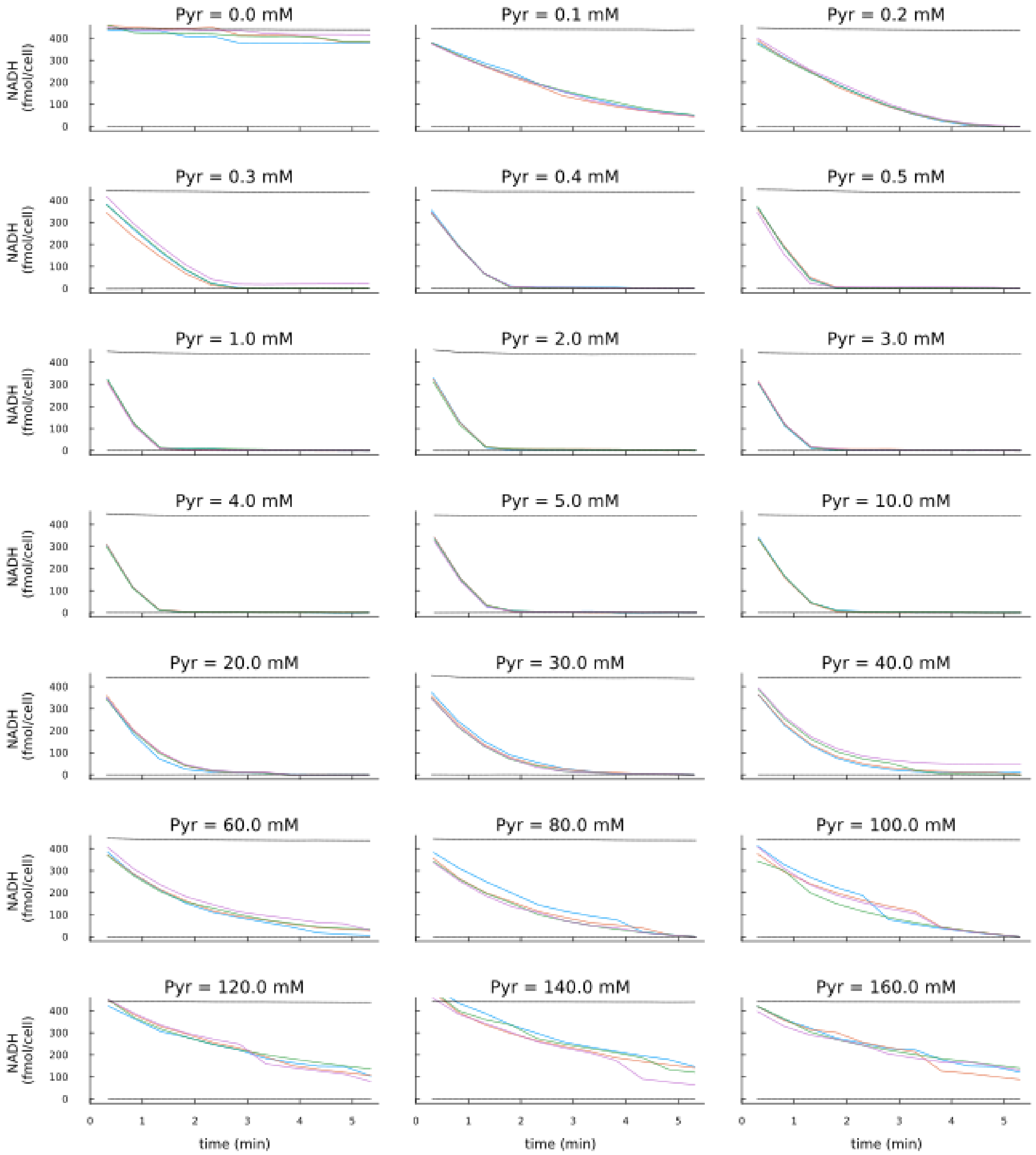


**Suppl. Fig. 23**: LDH enzymatic assay to quantify NADH consumption at increasing pyruvate concentrations. Amount of NADH per HepG2 for 5.5 min of reaction sampled every 30 seconds. On average, we had 20 seconds delay between pyruvate injection in the assay and the first measurement. Dotted black lines represent the positive (top, assay buffer with NADH) and negative (bottom, assay buffer) control values, and colored lines replicates for each experiment.

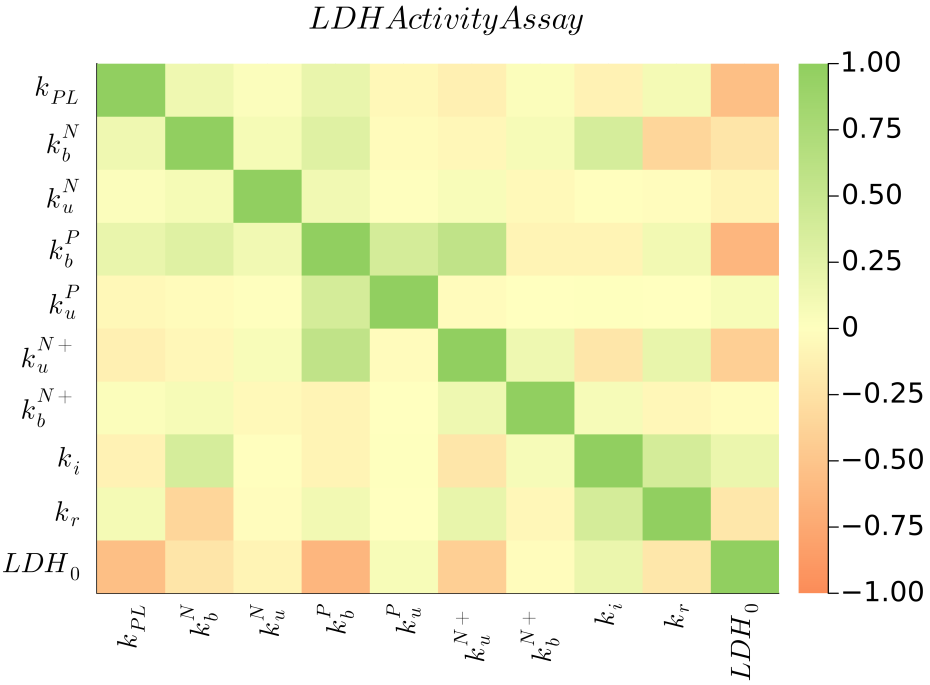


**Suppl. Fig. 24**: Correlation matrix for the $\mathcal{M}_{CR}(\theta_\gamma)$ posteriors without pyruvate membrane transporter obtained in the model inference using all the LDH activity assay data.

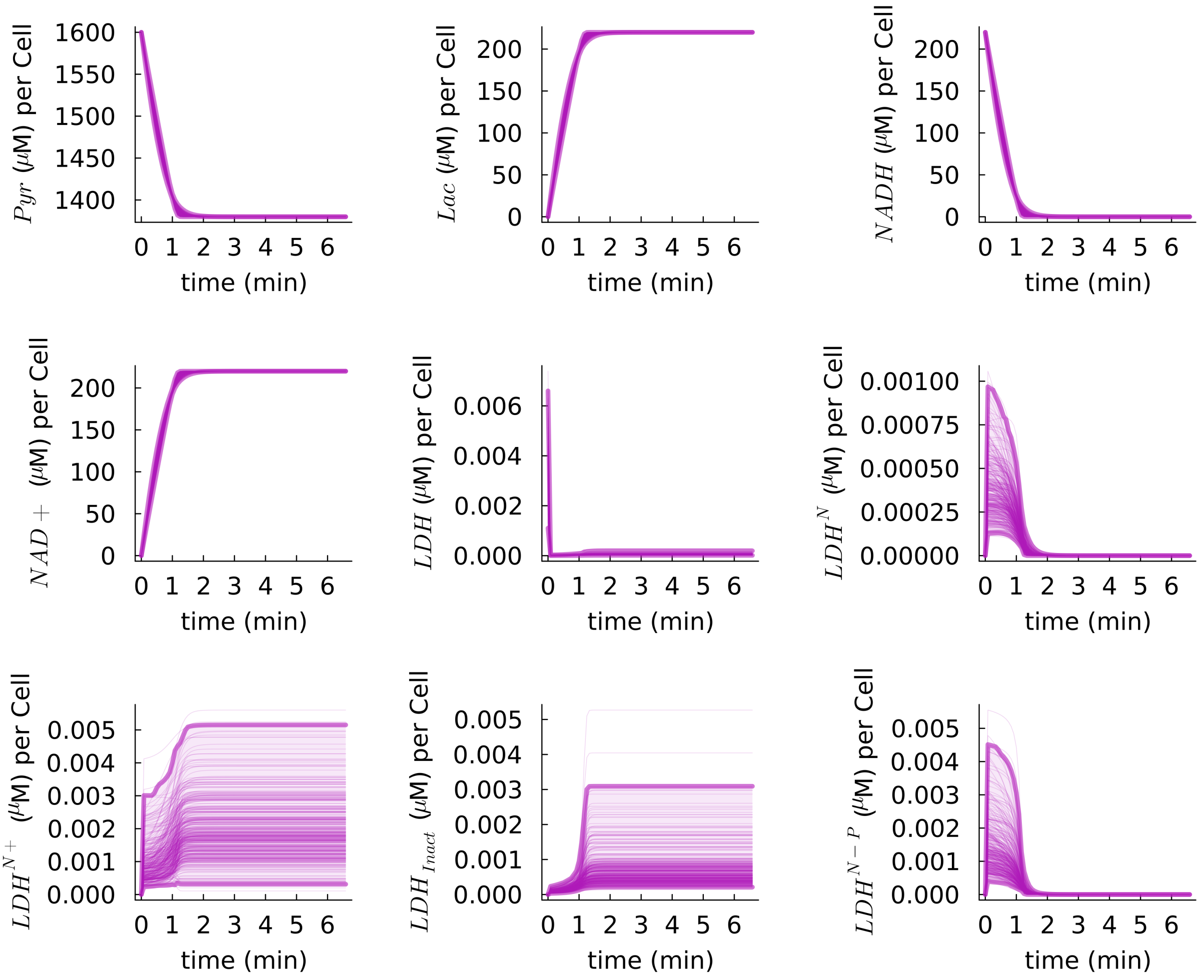


**Suppl. Fig. 25**: $\mathcal{M}_{CR}(\theta_\gamma)$ excluding membrane transport rate $k_{IN}$ system states $y$ posterior predictions for an experiment using 1.6 mM pyruvate, using the $P(\theta|D)$ obtained after inference using LDH activity assay experiments. Note that all concentrations are referenced to the total sample volume of the assay (200 $\mu$L).
